# Signature activities of 20S proteasome include degradation of the ubiquitin-tag with the protein under hypoxia

**DOI:** 10.1101/2019.12.20.883942

**Authors:** Indrajit Sahu, Sachitanand M. Mali, Prasad Sulkshane, Andrey Rozenberg, Cong Xu, Roni Morag, Manisha Priyadarsini Sahoo, Sumeet K. Singh, Zhanyu Ding, Yifan Wang, Sharleen Day, Yao Cong, Oded Kleifeld, Ashraf Brik, Michael H. Glickman

## Abstract

Careful removal of unwanted proteins is necessary for cell survival. The primary constitutive intracellular protease is the 26S proteasome complex, often found in equilibrium with its free catalytic subcomplex– the 20S core particle. Protein degradation by 26S is tightly regulated by prior ubiquitination of substrates, whereas 20S is amenable to substrates with an unstructured segment. Differentiating their contributions to intracellular proteolysis is challenging due to their common catalytic sites. Here, by chemically synthesizing a synoptic set of homogenous ubiquitinated proteins, we ascribe signature features to 20S function and demonstrate a unique property: degrading the ubiquitin-tag along with the target protein. Cryo-EM confirms that a ubiquitinated substrate can induce asymmetric conformational changes to 20S. Mass-spectrometry of intracellular peptidome under hypoxia and in human failing heart identifies the signature properties of 20S in cells. Moreover, the ability of 20S proteasome to clear toxic proteins rapidly, contributes to better survival under these conditions.

## Introduction

All eukaryotic cells continuously recycle proteins for survival; these include damaged proteins as well as retired proteins. The ubiquitin proteasome pathway is the major proteolytic system for cytosolic and nuclear proteins. Selection of target substrates by covalently tagging them with ubiquitin is the initial phase of this process. The resulting ubiquitin-conjugates are delivered to the 26S proteasome complex for proteolysis (Finley, 2009; Glickman and Ciechanover, 2002). Within the 26S holoenzyme, the 19S regulatory particle (RP) is responsible for recognizing the ubiquitin signal and unfolding the target protein, whereas the 20S core particle (CP) hydrolyses the unfolded polypeptide (Orlowski and Wilk, 2003; Saeki, 2017; Yu and Matouschek, 2017). To do so, the 19S RP utilizes three ubiquitin receptors (Rpn1/PSMD2, Rpn10/PSMD4 and Rpn13/ADRM1 subunits) and an AAA hexameric ring (RPT1-6 subunits). This 19S RP attaches to either side of the 20S barrel, defined by heptameric α-rings (α1-7 subunits). Substrates apparently traverse through a channel in the center of the α-ring on their way from the 19S RP to the proteolytic β-active sites (β1, β2 and β5 subunits) located within the central cavity of the barrel-shaped 20S CP (Groll et al., 2000; Groll et al., 1997; Unno et al., 2002).

Although the 20S subcomplex is an integral part of the 26S holoenzyme, it is quite abundant as a free complex in many cell types (Fabre et al., 2014). It has been suggested that free 20S may be a proteasome assembly intermediate, a 26S breakdown product (due to disassembly), or a stand-alone proteolytic enzyme (Bajorek et al., 2003; Demasi and da Cunha, 2018; Hendil et al., 2009; Hohn and Grune, 2014; Jung and Grune, 2008; Kumar Deshmukh et al., 2019; Livnat-Levanon et al., 2014l; Njomen et al., 2018; Pickering and Davies, 2012; Raynes et al., 2016; Sahara et al., 2014; Tomko and Hochstrasser, 2013; Tsvetkov et al., 2015). For instance, prokaryotes, which lack ubiquitin, do have 20S complexes and other ATP-dependent proteases, supporting the notion of 20S being the primordial protein-degrading machine (Majumder and Baumeister, 2019). However, without an associated unfoldase activity, free 20S CP *in vitro* proteolyse only unstructured proteins in a ubiquitin-independent manner (Glickman et al., 1998; Shringarpure et al., 2003). In eukaryotic cells, the 20S is activated by attachment to 19S RP, may be partly augmented by other non-ATPase activators such as PA28 or PA200 (Baldin et al., 2008; Fabre et al., 2014; Suzuki et al., 2009) although, it may also function independently and act directly on disordered proteins (Arrigo et al., 1988; Chondrogianni et al., 2015; Myers et al., 2018; Shringarpure et al., 2001; Suskiewicz et al., 2011) or oxidized/damaged proteins (Chondrogianni et al., 2015; Davies, 2001; Pickering and Davies, 2012; Raynes et al., 2016; Tonoki et al., 2009). Attachment of proteasome activators not only influences substrate selection but may also affect product outcome due to allosteric effects on β-catalytic active sites (Emmerich et al., 2000; Kisselev et al., 1999). Since the ratio of 20S to 26S proteasome varies across different cellular conditions, a dynamic equilibrium between the two species may be part of an adaptive response to cellular needs (Mayor et al., 2016).

Any physiological condition that demands an alteration to the proteome or impairs protein function requires enhanced capacity to remove the unnecessary load. Common stress conditions such as oxidation, temperature, ionization or toxins damage proteins, but also inevitably affect the ubiquitin-proteasome machinery. Proteasome impairment can lead to 26S accumulation in storage granules (Enenkel, 2018; Marshall and Vierstra, 2018; Peters et al., 2013), its disassembly (Bajorek et al., 2003; Livnat-Levanon et al., 2014; Wang et al., 2010), ubiquitination (Besche et al., 2014), or to proteophagy (Cohen-Kaplan et al., 2016; Marshall et al., 2015; Wen and Klionsky, 2016). Interestingly, 20S CP is relatively resistant to oxidation damage compared to 26S and persists as a stable complex under such conditions (Hohn and Grune, 2014; Reinheckel et al., 1998). Hence, most likely 20S plays a role under stress conditions, but how it serves to alleviate proteotoxicity is unclear.

An age-related decline of 26S proteasome function correlates with an inefficiency to remove damaged and aggregate-prone proteins (Baraibar and Friguet, 2012; Breusing and Grune, 2008; Bulteau et al., 2002; Saez and Vilchez, 2014; Tonoki et al., 2009). One explanation is that mitochondrial dysfunction leads to reactive oxygen species (ROS), which causes pervasive oxidative damage to proteins – including the ubiquitin proteasome machinery – thereby increasing the intracellular proteotoxic load (Bulteau et al., 2006; Cadenas and Davies, 2000). Inefficiency of mitochondria to respire at low oxygen (hypoxia) is a physiological condition that promotes oxidative damage to proteins (Guzy and Schumacker, 2006; Solaini et al., 2010). For instance, ischemic-related hypoxia, a pathological condition for failing heart, is characterized by oxidative stress (Okonko and Shah, 2015), and disassembly of 26S proteasome (Day et al., 2013; Predmore et al., 2010). Fine-tuning the proteolytic machinery may be a strategy, which cells utilize to survive under hypoxia. While many studies have focused specifically on the decline of 26S proteasome in acute conditions such in failing heart and during aging in general, we proposed to look at the residual activity of 20S proteasome under hypoxia.

Since the two proteasomes (26S, 20S) have the same active sites, deconvoluting the distinct contribution of 20S to bulk intracellular protein degradation has been challenging (Raynes et al., 2016). A precise and systematic approach is needed to establish a distinctive proteolytic function of 20S proteasome separate from 26S proteasome *in vivo*. Our ability to chemically synthesize homogenous polyubiquitinated proteins (Singh et al., 2016; Sun and Brik, 2019), visualize substrate interaction with proteasome (Ding et al., 2019), and identify MS/MS products of intracellular proteolysis (Huang et al., 2018; Kleifeld et al., 2011; Singh et al., 2016), enabled us to distinguish between 20S and 26S function, and use their signature properties to resolve their contributions in a cellular context.

## Results

### Hypoxia induced proteasome disassembly facilitates proteotoxic clearance

The proteasome populations respond to the cellular environment in order to maintain proteostasis. For instance, altering the redox potential alters the ratio between 26S and 20S proteasomes; specifically, 26S disassembles into stable 20S and 19S sub-complexes upon oxidative stress whether externally or internally induced (Livnat-Levanon et al., 2014). A rapid burst of reactive oxygen species (ROS) is generated in cells when expose to hypoxia (Fuhrmann and Brune, 2017). Therefore, we measured whether proteasome populations respond to hypoxia. HeLa cells growing under severe hypoxia (1% O_2_) showed a time-dependent disassembly of 30S and 26S proteasomes to 20S and 19S sub-complexes (**Figure 1A**) without affecting total proteasome content (**Figure 1C**). Within 36 hours of hypoxic growth, proteasome population shifted from ∼80% 30S/26S to ∼80% 20S (**Figure 1B**). This result indicates that it is possible to obtain cells with a high 20S proteasome content at 24 hours of hypoxia (**Figure 1D**), which intrigued us to measure protein turnover under this condition. In order to increase the flux of proteasome substrates we exploited puromycin-induced proteotoxicity (Yau et al., 2017) and quantified their clearance during a recovery period. Puromycin specifically incorporates at the C-terminus of nascent polypeptides and can easily be traced with an anti-puromycin antibody (**Figure S1A**). We observed that these nascent truncated proteins were cleared from hypoxic cells faster than from cells growing under normoxia (**Figure 1E and 1F**). The fast removal of puromycin-conjugates under hypoxia can be attributed to proteasome function than lysosome activity (**Figure S1B and S1C**). Nevertheless, proteolysis was largely unaffected by the E1 inhibitor (PYR41), suggesting that large portion of puromycin-conjugates followed ubiquitin-independent degradation pathway (**Figure S1C**).

**Figure 1.**
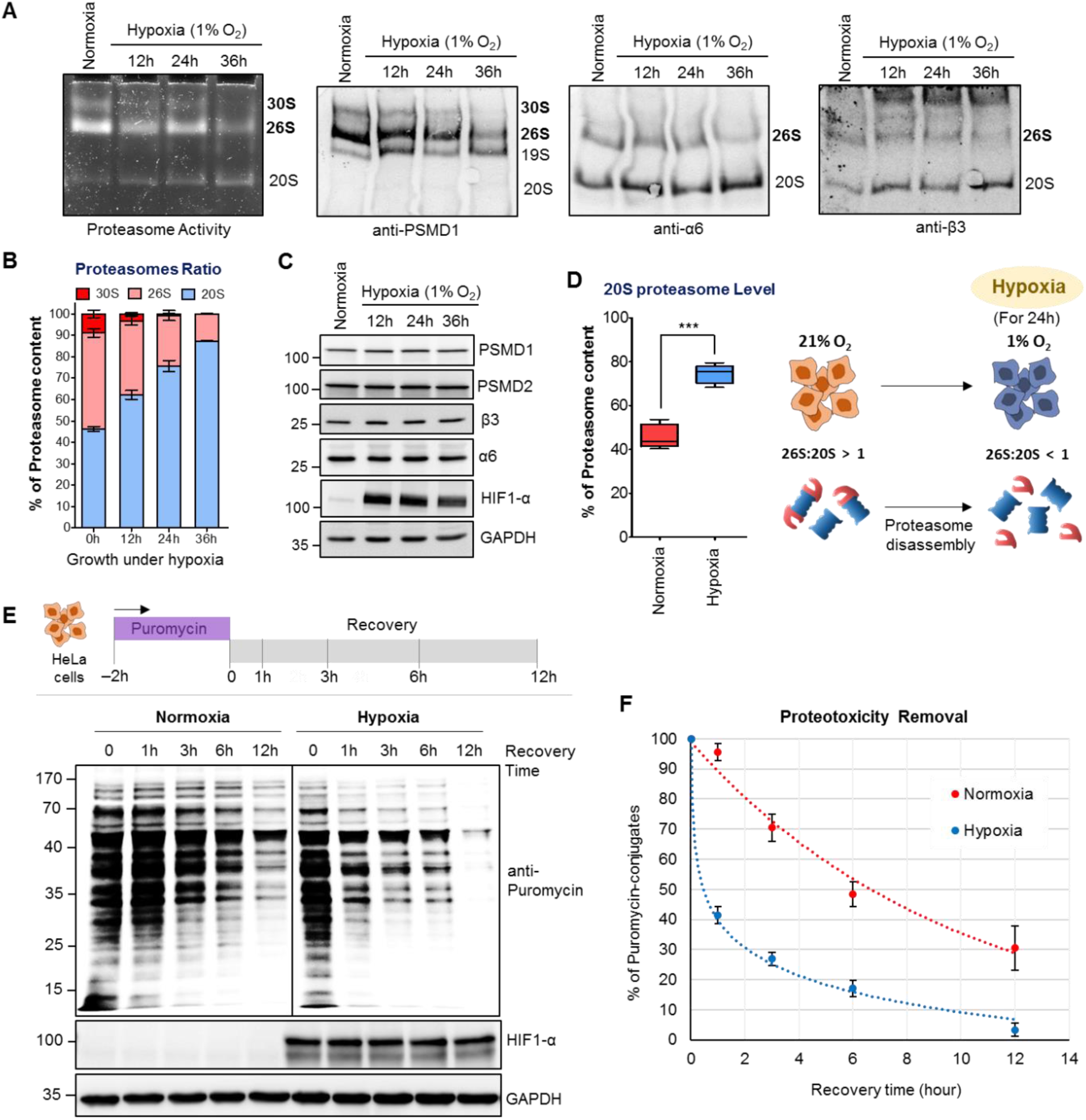
Hypoxia induced proteasome disassembly leads to rapid removal of nascent truncated proteins. (**A**) HeLa cells were grown either under normoxia (21% O_2_) or hypoxia (1% O_2_) for indicated time periods. Cells lysates were resolved in 4% native gel for proteasome in-gel activity assay or for native immunoblot (IB) using proteasome subunit antibodies (The 19S RP subunit PSMD1, and 20S CP subunits α6, β3). (**B**) The proteasome ratio in normoxic cells and hypoxic cells. Data represents the percentage of proteasome content (±SD) calculated from α6-native IB of three experiments at each time period. (**C**) The lysates from (A) were resolved by SDS-PAGE to detect proteasome subunits by IB (HIF1-α is the marker for hypoxia). (**D**) The levels of 20S proteasome in HeLa cells under normoxia and hypoxia (24 h). Data represents the average percentage of 20S proteasome as determined in (B). The cartoon illustrates proteasome disassembly under severe hypoxia into 20S and 19S sub-complexes. (**E)** HeLa cells were grown under hypoxia for 24 h followed by 2 h puromycin (5 µg/mL) treatment and then chased (Recovery) over 12 h. Cell lysates were resolved by SDS-PAGE to detect puromycin-conjugates by IB. (**F**) The rate of puromycin-conjugates removal under normoxia and hypoxia. Data represents the average percentage of puromycin-conjugates (±SEM) at each time points of the recovery phase, quantified from IB of three independent experiments. See also **Figure S1**

### Elevated 20S proteasome levels enhance intracellular degradation of a model substrate

In order to distinguish between the contribution of 26S and 20S proteasomes towards ubiquitin-independent proteolysis, we designed a model substrate for 20S proteasome based on human cyclin B1. Cyclin B1 generally undergoes proteasomal degradation upon its ubiquitination by activated APC/C complex at the end of M-phase of each cell cycle (Hershko et al., 1994; King et al., 1995). Structurally, cyclin B1 has a disordered N-terminal region, which contains a degron and 15 lysine residues (Yamano et al., 1998). Since it is structurally unstable, we considered whether this segment could serve as a direct target for 20S proteasome (**Figure S2A**). Hence, we designed a model substrate based on the N-terminal segment (88 amino acid) of cyclin B1 with HA tag at N-terminus (HA-CyclinB1-NT; **Figure S2B**). Upon expression of HA-CyclinB1-NT in HEK293 cells (**Figure S2C**), the protein was below detection level, presumably due to its rapid turnover (**Figure S2D**). In order to monitor intracellular turnover of HA-CyclinB1-NT, we fused a stable viral protease (NS3-pro) with its own cleavage site upstream to HA-CyclinB1-NT (**Figure 2A and S2E**). Following translation, NS3-pro autocatalytically cleaves the propeptide releasing HA-CyclinB1-NT (**Figure S2F and S2G**). We confirmed that HA-CyclinB1-NT synthesized and released by this method is degraded via proteasomal pathway (**Figure S2H**) rather than the lysosomal pathway (**Figure S2I**).

**Figure 2.**
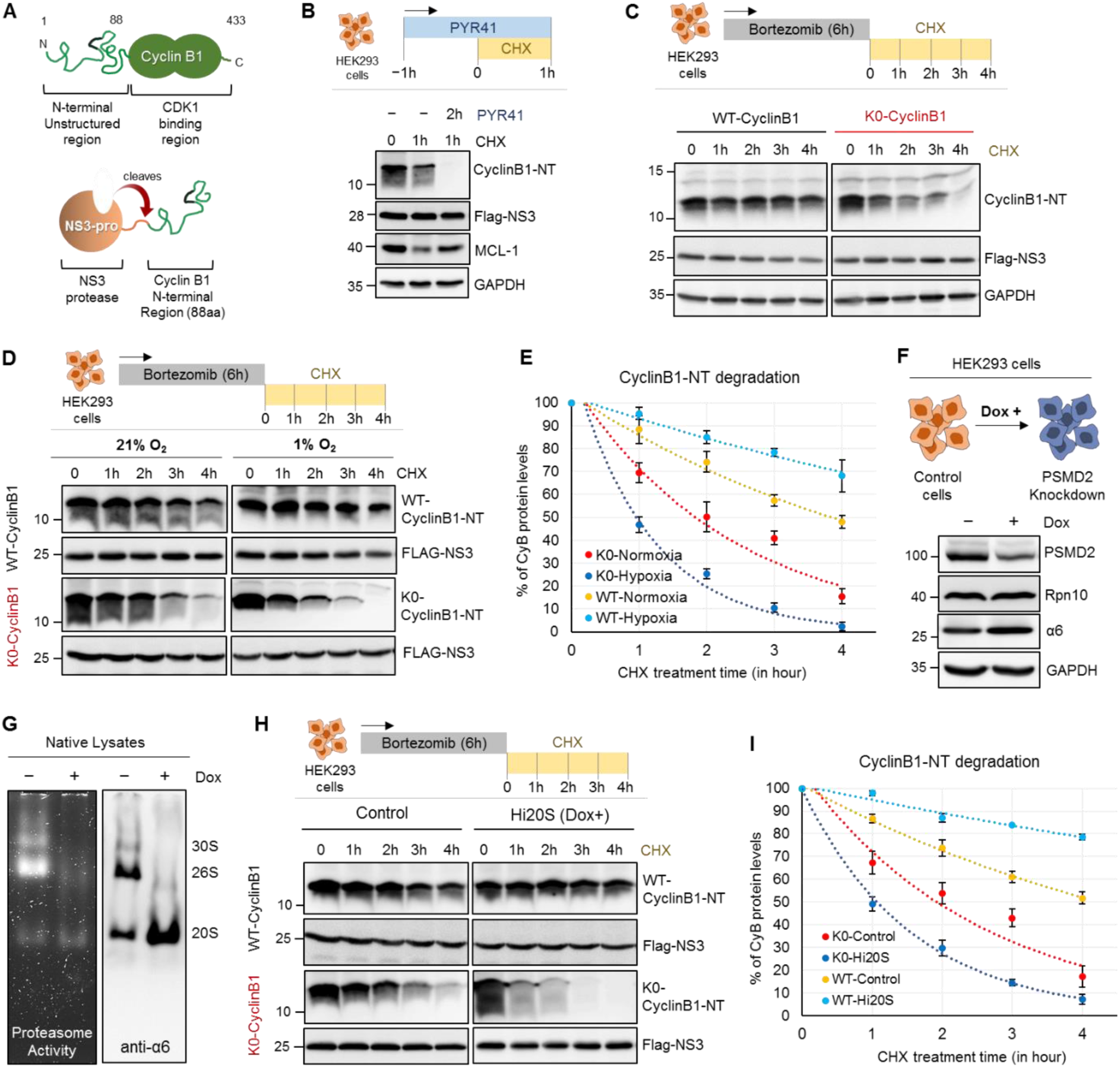
Non-ubiquitinated CyclinB1-NT is degraded faster by 20S than by 26S proteasome. (**A**) The cartoon represents: (Top) Cyclin B1 with its unstructured N-terminal region (NT) and C-Terminal CDK1 binding region, (Bottom) the fusion protein of cyclin B1 N-terminus (1-88 aa residue) with NS3-protease for expression in mammalian cells. (**B**) HEK293 cells expressing HA-tagged CyclinB1-NT were treated with 100 µg/mL cycloheximide (CHX) for 1 h, with or without E1-inhibitor (PYR41) at 25 µM. Various proteins in cell lysates were detected by IB. (**C**) HEK293 cells expressing either HA-tagged WT- or K0-CyclinB1-NT were treated with a pulse of bortezomib at 1 µM concentration for 6 h followed by cycloheximide chase. Various proteins in cell lysates were detected by IB. (**D**) HEK293T cells grown under normoxia or hypoxia for 24 h were treated with bortezomib followed by cycloheximide chase. Proteins in lysates were detected by IB. (**E**) Line graph represents WT- and K0-CyclinB1-NT protein levels under normoxia and hypoxia at each time-point of cycloheximide treatment. Error bar were calculated from the value for three independent experiments with ±SEM. (**F**) PSMD2 inducible knockdown HEK293T cells treated with doxycycline at 1 µg/mL for three passages to deplete PSMD2 protein levels. Immunoblot shows protein levels of proteasome subunits (Rpn10, α6). (**G**) PSMD2 inducible KD Dox treated and untreated cells from (F) were lysed with ATP-buffer and resolved by native gel to detect proteasome activity (left) or proteasome content (right). (**H**) PSMD2 inducible KD Dox treated and untreated cells expressing either HA-tagged WT- or K0- CyclinB1-NT were pulsed with bortezomib followed by cycloheximide chase. CyclinB1-NT levels were detected by IB. (**I**) WT- and K0-Cyclin B1-NT protein levels at each time-point of cycloheximide treatment under PSMD2 knockdown (Hi20S) or control conditions. Error bar represents the value for three independent experiments with ±SEM. See also **Figure S2 and S3**

Given that HA-CyclinB1-NT is a substrate for proteasome, we then tested whether its turnover is also dependent on ubiquitination. Surprisingly CyclinB1-NT degraded faster in presence of an E1-inhibitor, PYR-41 (**Figure 2B and S2J**), unlike MCL-1 which was stabilized in presence of PYR-41 as expected for a ubiquitin-dependent substrate (Zhong et al., 2005). To validate this result, we immunoprecipitated HA-CyclinB1-NT upon treatment of bortezomib and/or PYR-41 and confirmed that PYR-41 treatment prevents polyubiquitination of HA-CyclinB1-NT (**Figure S2K**). That an E1 inhibitor actually accelerates a protein turnover suggests that degradation of CyclinB1-NT is faster by a ubiquitin-independent pathway than by a ubiquitin-dependent mode. Moreover, the enhanced degradation of CyclinB1-NT in presence of PYR-41 is mediated by proteasome (**Figure S2L**). To strengthen the observation of accelerated degradation by a ubiquitin-independent pathway, we substituted all lysine residues with arginine in HA-CyclinB1-NT and followed degradation of this mutant (K0). Due to further rapid turnover, we could not detect any K0-CyclinB1-NT protein unless we treated cells with bortezomib (**Figure S3A**). Releasing cells from bortezomib treatment while blocking synthesis of new proteins (by cycloheximide) showed that K0-CyclinB1-NT degrades faster than WT-CyclinB1-NT (**Figure 2C**). We found no evidence for ubiquitination on K0-CyclinB1-NT (**Figure S3B**), which ruled out the possibility of non-lysine ubiquitination mediated degradation (Fajerman et al., 2004; Tait et al., 2007). Correspondingly, degradation of this mutant protein was insensitive to E1 inhibition (**Figure S3C**) and yet like its wild-type counterpart requires proteasome (**Figure S3D**). Taken together, the above observations strongly suggest that the unstructured segment of cyclin B1 can be degraded by proteasome, albeit faster via a ubiquitin-independent mode.

With a model proteasome substrate for ubiquitin-independent pathway, we revisited the intracellular proteolysis under hypoxia. We trans-expressed K0-CyclinB1-NT in HEK293 cells to track its degradation and observed more rapid turnover of K0-CyclinB1-NT under hypoxia compared to normoxia (**Figure 2D and 2E**). Since proteolysis of K0-CyclinB1-NT is ubiquitin-independent, faster degradation under hypoxia is likely due to elevated 20S proteasome under these conditions (**Figure S3E**). In contrast, the turnover of ubiquitin-dependent WT-CyclinB1-NT slowed-down in hypoxic cells (**Figure 2E**) correlated with 26S proteasome disassembly (**Figure 1B**). The 19S RP provides the ATP- and ubiquitin-dependency to the 26S proteasome; hence, a useful approach to decipher the 20S proteasome role in cells, is to deplete 19S RP subunits (Tsvetkov et al., 2015; Tsvetkov et al., 2017). In order to elucidate the contribution of the two proteasome species towards ubiquitin-independent proteolysis, we adapted a model system for altered proteasome ratio (Tsvetkov et al., 2015; Tsvetkov et al., 2017) by knocking down PSMD2 in HEK293T cells (**Figure 2F**). By this doxycycline (Dox) inducible system, we generated clones with low levels of 30S and 26S proteasomes but elevated levels of 20S proteasome (Hi20S cells) (**Figure 2G**). Next, we trans-expressed either WT- or K0-CyclinB1-NT in the Dox induced and uninduced PSMD2-KD cells to monitor their proteolysis. In Hi20S cells (Dox induced), turnover of K0-CyclinB1-NT was faster than in control cells (Dox uninduced), whereas degradation of WT-CyclinB1-NT slowed-down in Hi20S cells (**Figure 2H and 2I**). Taken together, these results suggest a potential contribution of 20S proteasome towards ubiquitin-independent degradation of CyclinB1-NT. Moreover, clearance of nascent truncated proteins was also faster in Hi20S cells (similar to hypoxic cells as in Figure 1F) than in control cells (**Figure S3F and 3G**).

### *In vitro*, 20S and 26S proteasomes degrade the same substrate differently

Our observations up to here distinctly suggest a potential role for 20S proteasome in ubiquitin-independent degradation of disordered proteins yet leave open the questions of: (i) the 26S proteasome involvement in such ubiquitin-independent proteolysis, and (ii) the apparent slower turnover upon ubiquitination of the same substrates. To address these issues, we turned to an *in vitro* setting where we could directly compare the enzymatic properties of purified 20S and 26S proteasomes on a synoptic set of differentially ubiquitinated substrates.

As a valid substrate for ubiquitin-independent mode of degradation, the disordered segment of cyclin B1 (as in Figure 2) was chosen to compare the proteolytic properties of 20S and 26S proteasomes *in vitro*. HA-tagged CyclinB1-NT (1-88 aa) was synthesized chemically with all lysine residues changed to arginine except Lysine 64 (**See Methods**). Then one, two or four ubiquitin moieties linked via Lysine48 were conjugated with isopeptide bonds to Lysine64 of HA-CyclinB1-NT by a series of chemical ubiquitination steps as described in detail in Methods (**Figure 3A**). In all of the cases (MonoUb-, DiUb- and TetraUb-conjugates) the proximal ubiquitin was “Myc” tagged, whereas the distal ubiquitin of TetraUb-HA-CyclinB1-NT was “Flag” tagged (**Figure 3B**) to aid in tracking the fate of specific ubiquitins. The purity and the integrity of the final reagents was confirmed by SDS-PAGE and immunoblots (**Figure 3B**). Human 26S proteasome (lacking proteasome-associated DUBs) and 20S proteasome were purified following the protocol mentioned in Methods; then their activity, integrity and composition were verified by native-PAGE, in-gel activity assay and SDS-PAGE (**Figure 3C**).

**Figure 3:**
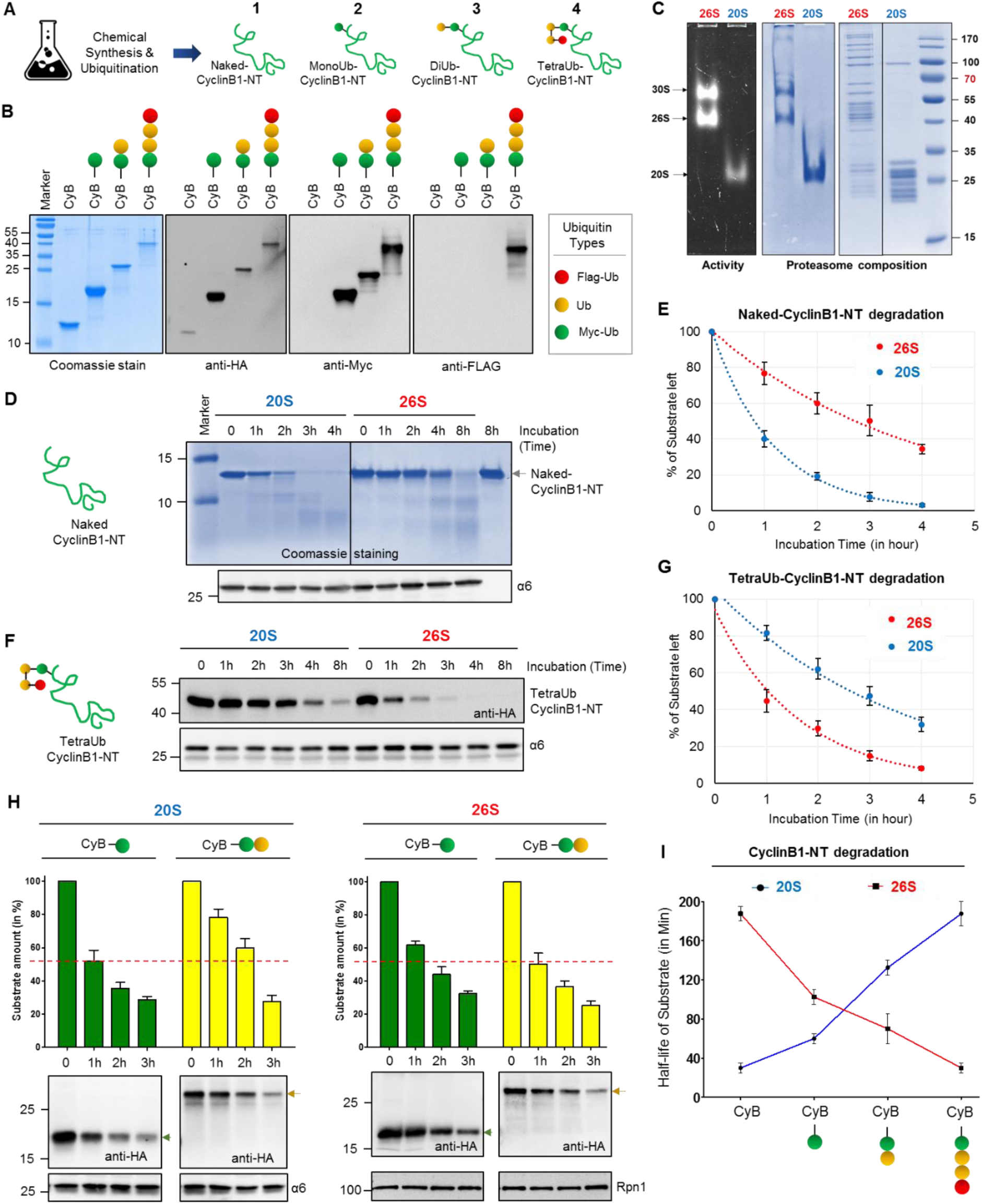
Chemically synthesized polyubiquitinated-cyclinB1-NT differentiates proteolytic activities of 26S and 20S proteasomes. (**A**) A cartoon represents the four chemically synthesized ubiquitin conjugates of CyclinB1-NT. Chemically synthesized HA-CyclinB1-NT (**1**) bearing a single mercaptolysine at position 64 was chemically ligated to synthetic MycUb-thioester to furnish MonoUb-CyclinB1-NT (**2**). Similar ligation to Myc-diUb^K48^-MPA yielded DiUb-CyclinB1-NT (**3**). For TetraUb-CyclinB1-NT (**4**), Myc-DiUb^K48^-HA-CyclinB1-NT bearing a single mercaptolysine at position 48 of the distal ubiquitin was ligated to FlagDiUb^K48^-thioester. All products were desulfurized to obtain the native isopeptide bond; see also Methods for chemical details. Ubiquitin units are untagged (yellow), Myc-tagged (green) or Flag-tagged (red). (**B**) All four ubiquitinated CyclinB1-NT conjugates (indicated in A) were resolved by SDS-PAGE and stained with Coomassie or probed for IB. (**C**) Purified Human 30S/26S and 20S proteasomes from human erythrocytes were resolved by native gel for in-gel activity (left), for native Coomassie (center), or by SDS-PAGE for Coomassie staining. (**D**) Naked-CyclinB1-NT was incubated at 37°C with purified 20S or 26S proteasome at 1:50 (Sub:Proteasome) molar ratio for indicated time periods. The reaction mixture was resolved by 16% Tris-Trycine denaturing gel for Coomassie staining and IB with α6 antibody as proteasome loading control. (**E**) The percentage of residual Naked-CyclinB1-NT substrate at each time point after degradation by either 20S or 26S proteasome. Error bar represents the value for three independent experiments with ±SEM. (**F**) TetraUb-CyclinB1-NT was incubated at 37°C with purified 20S or 26S proteasome at 1:50 (Sub:Proteasome) molar ratio for indicated time periods. The reaction mixture was was resolved by SDS PAGE for IB with anti-HA or α6 antibodies. (**G**) The percentage of residual TetraUb-CyclinB1-NT substrate at each time point after degradation by either 20S or 26S proteasome. Error bar represents the value for three independent experiments with ±SEM. (**H**) MonoUb-CyclinB1-NT and DiUb-CyclinB1-NT were incubated separately at 37°C with either purified 20S or 26S proteasome at 1:50 (Sub:Proteasome) molar ratio for indicated time periods. Each reaction mixture was resolved by SDS PAGE for IB (lower panel). The bar graph summarizes the values ±SEM quantified by anti-HA IB (normalized to either anti-α6 or anti-Rpn1 IB) from three independent experiments. (**I**) Half-life of CyclinB1-NT upon attachment of ubiquitin units. Error bar represents the value for three independent experiments with ±SEM.

By *in vitro* reactions we observed that Naked-CyclinB1-NT was proteolysed by purified 20S proteasome faster than by 26S proteasome (**Figure 3D and 3E**), whereas the converse was true for TetraUb-CyclinB1-NT (**Figure 3F and 3G**). However, the rate of proteolysis by each enzyme for its preferred CyclinB1-NT substrate (naked-for 20S and tetra ubiquitinated-for 26S), was similar (**Figure 3E and 3G**). While ubiquitination of a given substrate accelerated proteolysis by 26S proteasome, 20S proteasome was attenuated by conjugation of ubiquitin to the substrate. In order to test this premise further, we next considered shorter ubiquitin modifications on the same substrate (MonoUb-CyclinB1-NT and DiUb-CyclinB1-NT). A gradual decrease in rate of CyclinB1-NT processing by 20S proteasome was observed, that proportionate to the number of ubiquitin units attached; whereas the inverse phenomena was detected for 26S proteasome (**Figure 3H and 3I**). For this synoptic set of substrates, ubiquitination shifts a 20S-substrate to a 26S-substrate with the implication that conditional deubiquitination could convert back a ubiquitin-conjugate (disordered) to a preferred substrate for 20S proteasome. Comparison of *in vitro* 20S and 26S proteolytic rates elucidates why *in vivo* ubiquitin-independent degradation of an unstructured protein (CyclinB1-NT) is faster than ubiquitin-dependent degradation. For this given substrate, ubiquitination is apparently the rate-limiting step in ubiquitin-proteasome degradation pathway.

### Signature products of 20S proteasome activity are present in cells

The disordered nature of CyclinB1-NT renders it a dual substrate for the two proteolytic enzymes – 20S and 26S proteasomes – each with a preference for a ubiquitination state (as in Figure 3). Since the two enzymes proteolyse their substrate with different rates, they may also generate different peptide products from the same substrate. Defining the product repertoire would enable us to identify their signature activities under different cellular conditions. Therefore, we incubated Naked-, MonoUb- and TetraUb-CyclinB1-NT with either purified proteasome separately, and terminated reactions at early time points to minimize product reprocessing (**Figure 4A**). MS/MS analysis of the isolated peptide products (**Figure S4A**) showed differential cleavage patterns, P1 sites, and peptide size distribution for each substrate (**Figure 4B**), although 20S and 26S proteasomes did not differ in their amino acid preference at cleavage sites (**Figure S4B**). 20S proteasome generated longer peptides from all substrates than that of 26S proteasome (**Figure 4C**). Moreover, 20S proteasome showed a marked preference for specific sites within a given substrate, in essence defining its signature cleavage patterns (**Figure 4D**). Repeating the analysis with a ubiquitinated version of the same substrate decreased the number of 20S preferred cleavage sites. The diverse product profile of naked-CyclinB1-NT cleavage reflects an overall randomness of the cleavage pattern by 20S relative to 26S proteasome and this randomness declined with ubiquitination of substrates (**Figure S4C**). Identifying these signature activities in cells could differentiate between the proteolytic contribution of 20S and 26S proteasomes. To do so, we ectopically expressed K0-CyclinB1-NT in control and Hi20S cells and performed intracellular peptidomic analysis (**Figure 4E**). A larger variety of K0-CyclinB1-NT peptides were identified in Hi20S cells than in control cells. Analysis of cleavage frequency at P1 positions of K0-CyclinB1-NT in Hi20S cells recapitulated the *in vitro* 20S signature cleavage pattern (**Figure 4F**). These Hi20S cells also generated longer peptides from K0-CyclinB1-NT than the control cells (**Figure 4G**). Moreover, analysis of total peptidome also showed a significantly larger peptide size distribution in Hi20S cells (**Figure 4H**), which again reiterated 20S signature behavior. Tracing back from the peptide pool identified 787 target proteins unique to Hi20S cells, with a higher median protein disorder score (**Figure 4I and S4D**). Similarly, in hypoxia-induced Hi20S cells, total peptidome showed a significantly larger peptide size distribution (**Figure S4E**) and a significant higher median substrate-protein disorder score (**Figure S4F**) than that of normoxic cells. Together, these results identify the presence of 20S signature activity in cells and suggest that 20S proteasome is actively contributing to overall proteolytic activity in mammalian cells.

**Figure 4:**
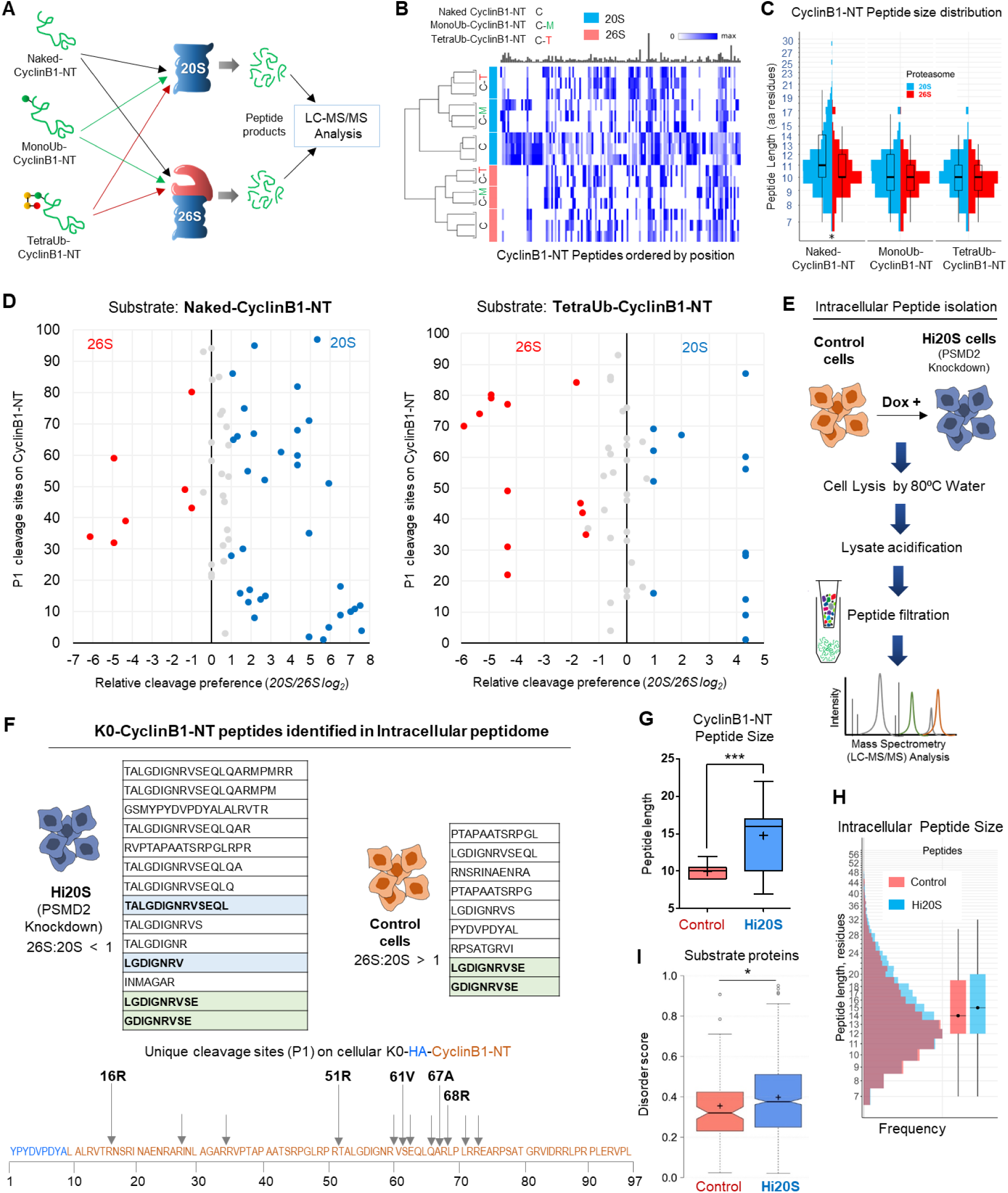
20S proteasome proteolytic signature detected in cells. (A) Naked-CyclinB1-NT, MonoUb-CyclinB1-NT or TetraUb-CyclinB1-NT were incubated separately at 37°C with either purified 20S or 26S proteasome at 1:50 (Substrate:Proteasome) molar ratio. Peptide products at time points shorter than substrate half-life for each reaction were isolated and analyzed by LC-MS/MS. (**B**) Heat map represents the CyclinB1-NT-derived peptides obtained from each substrate by 20S or 26S proteasome (triplicate or duplicate experiments). Each peptide product is positioned along the 99 aa primary sequence of CyclinB1-NT (based on its N-terminal residue). Color intensity reflect PSM counts for each peptide normalized to the maximum observed counts. The bar plot above (in grey) represent relative sizes of the corresponding maxima. The dendrogram to the left was obtained by performing MDS analysis on the PSM counts of the top 100 peptides and clustering the samples based on the corresponding MDS distances. (**C**) Size distribution of CyclinB1-NT peptide products in each sample from (A). Overlaid box plot summarizes median peptide length. (**D**) Scatter plot represents the relative cleavage preference for each proteasome species at P1 positions on CyclinB1-NT in its Naked- and TetraUb-forms. Cleavage preference for each enzyme was calculated from the MS/MS count of each peptide contributing to a given P1 site and the relative ratio plotted as log_2_ (described in Methods). The Y-axis represents the amino acid residue number of the HA-CyclinB1-NT sequence. Red/Blue dots indicate a ≥2 fold preference (Red, 26S; Blue, 20S; grey, no significant preference). (**E**) The method for intracellular peptide isolation and identification. PSMD2 inducible knockdown HEK293 cells (±doxycycline) expressing HA-tagged K0 Cyclin B1-NT were grown with aminopeptidase inhibitor (1 μM CHR 2797) for 24 h. Intracellular peptides were captured as described in Methods and subject to non-tryptic LC-MS/MS analysis. (**F**) Tables shows all peptides of K0-CyclinB1-NT identified from intracellular peptidomics of Hi20S and control cells. Peptides highlighted in light-blue match to peptides obtained from cleavage of naked-CyclinB1-NT by 20S proteasomes in vitro. The light-green highlighted peptides were identified in both cell types. In the illustration (bellow), the amino acid sequence corresponds to the HA-CyclinB1-NT (K0) trans-expressed in cells and arrows represent unique cleavage sites identified on K0-CyclinB1-NT in Hi20S cells. The assigned P1 positions match to the preferred P1-positions for 20S proteasomes cleavage in vitro of naked-CyclinB1-NT. (**G**) Size distribution of intracellular CyclinB1-NT derived peptides (line, median; + mean size) obtained from control and Hi20S cells (from triplicate dataset). (**H**) Size distribution size of all intracellular peptides obtained in control or Hi20S cells from triplicate datasets. Box plot on right represents median peptide size. (**I**) Disorder score distribution of potential proteasome substrates unique to each cellular condition (control, Hi20S). Proteins were assigned from intracellular peptide captured, and scored based on intrinsic disordered elements (notch, median; + mean). See also **Figure S4**

### 20S proteasome degrades ubiquitin along with an unstructured conjugate

The role of 26S proteasome-associated DUBs is to release the ubiquitin-tag from the substrate for recycling purpose. Compromising this deubiquitinase activity can lead to ubiquitin degradation along with the substrate (Guterman and Glickman, 2004; Hanna et al., 2003; Singh et al., 2016). Having observed that 20S proteasome was able to proteolyse ubiquitinated CyclinB1-NT *in vitro* (as in Figure 3), we were intrigued how 20S, without associated deubiquitinase or unfoldase activities, could handle the ubiquitin tag. We found that 20S proteasome proteolysed TetraUb-CyclinB1-NT in its entirety without releasing any free ubiquitin, unlike 26S proteasome that released tetraubiquitin en-bloc (**Figure 5A and S5A**). Likewise, MonoUb- or DiUb-CyclinB1-NT were also proteolyzed entirely by 20S proteasome whereas 26S proteasome released the ubiquitin chains while proteolysing the substrates (**Figure 5B**). Having further observed that ubiquitin in free chains were not proteolysed (**Figure S5B and S5C**), we hypothesized that 20S proteasome most likely can degrade ubiquitin when it is conjugated to an unstructured substrate. In order to validate ubiquitin proteolysis, we isolated all peptide products from 20S proteasome reactions subjected to non-tryptic MS/MS analysis (**Figure 5C**). For all ubiquitin conjugates, peptides spanning the ubiquitin sequence were identified. In order to confirm that the entire length of a polyubiquitin chain can be proteolysed by 20S proteasome, we differentially tagged the distal and proximal ubiquitin units of TetraUb-CyclinB1-NT with Flag or Myc epitopes, respectively. Chimeric peptides spanning these epitopes provide evidence for proteolysis of the entire TetraUb chain (**Figure 5C**).

**Figure 5:**
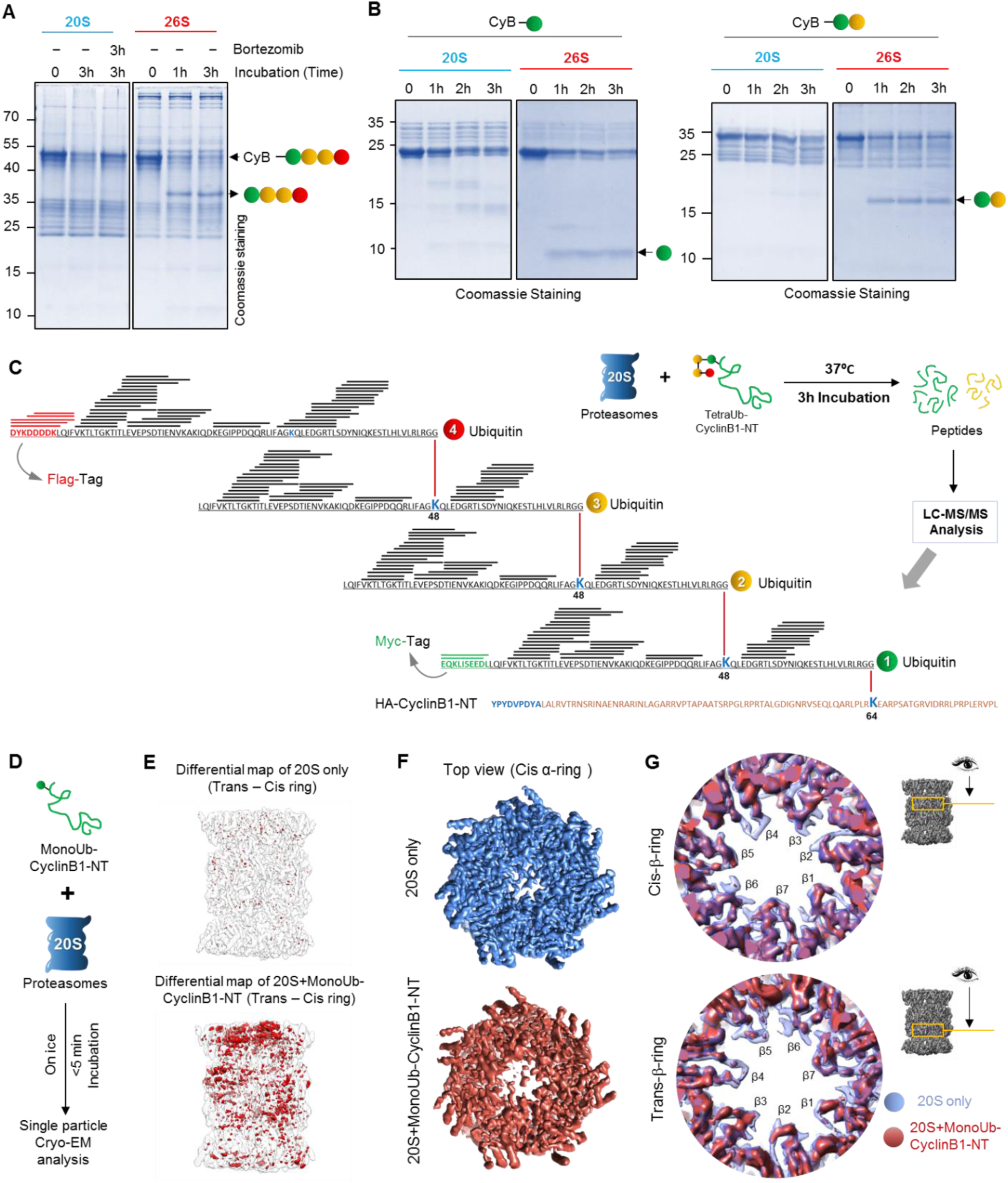
Conjugated ubiquitin is proteolysed by 20S proteasome in vitro. (**A**) TetraUb-CyclinB1-NT, and (**B**) MonoUb- or DiUb-CyclinB1-NT were incubated at 37°C with purified 20S or 26S proteasome at 1:50 (Substrate:Proteasome) molar ratio for indicated time periods. The reaction mixture was resolved by Tris-Tricine PAGE and stained with Coomassie. Brotezomib was added to one reaction to inhibit 20S proteasome activity. (**C**) The illustration represents the ubiquitin peptides obtained from TetraUb-CyclinB1-NT proteolysed by 20S proteasome detected by MS/MS analysis overlaid (as black lines) on the primary sequence of the polyUb chain. Red represents the Flag-tag sequence on the distal ubiquitin unit, while green represents the Myc-tag sequence on the proximal ubiquitin. (**D**) The work-flow model depicts sample preparation for cryo-EM of MonoUb-CyclinB1-NT incubated with purified 20S proteasome at 5:1 molar ratio on ice (briefly), frozen within 5 min and then vitrified for further cryo-EM single particle analysis. (**E**) Difference map rendering the symmetric pattern between the trans- and cis-rings of 20S alone map (upper) and 20S+monoUb CyclinB1-NT map (lower). The difference map (red density) was generated by using a 180° rotated (around the pseudo 2-fold axis) map minus its original unrotated-map (transparent grey). (**F**) The top view from cis α-ring of electron density maps of 20S-only (Blue) and S1 class of 20S+monoUb CyclinB1-NT (brick red) dataset. (**G**) Comparison of 20S alone (transparent blue) and S1 class of 20S+monoUb CyclinB1-NT (brick red) maps in the two β-ring annulus regions. This analysis suggests that for the 20S+monoUb CyclinB1-NT map, there are density loss in some annulus loop and β-sheet regions exposed or close to the catalytic chamber in both β-rings. See also **Figure S5**

Degradation of a highly stable globular protein such as ubiquitin by a 20S protease, which lacks an unfoldase activity, is quite surprising. In order to demonstrate association of a ubiquitin-conjugate with 20S, we performed cryo-EM single particle analysis on purified human 20S proteasome incubated with MonoUb-CyclinB1-NT and compared it with 20S alone (**Figure 5D**). Classification of the data from untreated 20S (20S alone) resolved one class of symmetric proteasome in the resting state (further classification did not yield significant differences between classes; **See Methods and Figure M14**). Similar analysis of substrate-incubated 20S resulted in a class (S1) comprising ∼29.1% of the particles in an asymmetric appearance between the cis and trans α-rings, especially at the gate region (**See Methods and Figure M15**). Incubating with substrate for a mere 5 minutes was sufficient to induce a fundamental conformational asymmetry to the 20S barrel by destabilizing subunits in one α-ring radiating into the central beta cavity (**Figure 5E**). Notable is the loss of electron density in the center of the cis α-ring especially in the N-termini of α2/α3/α4 subunits (that usually extend to cover the gate, but now showing disconnected features; **Figure 5F, S5D**). This partially “open gate” conformation reminiscent of activated 20S CP (Ding et al., 2019; Groll et al., 2000; Osmulski and Gaczynska, 2002; Rabl et al., 2008; Toste Rego and da Fonseca, 2019; Whitby et al., 2000), is apparently sufficient to facilitate entry of the branched polypeptide of CyclinB1-NT conjugated to ubiquitin. Overlaying electron density maps also highlight flexibility of beta annulus loops in both beta rings adjacent to the catalytic active sites (**Figure 5G**). These substrate-induced conformational changes are all consistent with a protein substrate entering from one side into the inner proteolytic chamber. Allosteric effects between the α-gate and β-active sites have been suggested to couple substrate translocation and proteolysis (Osmulski et al., 2009). The surprising ability of 20S proteasome to proteolyse ubiquitin along with its attached target protein stands out as a signature property of 20S given that the bulk of conjugated ubiquitin is recycled from the 26S proteasome by virtue of its deubiquitinase activity.

### 20S proteasome contributes to degradation of ubiquitin-conjugates under hypoxia

To search for the signature activity of 20S proteasome to degrade ubiquitin, control and Hi20S cells (**Figure 6A**) were treated with puromycin to induce misfolded proteins. Co-immunoprecipitation confirmed that part of the puromycinated nascent polypeptides are indeed ubiquitinated and are cleared from the cells engineered for high 20S content (**Figure 6B**). Nevertheless, control cells removed these polyubiquitinated conjugates faster than the non-ubiquitinated cognates, in accord with the expected activity of 26S proteasome. In a chase experiment, we observed ubiquitin units recycled from polyubiquitinated conjugates return to the pool of free monoUb in control cells, in contrast to Hi20S cell where mono ubiquitin did not accumulate over time (**Figure 6C**). This observation supports the ability of 20S to degrade ubiquitin-conjugates in cells. Next, we returned to hypoxia, the physiological condition that promotes the Hi20S state (**Figure 6D**). After 12 hours under hypoxia, high molecular weight polyubiquitinated conjugates accumulated, possibly due to impairment of 26S function (as in Figure 1). However, during longer hypoxic incubation, these conjugates cleared off without concomitant generation of free mono-ubiquitin units (**Figure 6E**) correlates with increased 20S proteasome content. Notably, it was Lysine48-linked polyubiquitinated conjugates that were removed, rather than Lysine63-linked (**Figure 6E**). Moreover, this decrease in polyubiquitinated conjugates was an outcome of degradation and not a loss-of-function of E1 activity (**Figure S6A**).

**Figure 6:**
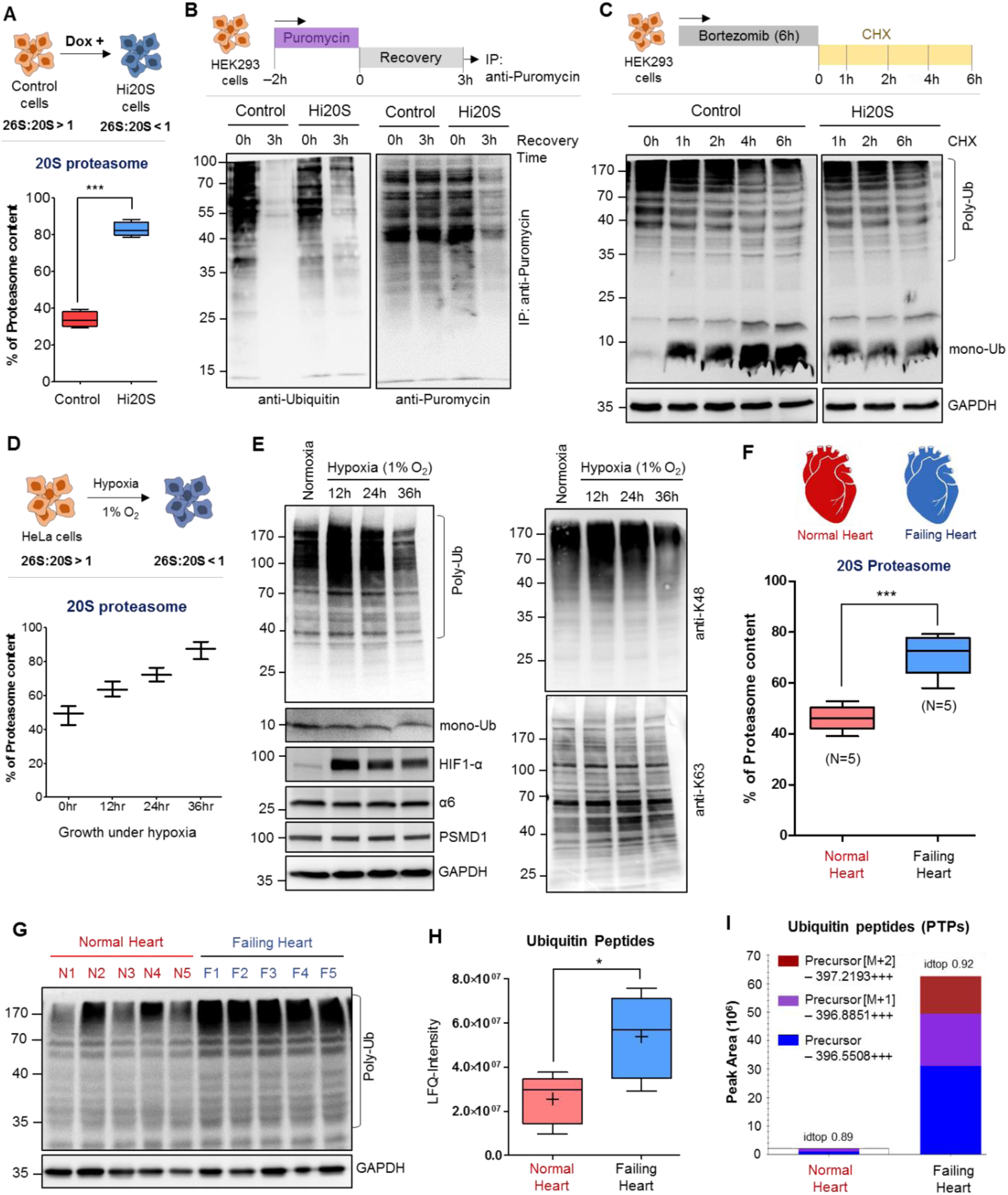
20S proteasome facilitate degradation of ubiquitin conjugates under hypoxia. (**A**) 20S proteasome content of control and Dox-induced Hi20S cells. Data represents the average value of three independent experiments (±SD). (**B**) Control and Hi20S cells were pulsed with puromycin for 2 h. Residual puromycin-conjugates were immunoprecipitated at 0 and 3 h of recovery, and IB for ubiquitin and puromycin content. (**C**) Control and Hi20S cells were pulsed with bortezomib for 6 h and chased with cycloheximide for up to 6 h of recovery; samples taken for ubiquitin IB. (**D**) 20S proteasome content of HeLa cells grown under normoxia (21% O_2_), or under hypoxia (1% O_2_) for indicated time. Data represents the percentage of proteasome content (±SD) calculated from α6 native IB of three experiments at each time period. (**E**) Polyubiquitin content of HeLa cells grown under normoxia or under hypoxia. IB using antiUb (left) or antiK48 and K63 linkages (right). (**F**) Proteasome content of human heart muscle tissue under Failing (N = 5) or Normal (N = 5) conditions. Proteasome enrichment from frozen tissue followed the protocol in Methods and ratio of proteasome species for each sample was measured from anti-α6 probed native IB. The box plot represents the percentage of 20S proteasome of Normal heart samples (N = 5) and Failing heart samples (N = 5). (**G**) Polyubiquitin content of human heart muscle lysates IB using antiUb. (**H**) Ubiquitin-derived peptides in heart muscle tissues. Human frozen heart muscle tissue under Failing (N = 5) and Normal (N = 5) conditions were subjected to intracellular peptide isolation following the protocol in Methods quantitatively analyzed for ubiquitin peptides by LC-MS/MS. The plot represents the LFQ-intensity of ubiquitin calculated from its peptides in each sample (line, median; +, mean). (**I**) Intensity of Ubiquitin peptides associated with enriched proteasome from heart muscle tissues. Proteasome was isolated from frozen human heart tissue, and integrated intensity proteasome trapped peptides (PTPs) were calculated by MS/MS. Bar graph summarizes ubiquitin peptides (from all charged forms) in all samples from both conditions. See also **Figure S6**

One example of a chronic-hypoxic pathological condition is ischemia-associated hypoxia as in end-stage failing heart (Giordano, 2005). Indeed, quantifying proteasome content in human myocardial tissue demonstrated an abnormally high ratio of 20S to 26S in failing heart (**Figure 6G and S6B**). We observed elevated polyUb-conjugates in failing heart tissue similar to our cultured cells under hypoxia, suggesting an impairment in 26S proteasome function (**Figure 6H**). An inevitable side effect of these conditions based on our observations so far would be proteolysis of ubiquitin. Evidence for proteolysis of the ubiquitin protein in failing heart tissue was procured by whole tissue peptidomics. Five ubiquitin peptides were identified in failing heart tissue with a significantly higher integrated intensity than in normal heart tissue (**Figure 6I, S6C and S6D**). Proteasome Trapped Peptides (PTPs) from a proteasome-enriched fraction of heart tissue identified the same ubiquitin-derived peptides in the 20S core complex, validating the source of the generated peptides (**Figure 6J, S6E and S6F**). Furthermore, a preference for disordered substrates, another 20S signature (Myers et al., 2018; Suskiewicz et al., 2011), corroborates elevated 20S function in these tissues (**Figure S6G and S6H**). Despite a skewed 20S/26S ratio under these hypoxic conditions, breakdown of polyUb-conjugates continues due to 20S proteasome function. Altogether, these results define an ability of 20S proteasome to degrade ubiquitin attached to a conjugate, a signature behavior distinct from 26S proteasome.

### 20S proteasome contributes to cell survival by alleviating proteotoxicity under hypoxia

So far, we have described how 20S proteasome can efficiently proteolyses truncated/misfolded proteins, generates a characteristic peptide product pattern, and degrades intact ubiquitin-conjugates, all distinct behaviors from 26S proteasome. In order to understand how this signature activity is beneficial to cells, we exposed cell culture to increasing proteotoxicity by puromycin treatment. Hi20S cells showed better survival at all puromycin concentrations than control cells (**Figure 7A, 7B, S7A**). Since under hypoxia 20S proteasome content is elevated, we further challenged the cells with increasing concentrations of puromycin. Similar to the Hi20S cells, hypoxic cells survived better at all concentrations of puromycin in comparison to normoxia (**Figure 7C, S7B**). Although cells under hypoxia showed low 26S proteasome content they grew at the same rate as under normoxia (**Figure S7C**), suggesting that in hypoxia 20S is an active proteasome species contributing to the overall proteostasis for cell survival.

**Figure 7:**
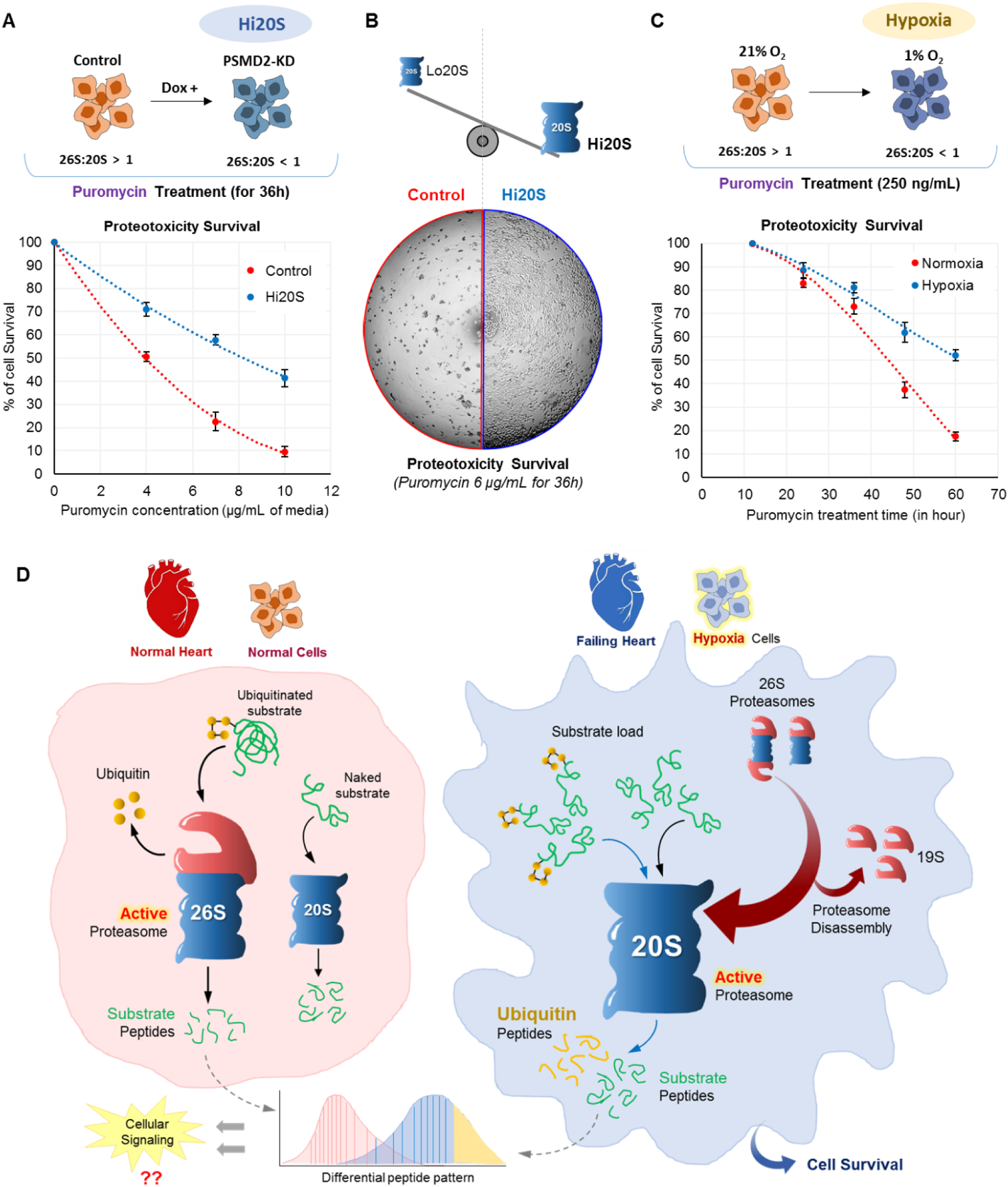
Hi20S proteasome condition facilitates cell survival under proteotoxic stress. (**A**) Cell survival at increasing puromycin exposure. Control and Hi20S cells were grown for 36 h in the presence of increasing puromycin concentrations. Cell survival was quantified by MTT assay. Data represents the average of three experimental values (±SD). (**B**) Phase contrast image of culture plate showing survival of control (Lo20S) and Hi20S cells after 36 h of puromycin treatment (6 μg/mL). (**C**) Cell survival upon puromycin exposure during hypoxia. HeLa cells were grown under normoxia and hypoxia (1% O_2_) with puromycin (250 ng/mL) for up to 60hr. Cell survival was quantified by MTT assay. Data represents the average of three experimental values (±SD). (**D**) The model demonstrates the division of roles between 26S and 20S proteasomes. Under “normal” conditions, each proteasome degrades its preferred substrates (left). Under hypoxic conditions, proteasome disassembly leads to high 20S levels, which participates in bulk protein degradation, including some ubiquitin-conjugates. Consequently, the intracellular peptidome changes. See also **Figure S7**

## Discussion

The 26S proteasome is a dynamic complex that exists in equilibrium with its free core particle, the 20S. Under normal growth conditions, free 20S core particle is quite abundant, though varies in different cell types or in response to environmental conditions. As described herein, we found that the ratio of 20S to 26S increases under severe hypoxia and in end-stage failing heart, a pathological condition that occurs at ischemia-associated hypoxia. Shortly after the discovery of the proteasome, it was thought that independently of the ubiquitin-dependent ATP-stimulated 26S proteasome “the (20S) proteasome by itself constitutes an enzyme system that must be important in the turnover of certain cytosolic and nuclear proteins” (Arrigo et al., 1988). However, lacking ubiquitin-receptors and ATP-dependent unfoldases, 20S is usually considered a less versatile protease than 26S proteasome, limited to only unstructured polypeptides. We now show that 20S does indeed contribute to bulk degradation of proteins according to cellular demands. In the case of hypoxia or failing heart, evidently cells alter proteasome ratio harnessing the efficiency of 20S proteasome for removing toxic proteins; an adaptive response to cellular environment (**Figure 7D**). Moreover, degrading these misfolded proteins persists even when they are conjugated to polyUb chains, justifying the definition of 20S as an active “Proteasome”.

An outcome of 20S proteasome activity is ubiquitin degradation (**Figure 7D**). At proteostasis, 26S and 20S segregate substrates between them based on their preferences: polyubiquitinated proteins to 26S and non-ubiquitinated (misfolded) polypeptides to 20S. Shortage of 26S, as we observed in early stages of hypoxia and in failing heart, may lead to temporary accumulation of Lysine48-linked polyUb conjugates. Under these perturbed conditions, 20S proteasome participates in removal of ubiquitinated conjugates, but without recycling ubiquitin. We speculate that prolonged Hi20S conditions would lead to a new proteostatic balance whereby the cellular content of both 26S proteasome and Lysine48-linkaged conjugates decrease simultaneously. In parallel, cellular DUBs could also aid 20S proteasome function by removing some of the polyUb-tags from substrates. In the current study we provide direct evidence for the ability of the 20S proteasome to proteolyse a ubiquitin chain (TetraUb) attached to a substrate, *in vitro*. Cryo-EM analysis provides further insight to the conformational changes of 20S proteasome at the initial steps of the reaction: engagement of a ubiquitin-conjugate induced sufficient flexibility in the α-ring, especially in the gate region, to enable isopeptide-linked branched polypeptides to enter through the 20S gate and reach the β-active sites. Interesting to note, we did not find evidence for proteolysis of free ubiquitin nor of ubiquitin chains, unless attached to an unstructured substrate. In essence, here the unstructured protein served as the targeting signal for ubiquitin degradation. This explains the source of ubiquitin-derived peptides in cells that rely on 20S proteasome for degradation of a significant portion of their proteome (Figure 5).

20S and 26S proteasomes generate different peptide products from the target proteins, even though they have the same catalytic β-active sites. Since we did not detect any difference in amino acid preference for cleavage at the P1 site by 20S or 26S, the differential cleavage pattern might be influenced by allosteric regulation from the 19S regulatory particle. By directly comparing purified 20S and 26S proteasome action on a set of identical substrates that differ only in the extent of ubiquitin conjugation, it was possible to deduce the unique properties of each protease. Although ubiquitin does not serve as a targeting signal for 20S proteasome, its conjugation does affect the overall product pattern. In general, 20S proteasome generates a greater variety and comparatively longer peptides from a given substrate relative to 26S proteasome. Longer peptides could retain secondary structure to some extent, which increases their potential to serve signaling roles (de Araujo et al., 2019). As a consequence, tweaking 26S/20S proteasome ratio would alter the intracellular peptidome, with ramifications to downstream signaling pathways.

To summarize, in this study we establish signature activities of 20S proteasome that are distinct from 26S proteasome: (1) Cleavage site preference, (2) Longer peptide products, (3) Disordered substrate preference and (4) Ubiquitin degradation. Both proteasomes differ not only in their substrate preference, but also in their product outcome. Their signatures can be used to predict cells that harbor altered 26S/20S proteasome ratio. Uncharacteristically high levels of 20S proteasome have been reported for a variety of stress conditions, including nutrient starvation (Bajorek et al., 2003), ROS (Livnat-Levanon et al., 2014), hypoxia (Predmore et al., 2010), and as a general mark of aging (Chondrogianni et al., 2015; Raynes et al., 2016). With the understanding that the 20S proteasome is a functional enzyme, its presence may not be merely a consequence of the stress (“disassembled proteasomes”) but rather provide a gain-of-function for survival. As demonstrated in current study, efficient removal of damaged proteins during hypoxia is attributed to elevated 20S proteasome. Likewise in aging, increased 20S proteasome levels may compensate for the loss of protein quality control and increased proteotoxic load. In this scenario, 20S proteasome is more than a marker for aging, as it may afford for healthy aging. Thus, conditional increase of 20S proteasome generates a de facto “Emergency proteasome” as an adaptive response to environmental or intracellular stress.

## Acknowledgments

We thank: Peter Tsvetkov (Broad Institute of MIT and Harvard) for sharing Cell lines and shRNA constructs; Noa Reis (Technion) for critical advice and aid in cloning; Inbar Magid Gold (Technion) for recombinant enzymes; Tehila Bar Kafra (Technion) for coding MS analysis software; N. Dahan (Technion LSE facility) for aiding FACS and microscopy; Ilana Navon and Tamar Ziv and rest of the team of Smoler Proteomics Facility for performing mass spectrometry analyses, Cheng Huang (Enzo) and Ralf Schittenhelm from Monash Biomedical Proteomics Facility for performing mass spectrometry analyses. This research is supported in part by Israel Science foundation grants (755/19 to MHG; 179/15 to AB; 1623/17 and 2167/17 to OK). NSFC-ISF (2512/18 to YC and MHG), NSF-BSF (2017727 to MHG) and MSCA-IF Horizon-2020 (2024849 to IS). IS is an MSCA fellow, PS is a TICC fellow, AB is a Jordan and Irene Tark Academic Chair, MHG is The Israel Isaac and Natalia Kudish Academic Chair.

## Author Contributions

IS, YC, OK, AB and MHG designed the research and developed the project. IS and PS conducted cell biology experiments. IS and RM conducted cloning and biochemical experiments. SMM and SKS performed chemical synthesis of proteins. MPS carried out proteomic sample preparation. AR and OK analyzed proteomic data. CX, ZD, YW and YC performed cryo-EM analysis. SD procured human heart tissue. IS, OK, AB and M.H.G. wrote the first version of the manuscript. All authors read and commented on the manuscript.

## Declaration of Interests

Authors declare no conflict of interest.

## METHODS

### Plasmids and cloning

The DNA fragment of N-terminal part of Cyclin B1 (1-88 aa) with HA-tag (HA-CyclinB1-NT) was cloned into pCMV10-3XFlag vector. Both wild type and all lysine to arginine (K0) mutant fragments were used for cloning. (**Figure S2D**). Further, to express NS3 fused HA-CyclinB1-NT in mammalian cells we fused NS3Pro at the N-terminus of HA-CyclinB1-NT by designing two DNA fragments as bellow and cloning into EcoRI and HindIII sites of pYFP-SmaSh vector (Addgene: 111500).

#### Fragment 1

EcoRI-Kozak Sequence-ATG-3XFLAG-Linker-NS3Pro-NS5A5B cleavage site-BamHI DNA sequence: 5’ gaattcgccaccatggactacaaagaccatgacggtgattataaagatcatgacatcgattacaaggatgacgatgacaagatggcgcccatcacggcgtacgcccagcagacgagaggcctcctagggtgtataatcaccagcctgactggccgggacaaaaaccaagtggagggtgaggtccagatcgtgtcaactgctacccaaaccttcctggcaacgtgcatcaatggggtatgctgggcagtctaccacggggccggaacgaggaccatcgcatcacccaagggtcctgtcatccagatgtataccaatgtggaccaagaccttgtgggctggcccgctcctcaaggttcccgctcattgacaccctgtacctgcggctcctcggacctttacctggtcacgaggcacgccgatgtcattcccgtgcgccggcgaggtgatagcaggggtagcctgctttcgccccggcccatttcctacttgaaaggctcctcggggggtccgctgttgtgccccgcgggacacgccgtgggcctattcagggccgcggtgtgcacccgtggagtggctaaagcggtggactttatccctgtggagaacctagagacaaccatgagatccccggtgttcacggacgtgggcaggatcgtcttgtccgggaagccggcaggcagtagcggaagcagtattatacctgacagggaggttctctaccaggagttcgaagatgtcgtgccatgctcaatgggat cc 3’

#### Fragment 2

BamHI-HA-Cyclin B1-NT-HindIII DNA Sequence: (1) WT-CyclinB1-NT: 5’ ggatccatgtatccatatgatgttccagattatgctatggcgctccgagtcaccaggaactcgaaaattaatgctgaaaataaggcgaagatcaacatggcaggcgcaaagcgcgttcctacggcccctgctgcaacctccaagcccggactgaggccaagaacagctcttggggacattggtaacaaagtcagtgaacaactgcaggccaaaatgcctatgaagaaggaagcaaaaccttcagctactggaaaagtcattgataaaaaactaccaaaacctcttgaaaaggtacctatgtgaaagctt 3’ DNA Sequence: (1) K0-CyclinB1-NT: 5’ ggatccatgtatccatatgatgttccagattatgctctcgcgctccgagtcaccaggaactcgcgcattaatgctgaaaatcgggcgcgcatcaacatggcaggcgcacgtcgcgttcctacggcccctgctgcaacctcccgccccggactgaggccaagaacagctcttggggacattggtaaccgcgtcagtgaacaactgcaggcccggatgcctatgcgcagagaagcacgaccttcagctactggacgcgtcattgatcggcgcctaccacgccctcttgaaagagtacctctctgaaagctt 3’ To knock down PSMD2 in HEK293T cells we used doxycycline inducible pTRIPZ-shRNA_PSMD2 construct (a gift from Peter Tsvetkov, Broad Institute).

### Cell Culture, Growth conditions, Transfections and reagents

HEK293T and HeLa cells were grown in HiGlucose DMEM media (*Sigma*) supplemented with 10% FBS (*Biological Industries*) and antibiotics in 5% CO_2_ incubator at 37°C. For hypoxia treatment cells were grown in above media in hypoxia chamber (*Baker Ruskinn*) with 1% O_2_ and 5% CO_2_ at 37°C. All transient transfection was performed using the X-tremeGENE HP transfection reagent (*Roche*). To generate doxycycline inducible PSMD2 knock-down clones in HEK293T cells, linearized pTRIPZ-shRNA_PSMD2 (From 5’LTR to 3’LTR) was transfected followed by clone selection under puromycin at 1 µg/mL concentration. High RFP expressing, Puromycin resistant cells were selected by FACS and grown at 0.5 µg/mL puromycin to generate stable clones. For induction of PSMD2 shRNA expression the stable clone cells were grown with doxycycline at 1 ug/mL concentration for two passages. For various experiments different reagents e.g., Bortezomib (1 µM), Chloroquine (50 µM), E1-inhibitor PYR-41 (10 µM), Cyclohexamide (100 µg/mL), aminopeptidase inhibitor CHR 2797 (1 µM) and Puromycin (250 ng/mL to 10 µg/mL) were used.

### Chemical synthesis and ubiquitination of conjugates

#### General Methodology

SPPS was carried out manually in syringes, equipped with teflon filters, purchased from Torviq or by using an automated peptide synthesizer (*CSBIO*). Analytical HPLC was performed on an instrument (*Thermo Scientific*, Spectra System p4000) using an analytical column (Xbridge BEH300 C4 3.5 µm, 4.6 × 150 mm) at flow rate of 1.2 mL/min. Preparative HPLC (Waters) was performed using X Select C18 10µm 19 × 250 mm and semi preparative HPLC (*Thermo Scientific*, Spectra System scm1000) was performed using Jupiter C4 10 µm, 300 Å, 250 × 10 mm column, at flow rate of 15 and 4 mL/min respectively. Commercial reagents were used without further purification: Analytical grade DMF (*Biotech*), Resins (*Creosalus*), protected amino acids (*GL Biochem*), and activating reagents HBTU, HOBt, HCTU, HATU (*Luxembourg Bio Technologies*).

Buffer A: 0.1% TFA in water; Buffer B: 0.1% TFA in acetonitrile.

#### Synthesis of HA-CyclinB1-NT

HA tag = YPYDVPDYA

#### Sequence

NleALRVTRNSRINAENRARINNleAGARRVPTAPAATSRPGLRPRT**A**LGDIGNRVSEQLQARNlePLRK**(64)**EARPSATGRVIDR RLPRPLERVPNle

#### For synthesis of HA-CyclinB1-NT the sequence was divided in to two fragments

1. **C**LGDIGNRVSEQLQARNlePMR**K(64)**EARPSATGRVIDRRLPRPLERVPNle
2. YPYDVPDYA NleALRVTRNSRINAENRARINNleAGARRVPTAPAATSRPGLRPRT-Nbz

#### Synthesis of Differentially tagged TetraUb^K48^-HA-CyclinB1-NT

**Scheme 1:**
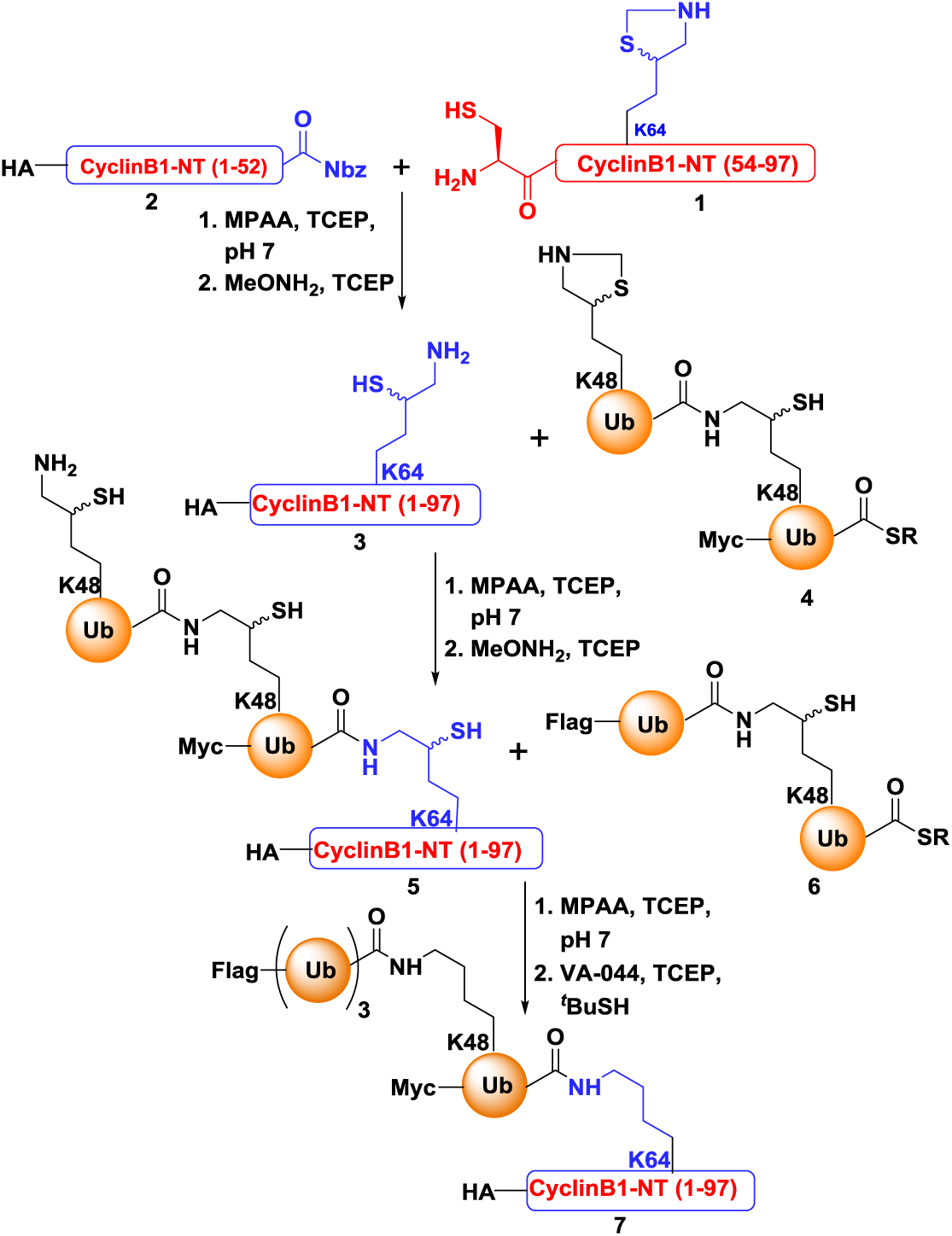
Schematic representation of strategy used for the synthesis of differentially tagged tetraUbK48-HA-CyclinB1-NT, (Nbz = N-acyl-benzimidazolinones, SR = 3-mercaptopropionic acid)

#### Synthesis of fragment 1, CyclinB1-NT (53-97)

CLGDIGNRVSEQLQARNlePMR**K(64)**EARPSATGRVIDRRLPRPLERVPNle

**Scheme 2:**
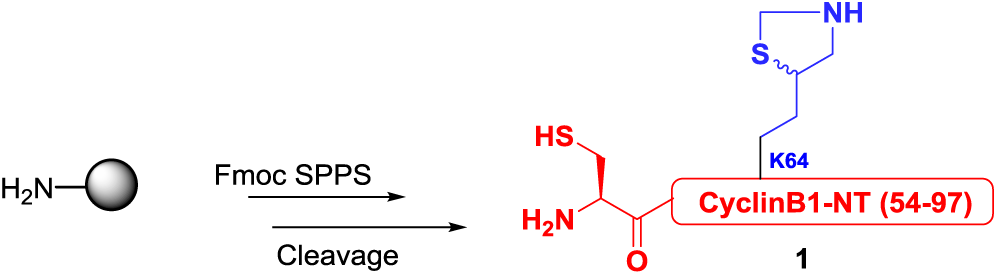
Synthesis of Fragment **1**.

The synthesis was carried out using Fmoc-SPPS on Rink amide resin (0.27 mmol/g, 0.1 mmol scale). Peptide synthesis was performed on peptide synthesizer in presence of 0.4 mmol of amino acid, HCTU) and 0.8 mmol of DIEA. For the synthesis of fragment **1**, the pre-swollen resin was treated with 20% piperidine in DMF containing 0.1 mmol HOBt (3 ×3 cycles) to remove the Fmoc-protecting group. The Thz protected δ-mercaptolysine was manually coupled for 1.5 h at the position Lysine64 using HATU/DIEA. The remaining amino acids were coupled on an automated peptide synthesizer.

#### Cleavage from the resin

The resin was washed with DMF, MeOH, DCM and dried. The peptide was cleaved using TFA:triisopropylsilane (TIS):water (95:2.5:2.5) cocktail for 2 h. The cleavage mixture was filtered and the combined filtrate was added drop-wise to a 10 fold volume of cold ether and centrifuged. The precipitated crude peptide was dissolved in acetonitrile-water (1:1) and was further diluted to ∼30% with water and lyophilized. The HPLC analysis was carried out on a C18 analytical column using a gradient of 0-60% B over 30 min. For preparative HPLC, the same gradient was used to purify the fragment **1** in ∼50% yield.

**Figure M1:**
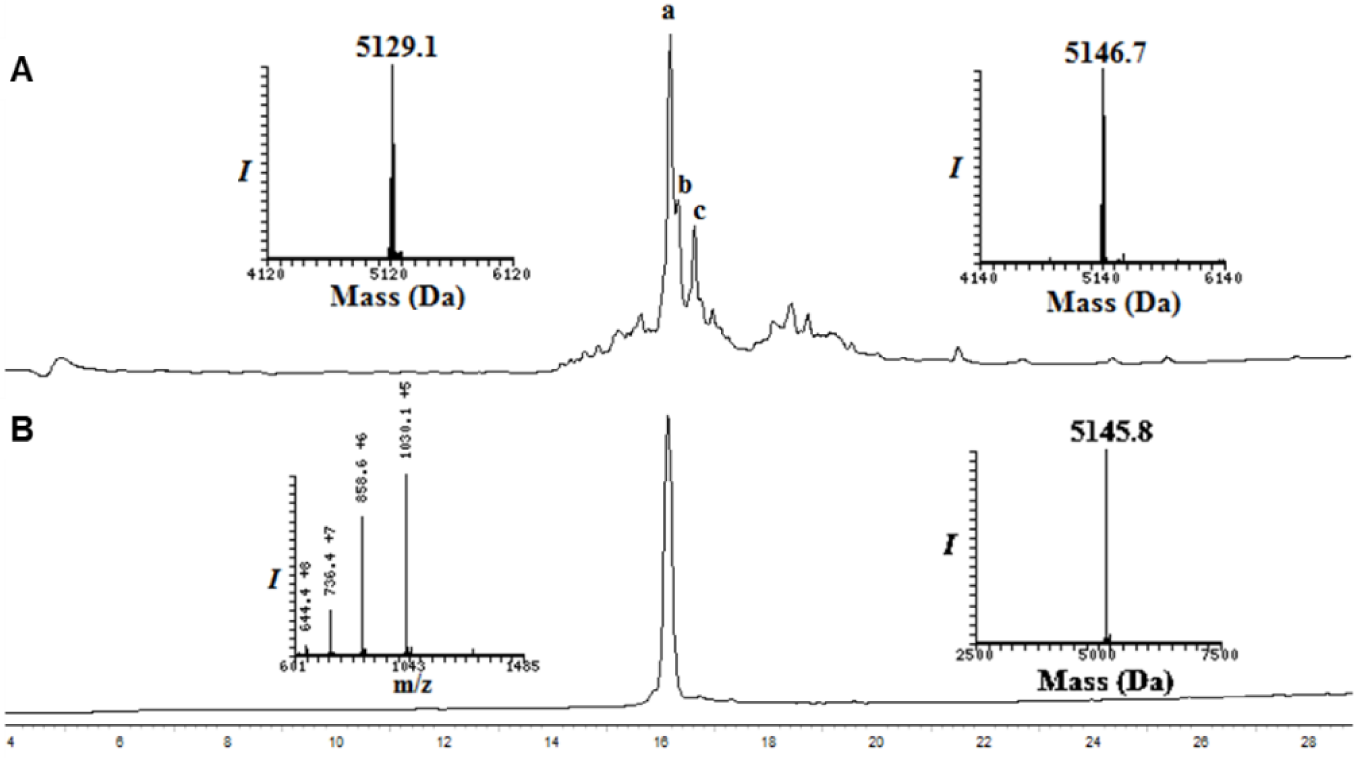
Analytical HPLC and mass traces of fragment-1, **CyclinB1-NT (53-97)**. (**A**) HPLC analysis of crude fragment **1**. Peak **a** corresponds to fragment **1** with the observed mass 5146.7Da (calcd 5146.9 Da); Peak **b** corresponds to a deletion - 18 from the parent peptide. (**B**) Purified fragment **1**.

### Synthesis of fragment 2, HA-CyclinB1-NT (1-52)-Nbz

YPYDVPDYA NleALRVTRNSRINAENRARINNleAGARRVPTAPAATSRPGLRPRT-Nbz

**Scheme 3:**
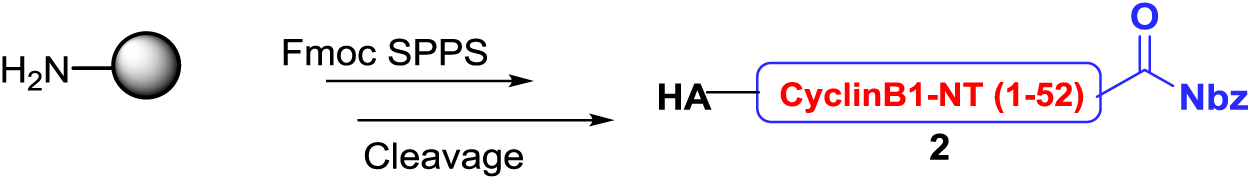
Synthesis of Fragment **2**, (Nbz = N-acyl-benzimidazolinones)

The synthesis was carried out using the N-acylurea method on Rink amide resin (0.27 mmol/g, 0.1 mmol scale). Mono-Fmoc-3,4-diaminobenzoic acid (Fmoc-Dbz) activated with HBTU/HOBt was coupled to the resin for 1 h (2 cycles). Subsequently, the resin was washed with DMF and DCM (3 x 5 mL) and was treated with a solution of allyl chloroformate (50 equivalents) and DIEA (0.2 mmol) in DCM for 12 h. Following the Fmoc removal Peptide synthesis was performed on peptide synthesizer in presence of 0.4 mmol of amino acid and coupling agent (HCTU) while 0.8 mmol of DIEA. The remaining amino acids were coupled using peptide synthesizer as described above. After completion of peptide synthesis, The Alloc-protected resin was washed and swollen in DCM. A solution of PhSiH_3_ 20 equivalents with respect to initial loading of the resin and Pd (PPh3) 4 (0.35 eq) in DCM was added to the resin and shaken for 1 h at 25°C to remove alloc protection. The resin was washed with DCM and a solution of *p*-nitrophenylchloroformate (100 mg, 5 equivalents) in 5 mL of DCM was added and shaken for 1 h at RT. The resin was washed with DCM (3 × 5 mL). Following these steps, a solution of 0.5 M DIEA in DMF (5 mL) was added and shaken for additional 30 min to complete the cyclization. The resin was washed using DMF (3 × 5 mL). Cleavage and purification were carried out as described above to afford the fragment **2** in 40-50% yield.

**Figure M2:**
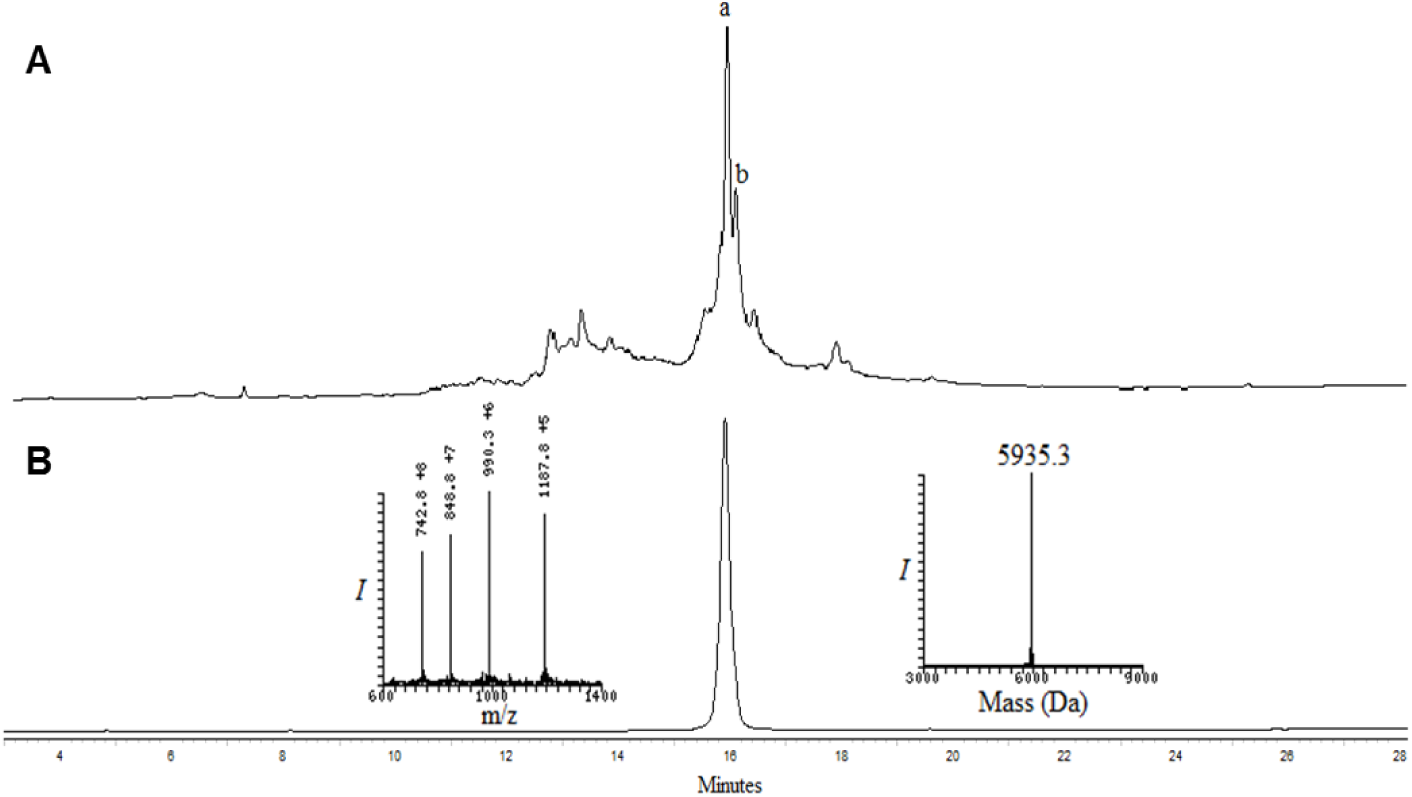
Analytical HPLC and mass traces of fragment 2, **HA-CyclinB1-NT (1-52)-Nbz**. (**A**) HPLC analysis of crude fragment **2**. Peak a corresponds to fragment **2** with the observed mass 5935.3 Da (calculated 5935.5 Da); Peak **b** corresponds to a deletion from the parent peptide. (**B**) Purified fragment **1**.

### Native chemical ligation between fragment 1, CyclinB1-NT (53-97) and fragment 2, HA-CyclinB1-NT (1-52)-Nbz

#### Ligation

fragment **1**, CyclinB1-NT (53-97) (10 mg, 1.9 × 10^-3^ mmol) and fragment **2**, HA-CyclinB1-NT (1-52)-Nbz (13.8 mg, 2.3 × 10^-3^ mmol) were dissolved in argon purged 6 M Gn·HCl, 200 mM Na_2_HPO_4_ buffer (971 µL, 2 mM) containing 20 equivalents of MPAA and 10 equivalents of TCEP, pH ∼7.3. The reaction was incubated at 37°C for 4 h. Progress of the reaction was monitored by analytical HPLC using C18 column with a gradient of 5-55% buffer B over 45 min. After completion of ligation, the reaction mixture was diluted to 1mM concentration with a solution of 30 equivalents of methoxylamine and 15 equivalents of TCEP at pH = 4. The reaction mixture was incubated in 37°C for 12 h. The progress of the reaction was monitored by analytical HPLC using C18 column with a gradient of 5-55% B over 45min and LC-MS analysis. For semi-preparative HPLC, the same gradient was used to isolate the ligation product **3** in 63% yield (∼13 mg).

**Scheme 4:**
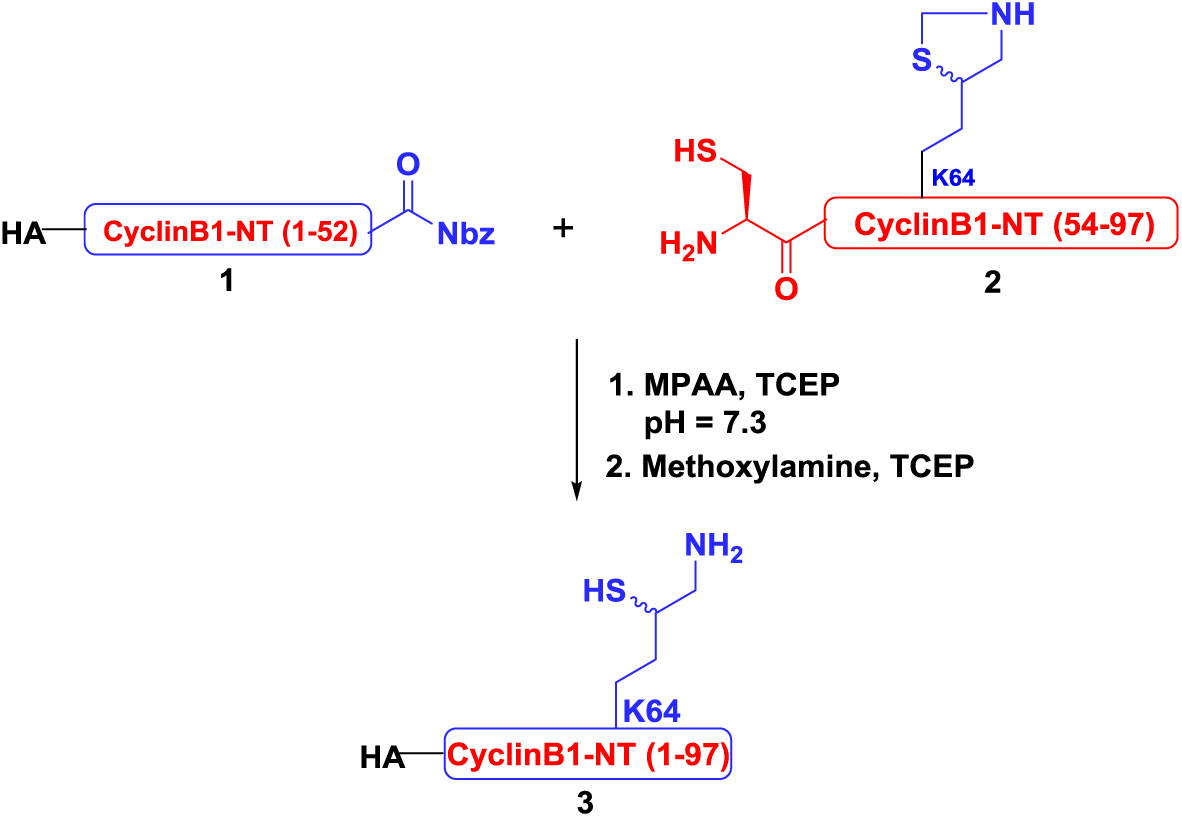
Synthesis of HA-CyclinB1-NT (1-97), **3**.

**Figure M3:**
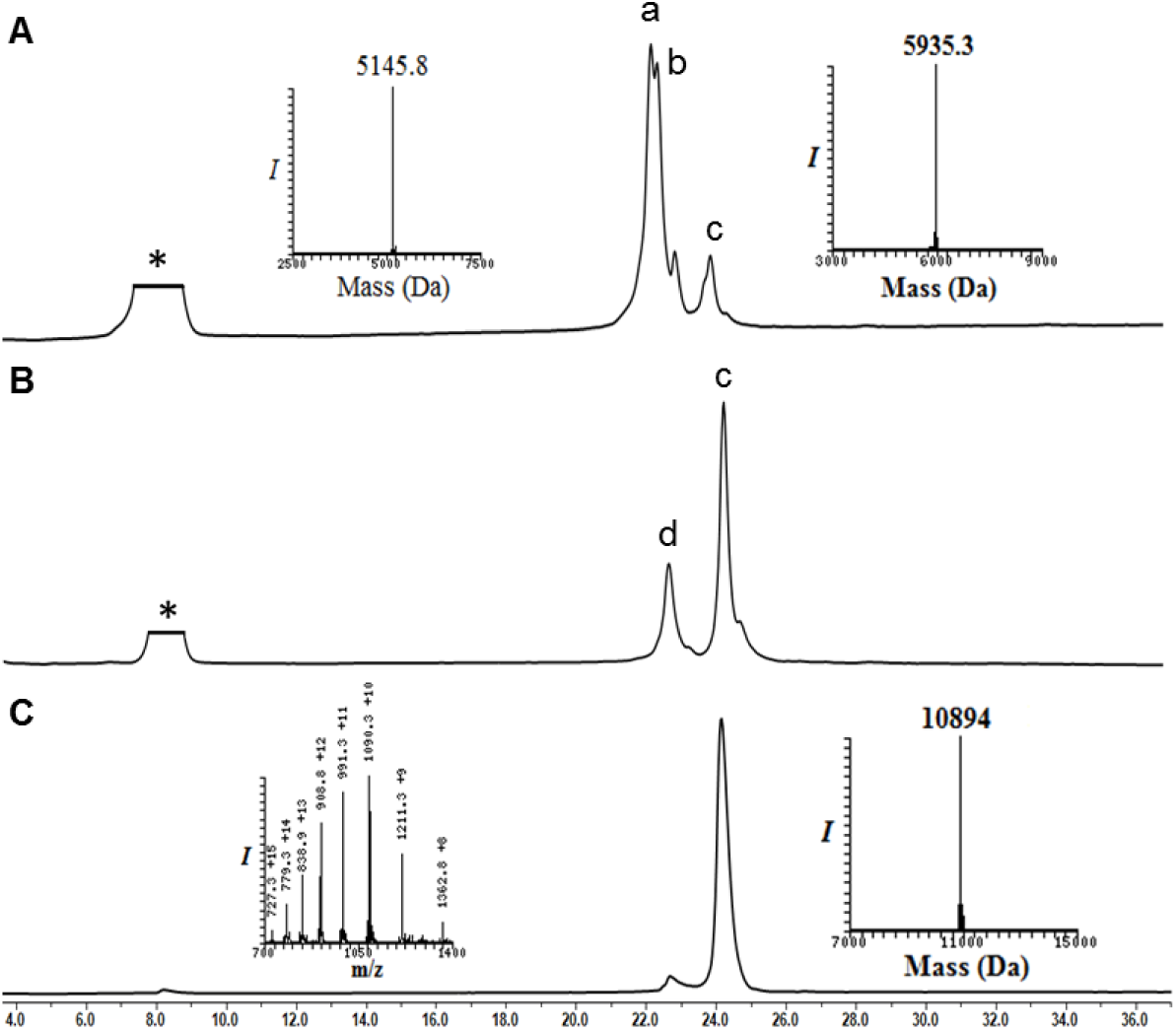
Analytical HPLC and mass traces for the ligation of fragment 1 and fragment 2. (**A**) Ligation at time zero, peak **a** corresponds to fragment **1**, peak **b** corresponds to fragment **2**, peak **c** corresponds to ligation product **3**. (**B**) Ligation after 6 h, peak **c** corresponds to ligation product **3** with the observed mass 10894Da (calculated 10893 Da), peak **d** corresponds to Nbz hydrolyzed fragment **2**. (**C**) Purified HA-CyclinB1-NT, **3**.

### Desulfurization of HA-CyclinB1-NT

**3** was subjected for the desulfurization using VA-044 (20 equivalents, 0.1 M), TCEP (0.25 M) in the presence of 10% t-BuSH (v/v) at 37°C for 6 h. The reaction was followed using analytical HPLC (C18 column) and a gradient of 5-55% B over 60 min and LC-MS analysis. For semi-preparative HPLC, the same gradient was used to afford the desulfurized HA-CyclinB1-NT ∼60% yield

**Scheme 5:**
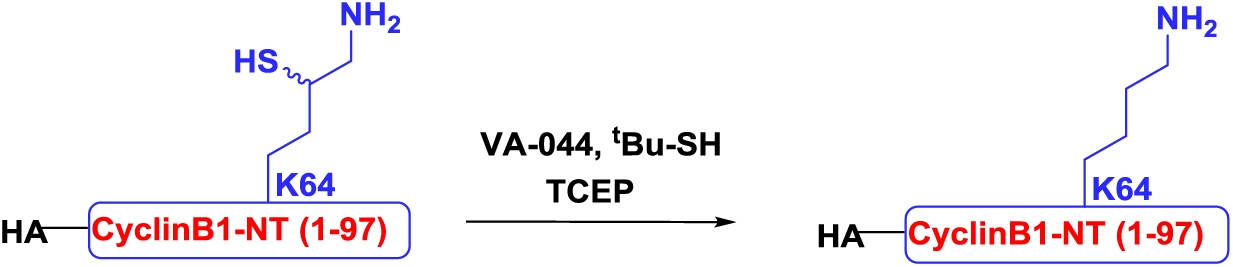
Desulfurization of HA-CyclinB1-NT-(1-97)

**Figure M4:**
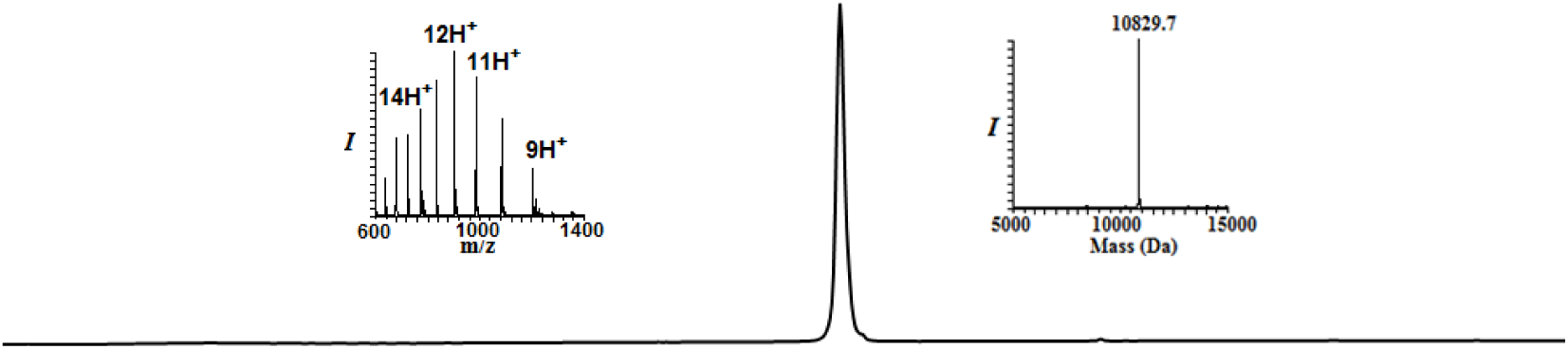
Analytical HPLC and mass traces for the desulfurized and purified HA-CyclinB1-NT with the observed mass 10829.7Da (calculated 10829.5Da)

### Synthesis of Ubiquitin Monomers

The synthesis of Myc-Ub1-N-Methyl-Cysteine, Ub2/3(K48*)-N-Methyl-Cysteine and Flag-Ub4-N-Me-Cysteine were carried on Rink amide resin (0.27 mmol/g, 0.1 mmol scale). The pre-swollen resin was treated with 20% piperidine in DMF containing 0.1 mmol HOBt (3 cycle 3 min each) to remove resin bound Fmoc-protecting group. Initially, Fmoc-S-*o*-nitrobenzyl N-methyl cysteine activated with HATU/DIEA was coupled to the resin for 1.5 h. Subsequently, the resin was washed with DMF. Following the Fmoc removal, all amino acids were coupled using an automated peptide synthesizer in presence of 0.4 mmol of amino acid, 0.8 mmol of DIEA and 0.4 mmol of coupling agent (HCTU) to the initial loading of the resin. Pseudoproline dipeptides Leu-Ser, Asp-Gly(Dmb), Ile-Thr and Leu-Thr were manually coupled at positions Leu56-Ser57, Asp52-Gly53, Ile13-Thr14 and Leu8-Thr9 respectively using 0.25 mmol of Fmoc-Leu-Ser(ΨMe, Mepro)-OH, Fmoc-Asp(OtBu)-(Dmb)Gly-OH, Fmoc-Ile-Thr(ΨMe, Mepro)-OH and Fmoc-Leu-Thr(ΨMe, Mepro)-OH respectively. During synthesis of Myc-Ub1-N-Me-Cysteine, Ub2/3(K48*)-N-Me-Cysteine 0.15 mmol thiazolidine (Thz)-protected δ-mercaptolysine was manually coupled for 2 h at the position 48 using HATU/DIEA. Analytical cleavage and HPLC analysis were performed after these couplings to ensure complete reaction. Norleucine was used in place of Met1 in ubiquitin sequence to avoid oxidation during synthesis. After the completion of synthesis, ubiquitin analogues were cleaved from resin as follow.

Cleavage from resin: The resin was washed with DMF, DCM and dried over high vacuum. A cocktail of TFA:DCM:triisopropylsilane:water (90:5:2.5:2.5) was added to resin and reaction mixture was shaken for 2 h at RT. The resin was filtered, and the combined filtrate was added dropwise to a 10 fold volume of cold ether and centrifuged. The precipitated crude peptide was dissolved (∼70%) in acetonitrilewater (1:1) and was further diluted to ∼30% with water and lyophilized. Myc tag-EQKLISEEDL and Flag tag-DYKDDDDK

### Thz deprotection of Myc-Ub1K48

The crude peptide (50 mg) was dissolved in 6M Gn·HCl buffer to a final concentration of ∼3 mM. Hydrazine hydrate (80% solution, 100 equivalents) was added and kept at 25°C for 1 h (pH ∼7). To the above reaction mixture, methoxylamine (0.2 M, 15 equivalents) and tris-(2-carboxyethyl)-phosphine (TCEP, 30 equivalents) were added and incubated at 37°C for 3 h to unmask the δ-mercaptolysine completely. The reaction was monitored using an analytical HPLC (C18 column) and a gradient of 0-60% B over 30 min and LC-MS analysis. For HPLC purification, a similar gradient was used to afford the corresponding peptide in ∼7% yield.

**Scheme 6:**
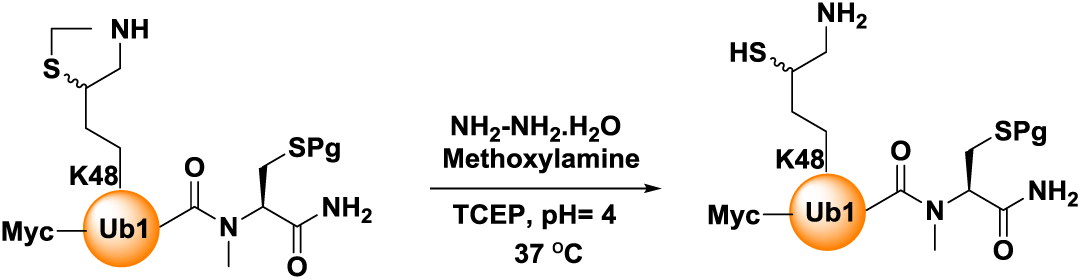
Synthesis of Myc-Ub1 building block.

**Figure M5.**
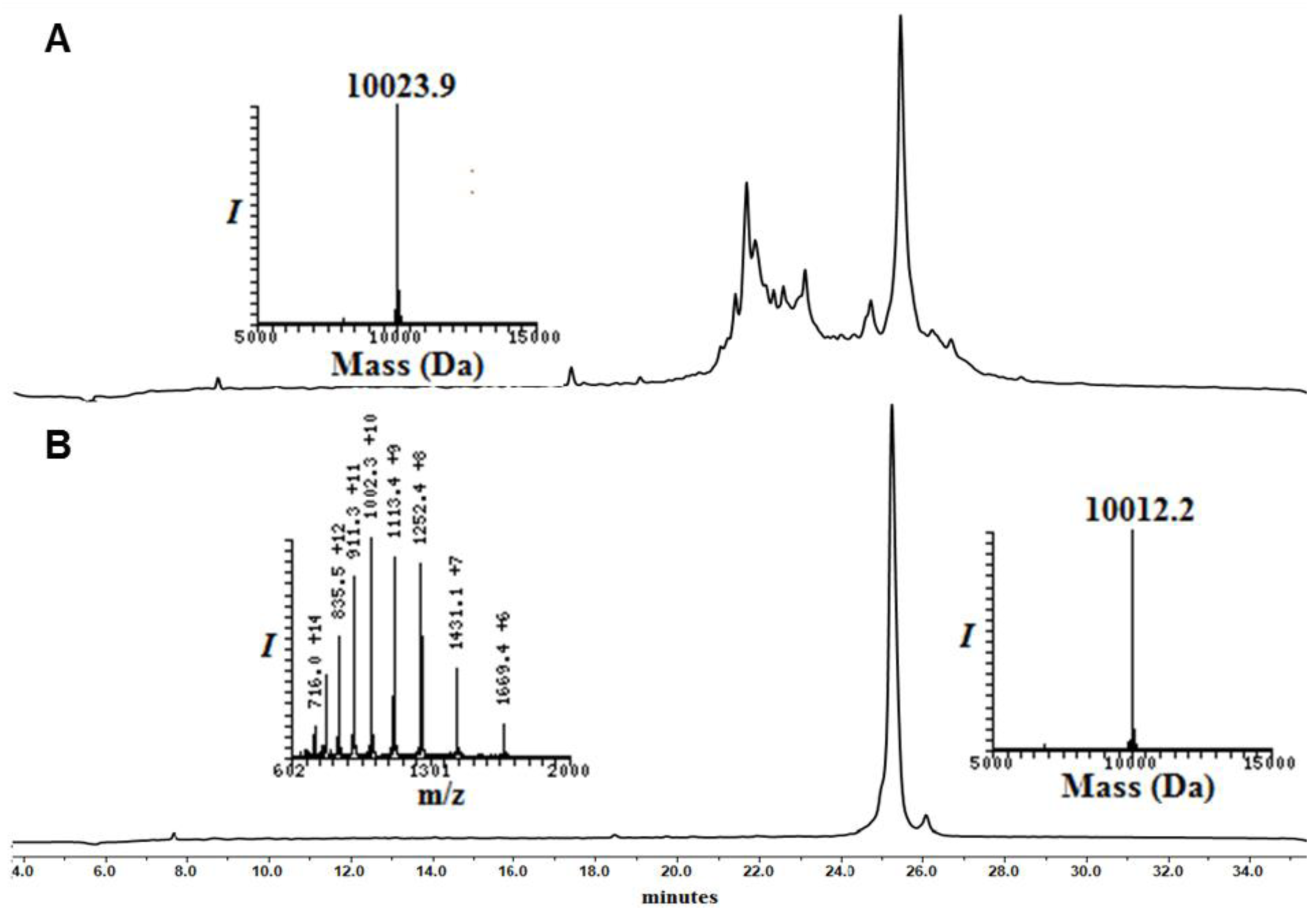
Analytical HPLC and mass traces of the synthesis of Myc-Ub1 building block with δ–mercaptolysine at K48: A) Crude peptide of Myc-Ub1; peak **a** corresponds to the desired product with the observed mass of 10023.9 Da (calculated 10027.1 Da). (**B**) Purified thz deprotected Myc-Ub1 after methoxylamine treatment with the observed mass 10012.2 Da (calculated 10015.1 Da);

#### Synthesis of Ub2 and Ub3

a. **Conversion of Ub2K48-N-methyl cysteine to 3-mercaptopropionic acid ester** Ub2K48-N-methyl cysteine (50 mg) was dissolved in 6 M Gn·HCl buffer (2.82 mL, 2 mM) and was subjected for the UV irradiation (365 nm, 2 h) to remove photolabile 2-nitrobenzyl protecting group of the C-terminal Cys, followed by the pH adjustment (∼1.5-2) and incubation with 20% (vol/vol) 3-mercaptopropionic acid (MPA) at 42°C for 20 h. The reaction was followed by HPLC using a C18 analytical column and a gradient of 0-60% B over 30 min and LC-MS analysis. For preparative HPLC, the same gradient was used to afford the Ub2K48-MPA thioester at a yield of ∼10% (∼4.9 mg).
b. **Thz deprotection of Ub3K48-N-methyl cysteine**

Similar procedure used for the thz deprotection of Myc-UbK48-N-Methyl cysteine was used for thz deprotection of Ub3K48-N-methyl cysteine to yield the product in 9% yield (∼4.5 mg).

**Scheme 7:**
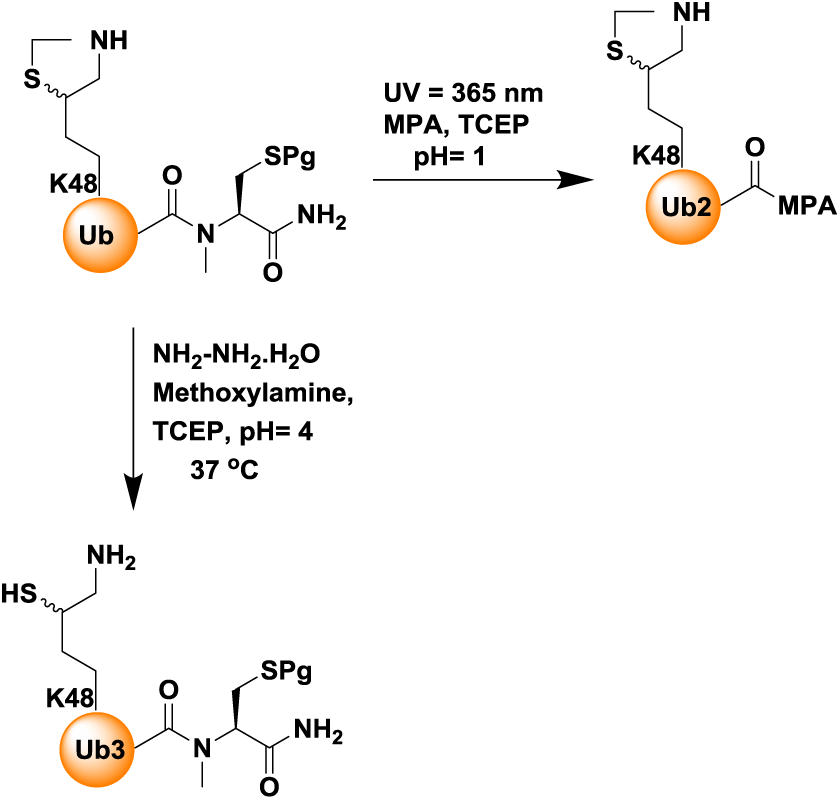
Synthesis of Ub2 and Ub3 building blocks, (Pg = O-Nitro Benzyl, MPA = 3-mercaptopropionic acid)

**Figure M6.**
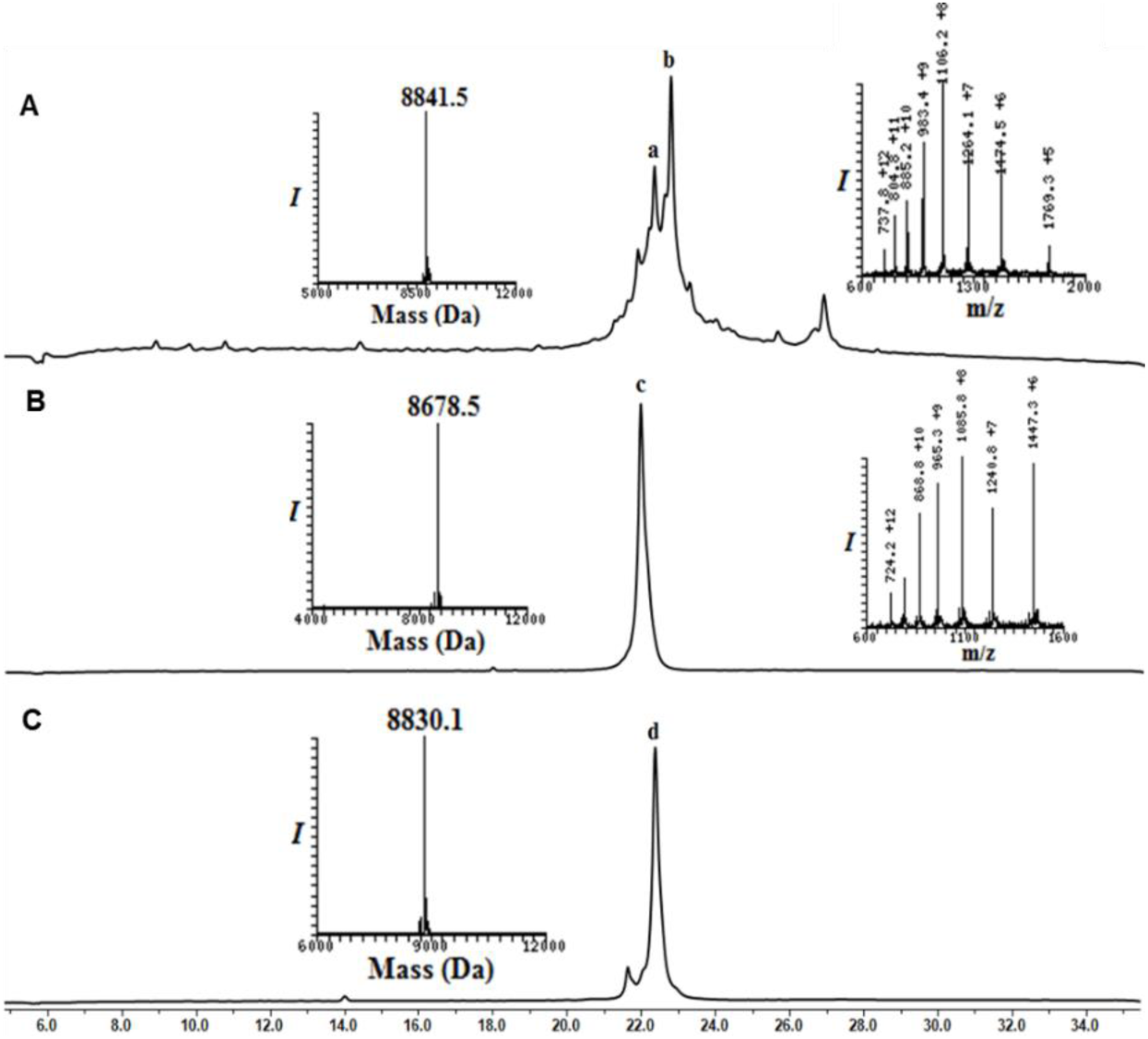
Analytical HPLC and mass traces of the synthesis of Ub2 and Ub3 building block with δ–mercaptolysine at K48: (**A**) Crude peptide of UbK48-N-Methyl cysteine; peak **b** corresponds to the desired product with the observed mass of 8841.5 Da (calculated 8,841.8 Da) peak **a** corresponds to deletion of arginine from Ub sequence. (**B**) Purified Ub2-MPA thioester after N-methyl cysteine switching to MPA thioester with the observed mass 8678.5 Da (calculated 8,678.8 Da). (**C**) Purified thz deprotected Ub3 after methoxylamine treatment with observed mass 8830.1 (calculated 8,830.8 Da).

#### Synthesis of Flag-Ub

DYKDDDDKNleQIFVKTLTGKTITLEVEPSDTIENVKAKIQDKEGIPPDQQRLIFAGKQLEDGRTLSDYNIQKESTLHLVLRLRGG-*N*-Methyl Cysteine

**Scheme 8:**
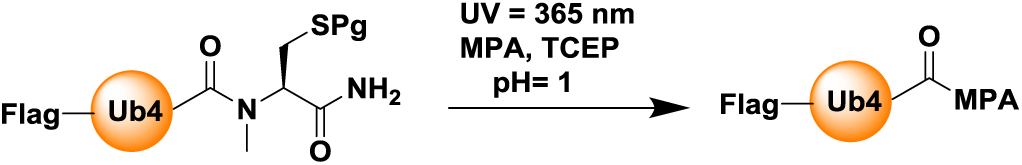
Synthesis of Flag-Ub4-MPA, (Pg = O-Nitro Benzyl, MPA = 3-mercaptopropionic acid)

**Figure M7.**
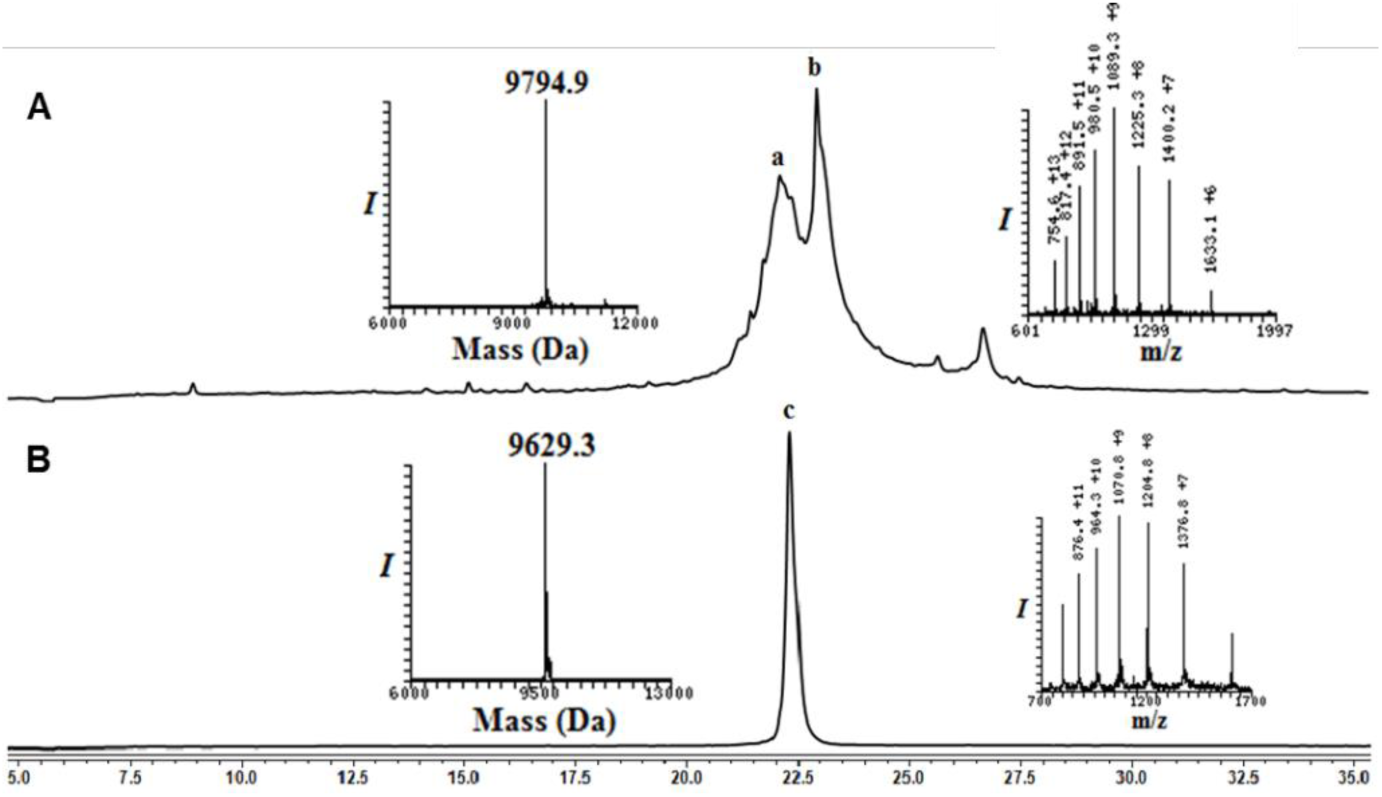
Analytical HPLC and mass traces of the synthesis of Flag-Ub4-MPA building block: (**A**) Crude Flag-Ub4-N-Methyl cysteine; peak b corresponds to the desired product with the observed mass of 9794.9 Da (calculated 9792.7 Da) peak a with unresolved mass; (**B**) Purified Flag-Ub4-MPA thioester after N-methyl cysteine switching to MPA thioester with the observed mass 9629.3 Da (calculated 9,630.7 Da)

#### Synthesis of MycUb-HA-CyclinB1-NT

Ligation of HA-CyclinB1-NT and MycUb1-MPA was carried out similar fashion as mentioned for conjugate **3** and product was subjected for the desulfurization to yield 40% of MycUb-HA-CyclinB1-NT.

**Scheme 9:**
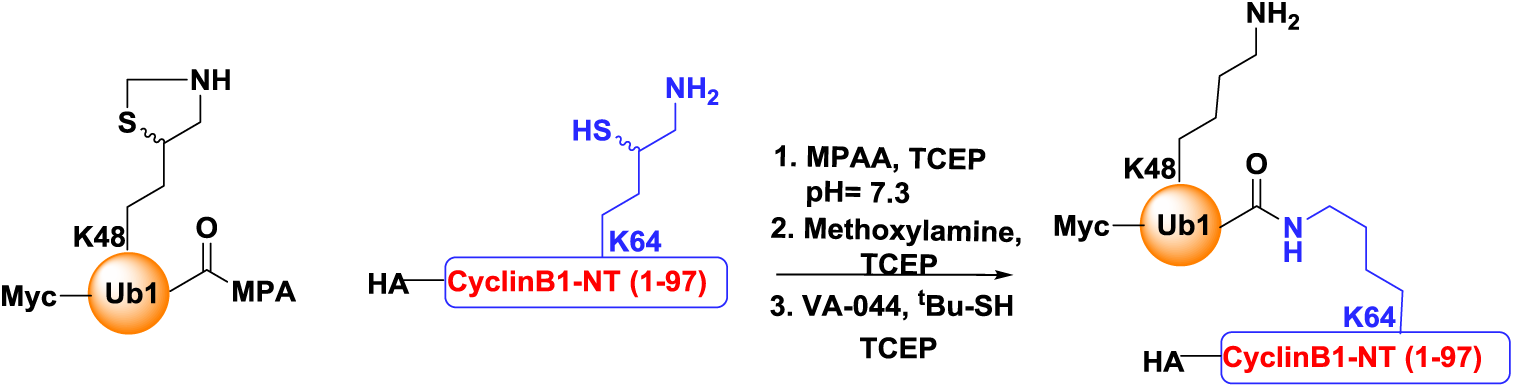
Synthesis of Myc-Ub-HA-CyclicB1-NT.

**Figure M8:**
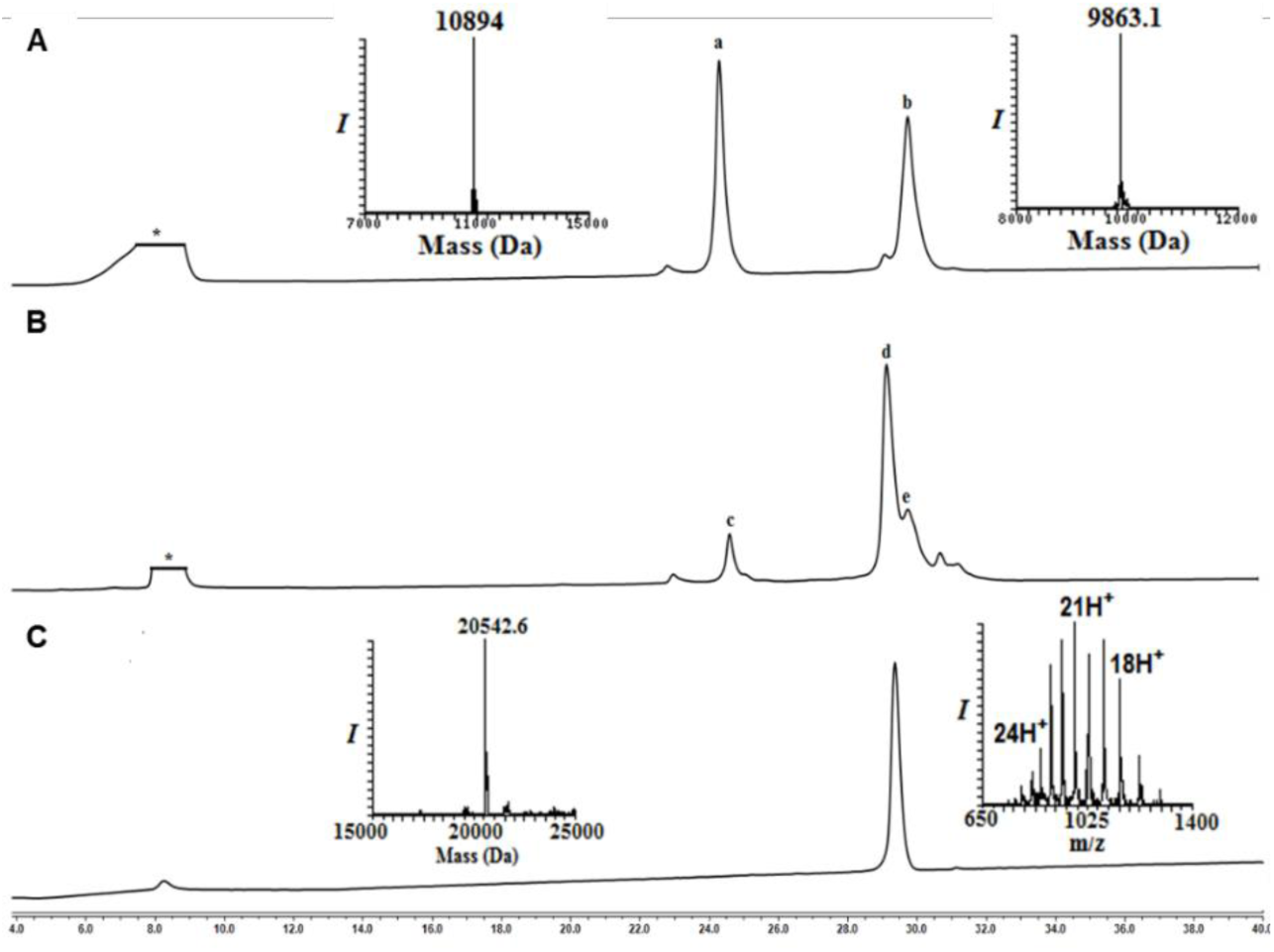
Analytical HPLC and mass traces for the ligation of HA-CyclinB1-NT (ThzOpen) and MycUbK48-MPA. (**A**) Ligation at time zero, peak **a** corresponds to HA-CyclinB1-NT (ThzOpen), peak **b** corresponds to MycUbK48-MPA. (**B**) Ligation after 6 h, peak **c** corresponds to remaining of HA-CyclinB1-NT (ThzOpen), peak **d** corresponds to the ligation product, peak **e** corresponds to the hydrolyzed MycUbK48-MPA. (**C**) Desulfurized and Purified MycUb-HA-CyclinB1-NT with observed mass 20542.6 Da (Calculated 20543.6 Da).

#### Synthesis of Myc-DiUb^K48^-MPA thioester, 4

Ligation: Thz deprotected MycUb1(K48*)-N-Me-Cysteine (10 mg, 1 mmol) and Ub2(K48)-MPA (10.4 mg, 1.2 mmol), were dissolved in 6 M Gn·HCl buffer (500 µL, 2 mM). To this solution, 20 equivalents each of MPAA and TCEP were added, the pH was adjusted to 7.3 and kept at room temperature for 30min and then at 37 °C for 4 h. The reaction was followed using analytical HPLC (C4 column) and a gradient of 5-55% B over 60 min. For preparative HPLC, the same gradient was used to isolate the ligation product in 52% yield (∼9.7 mg). The conversion of MycDiUb–N-methyl cysteine to MycDiUb–MPA thioester was achieved by first removing the photolabile 2-nitrobenzyl protecting group of the C terminal Cys (365 nm, 2 h), followed by adjustment of pH ∼1 and incubation with 20% (vol/vol) MPA at 42 °C for 20 h. The reaction was followed by HPLC using a C4 analytical column and a gradient of 5–55% B over 60 min and LC-MS analysis. For preparative HPLC, the same gradient was used to afford the Myc-DiUb^K48^-MPA thioester, **4** at a yield of ∼38% (∼3.6 mg).

**Scheme 10:**
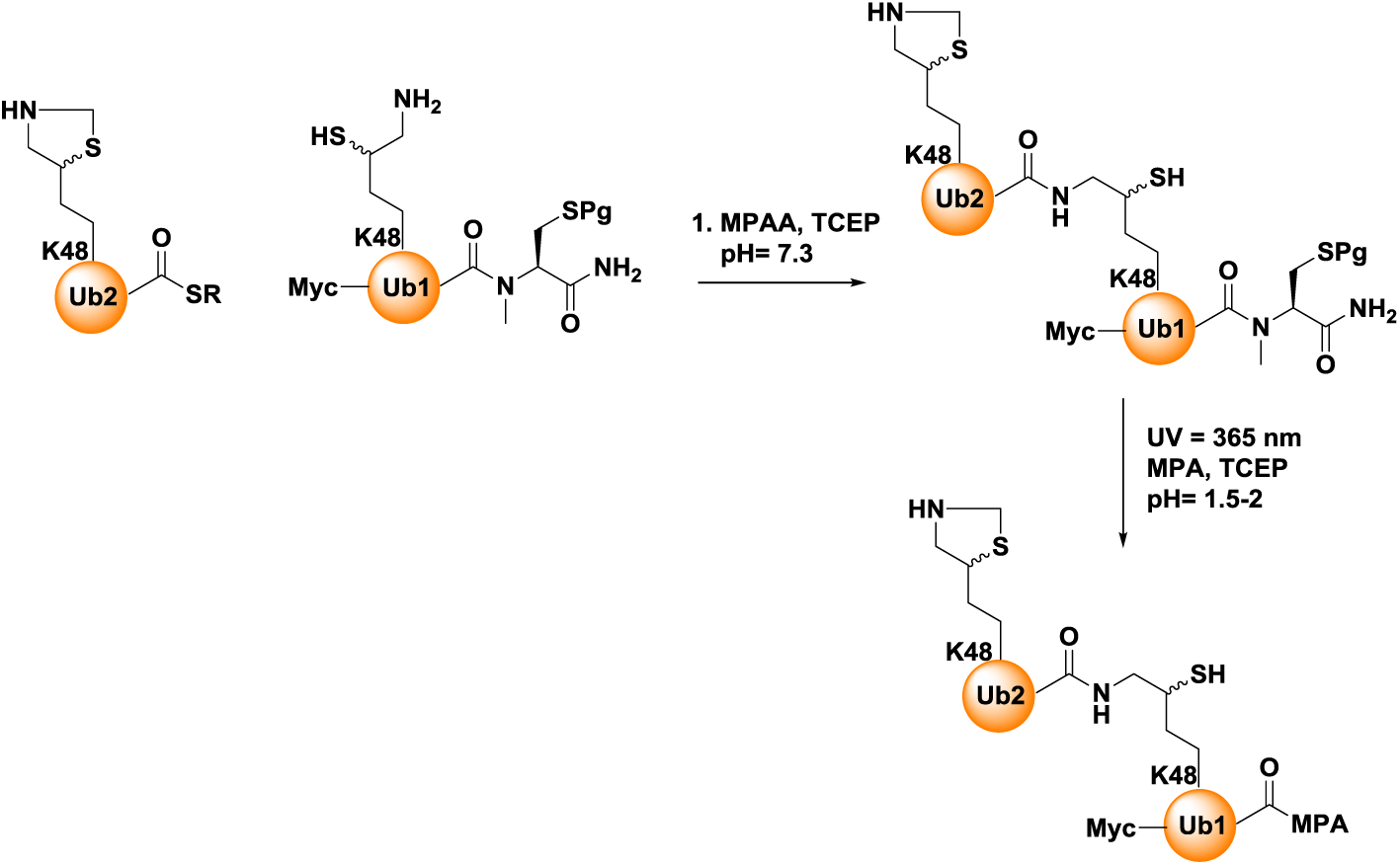
Synthesis of Myc-DiUb-MPA, 4 (Pg = O-Nitro Benzyl, SR = 3-mercaptopropionic acid)

**Figure M9.**
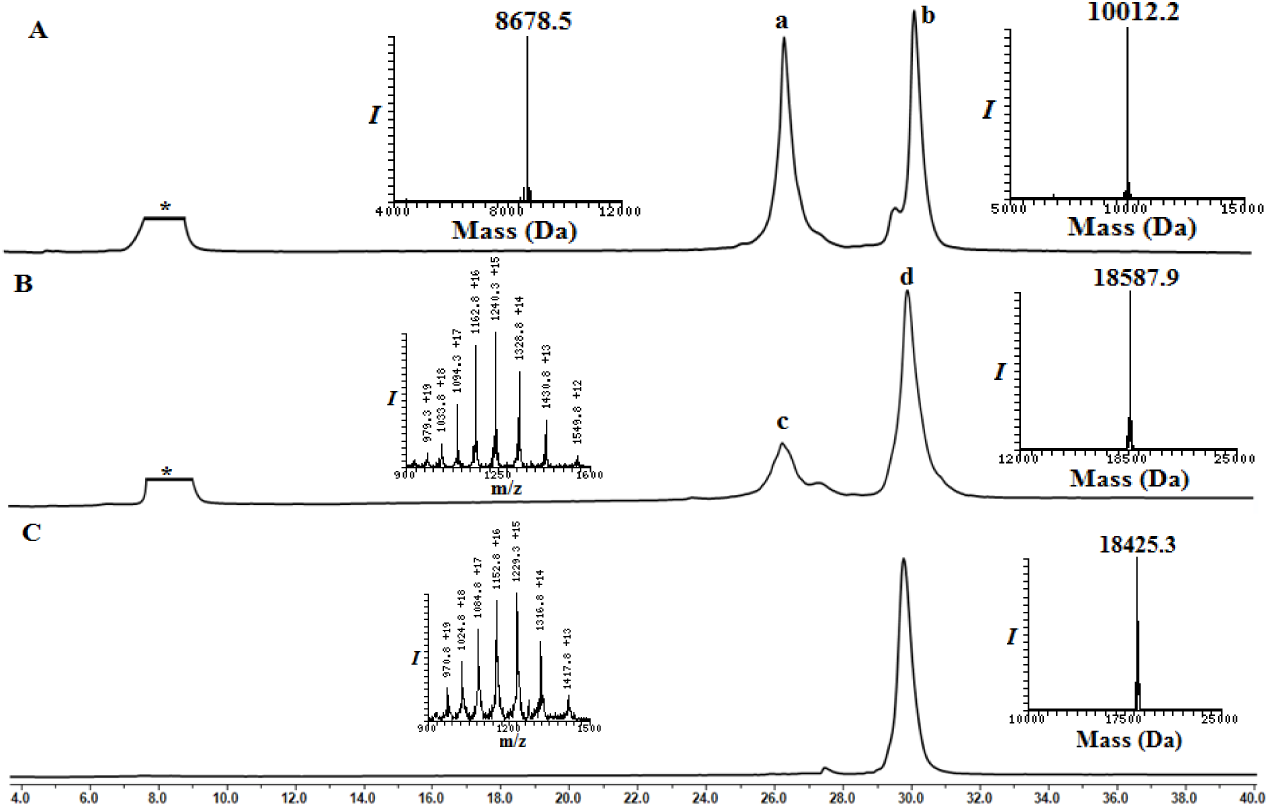
Synthesis of Myc-DiUb^K48^-MPA **4**: (**A**) Analytical HPLC and mass traces of the ligation at 0 h, peak **a** corresponds to Ub2K48-MPA, peak **b** corresponds to MycUb1(K48*)-N-methyl cysteine. (**B**) Ligation after 6 h; peak **c** corresponds to the hydrolyzed Ub2K48-MPA thioester, peak **d** corresponds to the ligation product, Myc-DiUb ^**K48**^-N-Me-Cysteine with the observed mass of 18,587.9 Da (calculated 18,588.9 Da). (**C**) Analytical HPLC and mass traces of purified Myc-DiUb^K48^-MPA after N-Me-Cysteine switching to MPA reaction with observed mass 18425.3 Da (calculated 18425.9 Da)

#### Ligation of HA-CyclinB1-NT (ThzOpen), 3 with Myc-DiUb^K48^-MPA, 4

HA-CyclinB1-NT (ThzOpen) (2 mg, 0.18 mmol) and Myc-DiUb^K48^-MPA (3.6 mg, 0.19 mmol), were dissolved in 6 M Gn·HCl buffer (92 µL, 2 mM). To this solution, 20 equivalents each of MPAA and TCEP were added, the pH was adjusted to 7.3 and kept at 37 °C for 6 h. The reaction was followed using analytical HPLC (C4 column) and a gradient of 5-55% B over 60 min. After completion of ligation, Myc-DiUb^K48^-HA-CyclinB1-NT was subjected for the Thz deprotection using hydrazine hydrate (80% solution, 100 equivalents) and kept at 25°C for 1 h (pH ∼7). To the above reaction mixture, methoxylamine (0.2 M, 15 equivalents) and tris-(2-carboxyethyl)-phosphine (TCEP, 30 equivalents) were added and incubated at 37 °C for 12 h to unmask the δ-mercaptolysine completely. The reaction was monitored using an analytical HPLC (C4 column) and a gradient of 5-55% B over 40 min and LC-MS analysis. For HPLC purification, a similar gradient was used to afford thz deprotected Myc-DiUb^K48^-HA-CyclicB1-NT, **5** in ∼40% yield (2.1 mg).

**Scheme 11:**
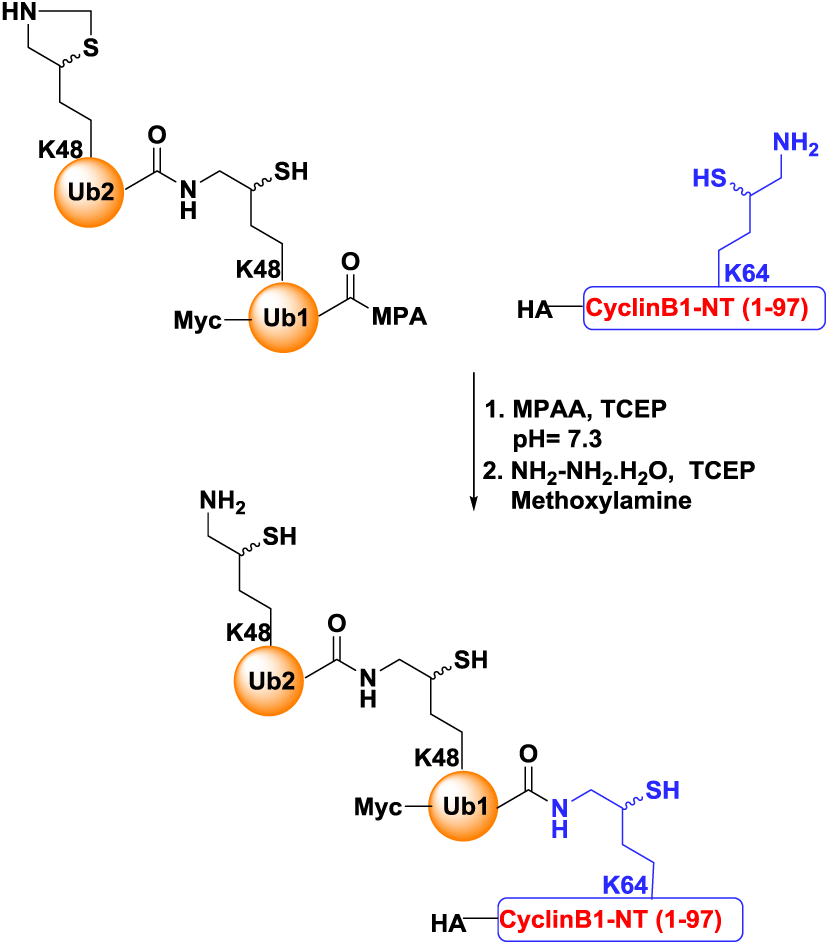
Synthesis of Myc-DiUb^K48^-HA-CyclinB1(NT)

**Figure M10.**
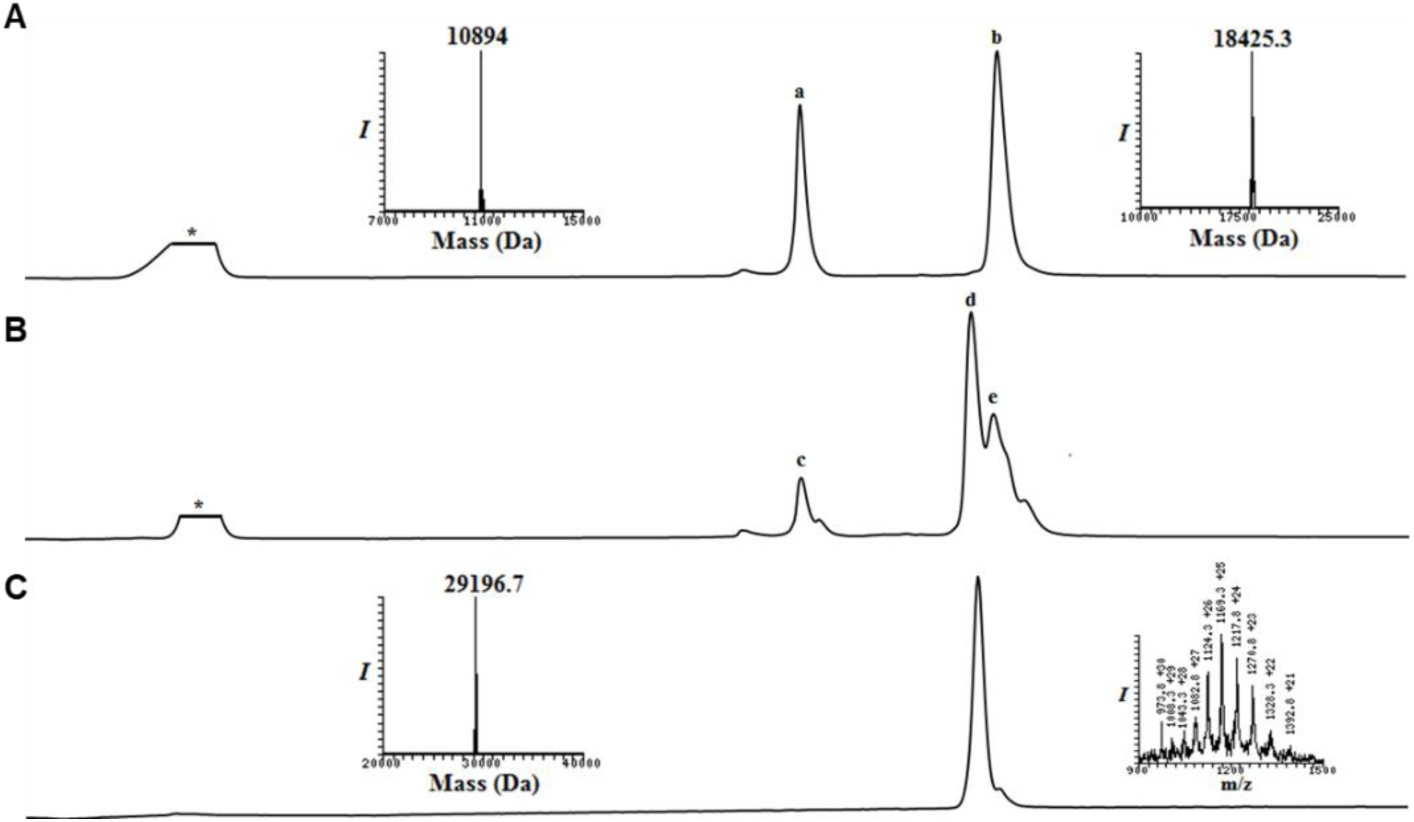
Synthesis of Myc-DiUb^K48^-HA-CyclinB1-NT: (**A**) Analytical HPLC and mass traces of the ligation at 0 h, peak **a** corresponds to HA-CyclinB1-NT, peak **b** corresponds to Myc-DiUb^K48^-MPA. (**B**) Ligation after 6 h; peak **c** corresponds to the remaining HA-CyclinB1-NT, peak **d** corresponds to the ligation product, peak **e** corresponds to hydrolyzed Myc-DiUb^K48^-MPA. (**C**) Analytical HPLC and mass traces of Thz deprotected and purified Myc-DiUb^K48^-HA-CyclinB1-NT with the observed mass of 29,196.7 Da (calculated 29197.3 Da).

#### Desulfurization of Myc-DiUb^K48^-HA-CyclinB1-NT

Desulfurization of Myc-DiUb^K48^-HA-CyclinB1-NT was carried out in similar fashion as mentioned for the conjugate **3**, to yield 35% of Myc-DiUb^K48^-HA-CyclinB1-NT.

**Figure M11:**
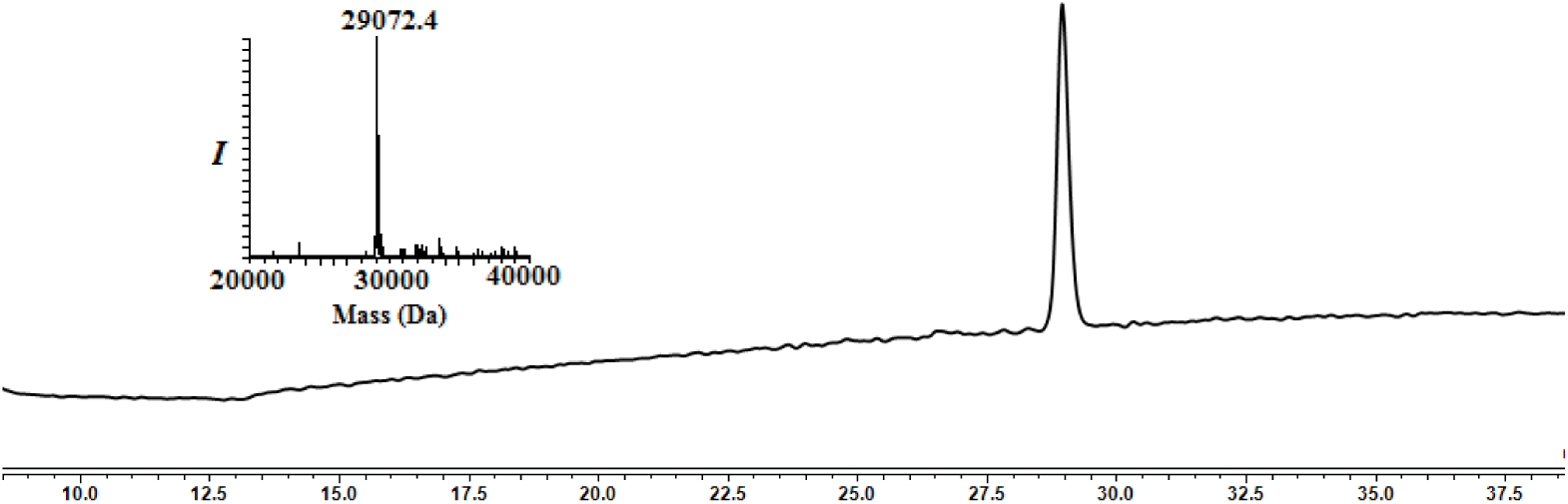
Analytical HPLC and mass traces for the desulfurized and purified Myc-DiUb^48^-HA-CyclinB1-NT with the observed mass 29072.4 Da (calculated 29072.3 Da)

#### Synthesis of Flag-DiUb^K48^-MPA, 6

Synthesis of Flag-DiUb^K48^-MPA was carried out in similar way as described for the conjugate **4** to yield 35% (3.1mg)

**Scheme 12:**
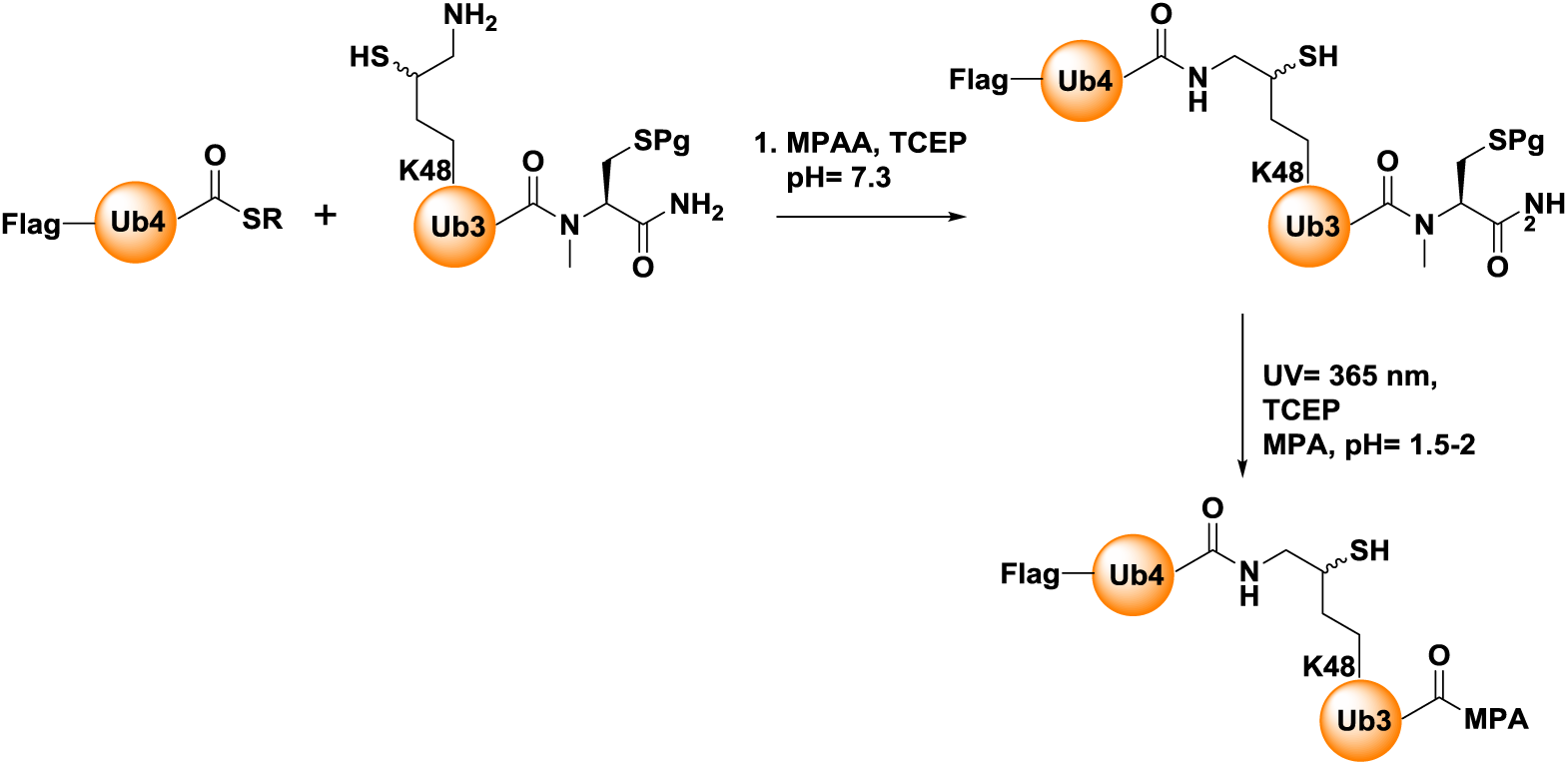
Synthesis of Flag-DiUb^K48^-MPA, **6**.

**Figure M12.**
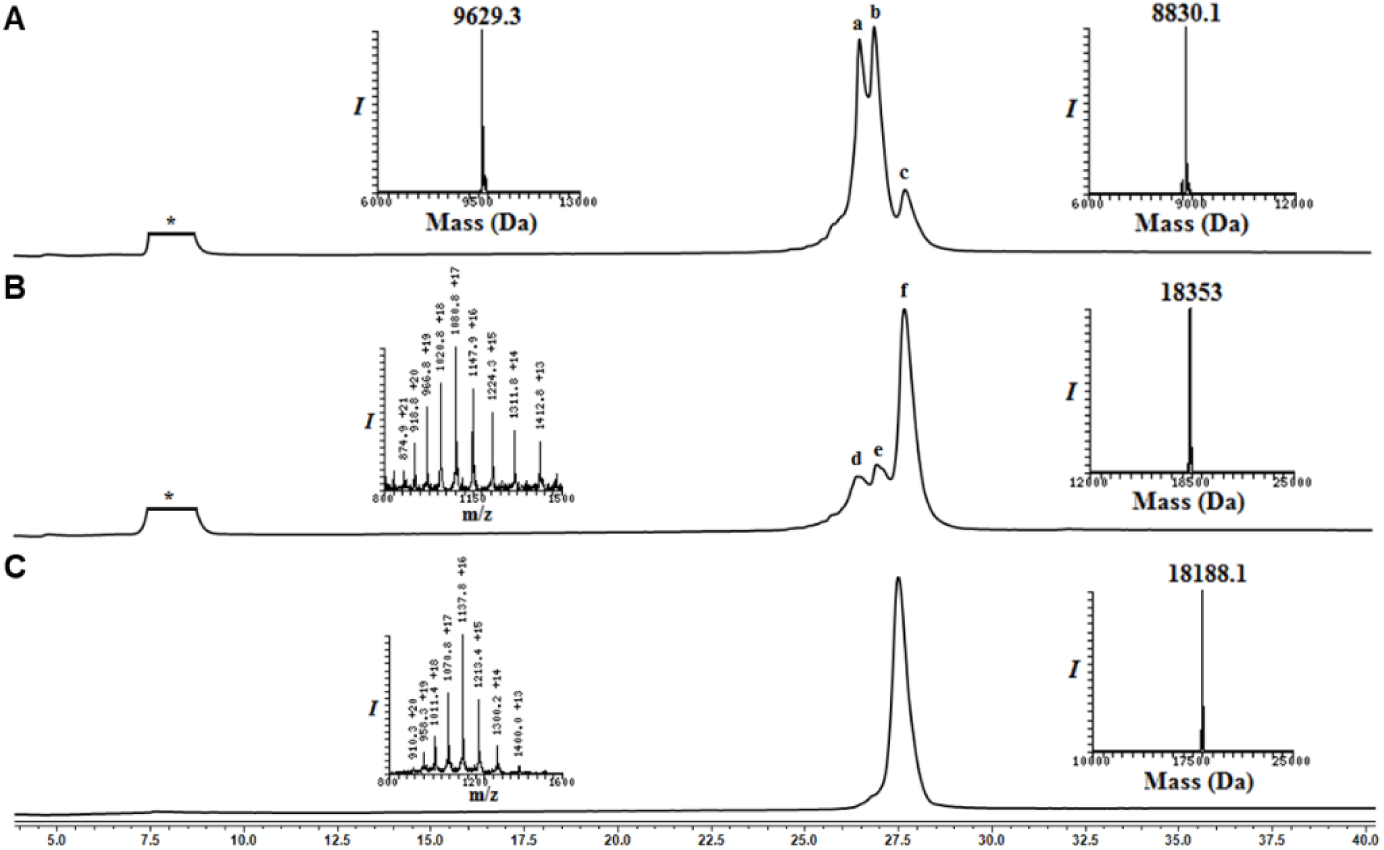
Synthesis of Flag-DiUb^K48^-MPA: (**A**) Analytical HPLC and mass traces of the ligation at 0 h, peak **a** corresponds to Flag-Ub4-MPA, peak **b** corresponds to Ub3(K48*)-N-methyl cysteine, peak **c** corresponds to ligated product. (**B**) Ligation after 6 h; peak **d** corresponds to the hydrolyzed Flag-Ub4-MPA thioester, peak **e** corresponds to remaining Ub3K48-N-methyl cysteine, peak **f** corresponds to the ligation product Flag-DiUb^K48^-N-Methyl cysteine with the observed mass of 18,353 Da (calculated 18,354.5 Da). (**C**) Analytical HPLC and mass traces of purified Flag-DiUb^K48^-MPA with observed mass 18188.1 Da (calculated 18191.5 Da)

#### Ligation of Myc-DiUb^K48^(Thz Open)-HA-CyclinB1-NT, 5 with Flag-DiUb^K48^-MPA, 6

**4** and **7** were ligated in a similar fashion as described for the synthesis of conjugate 8 and subjected for the desulfurization using VA-044 (20 equivalents, 0.1 M), TCEP (0.25 M) in the presence of 10% t-BuSH (v/v) at 37°C for 18 h. The reaction was followed using analytical HPLC (C4 column) and a gradient of 5-55% B over 1hr. For semi-preparative HPLC, the same gradient was used to afford the desulfurized TetraUb^K48^-HA-CyclinB1-NT ∼30% yield (∼1.6 mg).

**Scheme 13:**
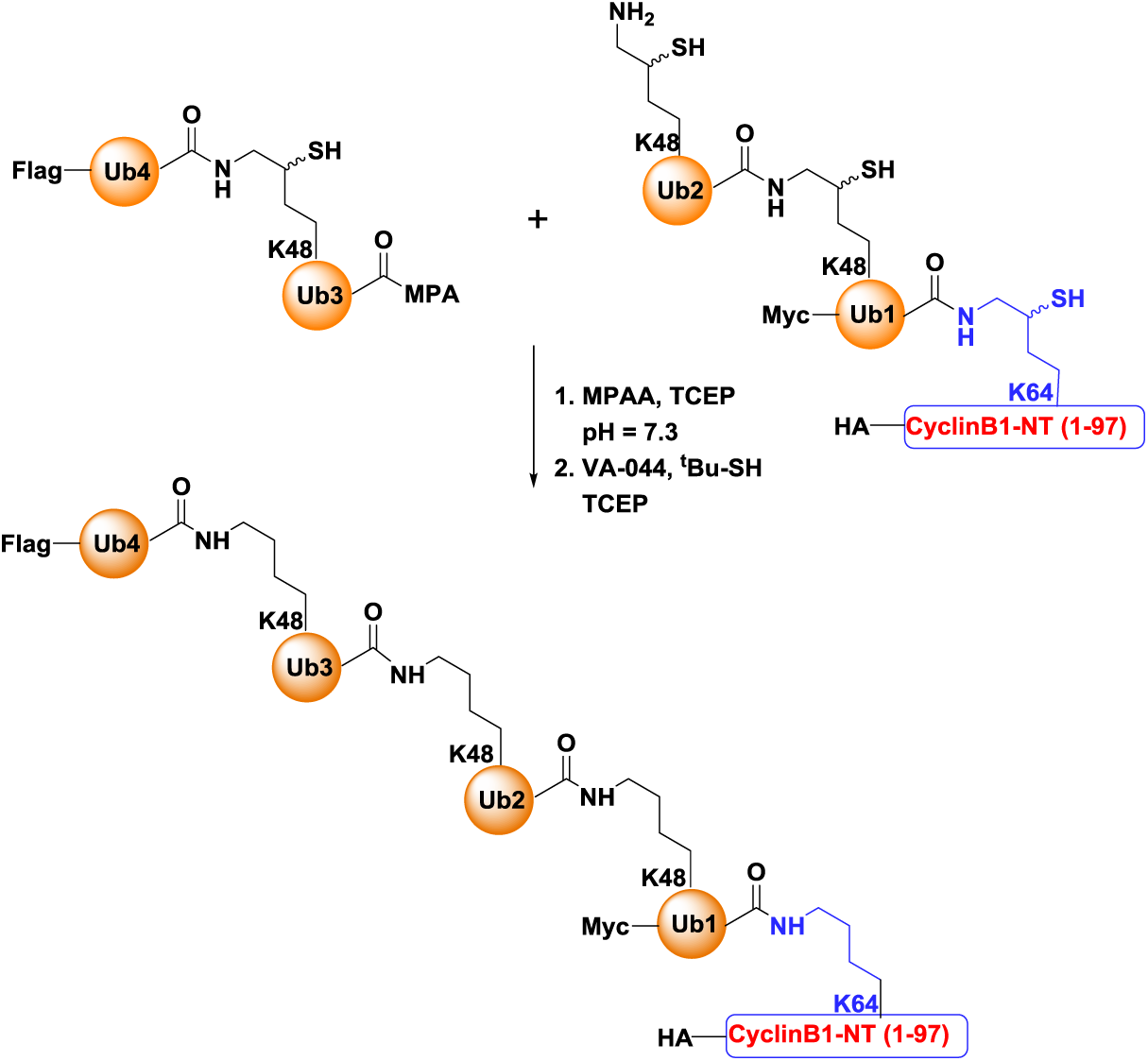
Synthesis of tetraUb-HA-CyclinB1-(NT)

**Figure M13.**
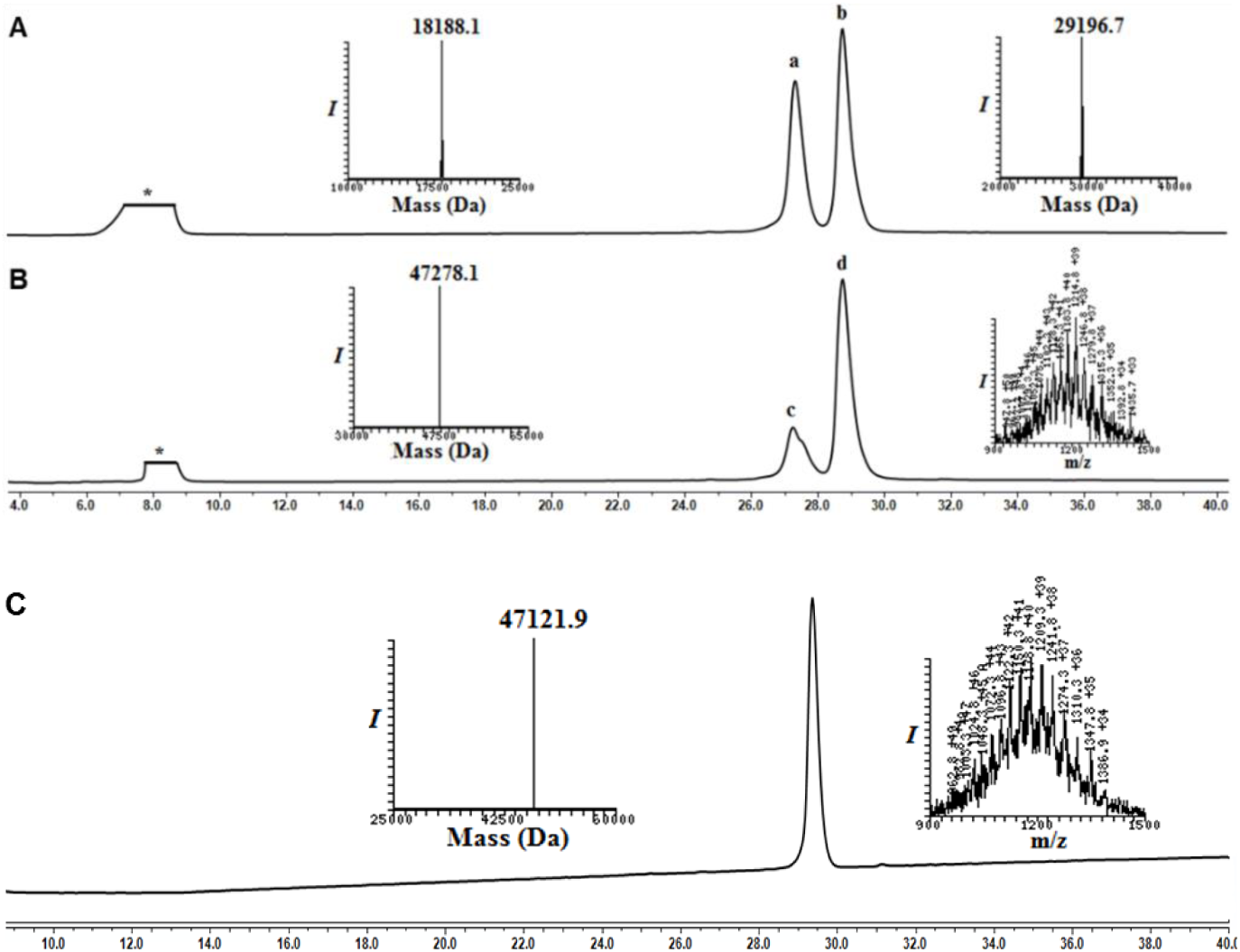
Synthesis of TetraUb^K48^-HA-CyclinB1-NT: (**A**) Analytical HPLC and mass traces of the ligation at 0 h, peak **a** corresponds to Flag-DiUb^K48^-MPA, peak **b** corresponds to Myc-DiUb^K48^-HA-CyclinB1-NT. (**B**) Ligation after 6hrs; peak **c** corresponds to the hydrolyzed Flag-DiUb^K48^-MPA thioester, peak **d** corresponds to ligation product TetraUb^K48^-HA-CyclinB1-NT with the observed mass of 47278.1 Da (calculated 47279.9 Da). **(C)** Analytical HPLC and mass traces for the desulfurized and purified TetraUb^K48^-HA-CyclinB1-NT with the observed mass 47121.9 Da (calculated 47124.9 Da).

### Purification of human Proteasomes

To obtain human proteasomes (Ding et al., 2019), human erythrocytes (RBCs purchased from The Israel blood bank) were washed with chilled 1X PBS and lysed in hypotonic buffer (25 mM Tris pH 7.4). The cell debris were separated by centrifugation and the clear red cell lysate were loaded onto 50 mL DEAE Affigel blue column (*Biorad*) and washed with Buffer A (25 mM Tris pH 7.4, 10% glycerol, 10 mM MgCl_2_, 1 mM ATP, 1 mM DTT). Proteasome was eluted with a gradient elution from 0%– 50% of Buffer B (25 mM Tris pH 7.4, 10% glycerol, 10 mM MgCl_2_, 1 mM ATP, 1 mM DTT with 1 M NaCl). The active fractions were then checked by 25 mM Suc-LLVY-AMC activity assay (Glickman and Coux, 2001; Leggett et al., 2005), pooled and loaded onto 8 mL Resource-Q column with Buffer A. Then proteasomes were eluted with a gradient elution from 0%–50% of Buffer B and active fractions were then checked by activity assays conducted either in a 96-well plate or native PAGE. To obtain the highest purity, gel filtration was performed for each fraction using a S300 (120 mL) column (GE Life Sciences) with Buffer A. The purity and integrity of proteasomes were checked in native-PAGE and SDS-PAGE, then concentrated, aliquoted, flash frozen in liquid nitrogen and stored at −80°C.

### Enzymatic ubiquitination and separation of K48-Ub chains

Recombinant Monomeric Ub, E2 conjugating enzymes, and human E1 were purified (Nakasone et al., 2013). Enzymatically synthesized Lysine48-Ub chains were obtained from a reaction containing 1 mM of Ub or Ub-His6, 80 nM E1 (UBA1), 40 mM E2-25K, 4 mM TCEP, and 15 mM ATP in a volume of 1 mL with a 50 mM Tris pH 8.0 buffer incubated at 37°C for 20 h (Mansour et al., 2015). Then, the reaction mixture was resolved by SEC-70 10/300 (*Biorad*) in Tris pH 7.4 buffer with 150 mM NaCl and 1 mM DTT. The ubiquitin chain fractions were confirmed by SDS-PAGE and fractions of shorter chains (≤4 Ub) were pooled and acidified with a final 50 mM NH4Acetate pH 4.0. Then the mixture loaded into Resource-S (6mL) column with cationic exchange buffer A (50 mM NH4 Acetate 4.5 pH), eluted with a gradient step from 10%–30% buffer B (50 mM NH4Acetate 4.5 pH with 1 M NaCl) for 20 column volumes (cv) and the purity of the chains were confirmed by SDS-PAGE. To obtain the highest purity, gel filtration was performed for each fraction using a SEC 70 10/300 (*Biorad*) in Tris pH 7.4 buffer with 150 mM NaCl and 1 mM DTT. Purity was confirmed by SDS-PAGE and proteins were concentrated, aliquoted, and stored at −20°C.

### Lysis of mammalian cell culture and Immunoblotting

Cell culture plates were washed twice with warmed 1X PBS and cell lysates were prepared by adding chilled NP-40 Lysis buffer (50 mM Tris 7.4 pH, 150 mM NaCl, 1% NP-40, 1X Protease inhibitor cocktail, 5 mM iodoacetamide) to the culture plates. The lysate content were collected in small centrifuge tube and incubated on ice for 30 min followed by centrifugation at 4°C/13000 rpm. The supernatant lysate was collected and stored at −20°C until further use. Denaturing immunoblotting was carried out using various antibodies.

### Immunoprecipitation for Puromycin-conjugates and Ubiquitin conjugates

The cell lysates were prepared as above with a minor change in NP-40 concentration (0.4%). Then pre-cleared cell lysates were prepared by incubating with Protein A/G conjugated agarose beads for 30 min at 4°C with slow rotation. 1 mg Pre-cleared cell lysates were then added to 20 µL of fresh Protein A/G conjugated agarose beads along with 4 µg of antibodies (anti-Puromycin/anti-Ubiquitin). The mixture was incubated for 2 h at 4°C with slow rotation followed by five 1 mL washes with chilled wash buffer (50 mM Tris 7.4 pH, 300 mM NaCl, 1X Protease inhibitor cocktail, 5 mM iodoacetamide). The target conjugates were then eluted from beads by mixing protein loading dye (without reducing agents BME/DTT) and heating for 2 min at 95°C followed by denaturing Western blot.

### Native cell lysis for proteasome activity

The cultured cells were collected from plate by trypsinization and lysed using ATP-buffer (25 mM Tris 7.4 pH, 10% glycerol, 10 mM MgCl_2_, 1 mM ATP, 1 mM DTT) with glass beads and vortexing at 4°C. Then after centrifugation the clear supernatant (Native lysate) were collected, aliquoted, flash frozen in liquid nitrogen and stored at – 80°C.

### Native gel, In-gel activity assay and Native Immunoblotting

Native lysates were run in 4% native PAGE in Native running buffer (100 mM Tris Base, 100 mM Boric acid, 1 mM EDTA, 2.5 mM MgCl_2_, 0.5 mM ATP, 0.5 mM DTT) for 2 h at 130 V. For In-gel activity assay the gel was incubated in ATP-Buffer with 25 µM Suc-LLVY-AMC at 37°C for 10 min and image under UV light. For Native western the gel was soaked in SDS-Running buffer for 20 min and then transferred onto PVDF membrane.

### *In vitro* substrate degradation assay

20 nM of human 26S or 20S proteasome and 2 μM of Naked CyclinB1-NT and ubiquitinated CyclinB1-NT conjugates were used for the degradation assay in a molar ratio of 1:100. 26S proteasome assay buffer contain 25 mM TRIS (pH 7.4), 10 mM MgCl_2_, 10% Glycerol, 1 mM ATP and 1 mM DTT. For 20S proteasome, assay buffer contains 25 mM TRIS (pH 7.4), 150 mM NaCl, 10% Glycerol. All degradation reaction were carried out at 37°C and terminated by adding SDS-loading dye.

### Peptide isolation from *in vitro* reactions

For proteasome processed peptide isolation the above proteasome-substrate reactions are terminated by heating mixtures at 95°C for 5 min. Then the mixture was passed through 30 KDa cut-off Amicon® Ultra - 0.5 mL Centrifugal Filter unit (*Merck-Millipore*). The flow-through was collected and desalted by C18 StageTip column.

### Intracellular peptide isolation

The cultured cells were collected from plate by trypsinization, then lysed by adding 80°C hot water and incubating the mixture at 80°C for 20 min (Dasgupta et al., 2014). Then the lysate was centrifuged at high speed (16,000g) for 30min and the supernatant was kept at –80°C over night. The lysate was then acidified with HCl to a final concentration of 0.01N HCl and was incubated on ice for 30min. Following 20 min centrifugation at 16,000g the pellet was discarded and the supernatant is passed through 30KDa cut-off Amicon® Ultra - 0.5mL Centrifugal Filter unit (*Merck-Millipore*). The flow-through was collected and desalted by C18 StageTip column.

### Human Heart Tissue Procurement

Ventricular myocardial tissue from patients with advanced heart failure (Failing heart) at the University of Michigan was collected at the time of cardiac transplantation as described in (Day et al., 2013). Non-failing ventricular myocardial tissue was collected from unmatched donors from the University of Michigan and other regional hospitals through a protocol with the Gift of Life Michigan Organ and Tissue Donation Program. This project was approved by the University of Michigan Institutional Review Board (IRB) and subjects gave informed consent. Each heart was perfused with ice-cold cardioplegia, samples were snap frozen in liquid N_2_ and stored at –80°C. Patient demographics were recorded at the time of tissue collection.

### Heart-muscle tissue lysate preparation for proteasomes and for intracellular peptides

The frozen heart muscle tissue (∼0.5 g) was resuspended with 300 µL of chilled PBS and quick homogenized into suspension. The tissue suspension was divided into two parts; (a) for proteasome preparation and (b) for intracellular peptide isolation. For proteasome lysate preparation 1 mL of chilled ATP buffer was added and lysed with glass beads and vortexed at 4°C. Then after high speed centrifugation (16,000g) for 30 min the clear supernatant were collected, flash frozen in liquid nitrogen and stored at – 80°C. For intracellular peptide isolation 1 mL of 80°C hot water was added to the tissue suspension and processed as described above.

### Proteasome enrichment and Proteasome Trapped Peptides (PTPs) isolation

The native proteasome lysates were subjected to ultra-centrifugation at 100,000 g for 45 min at 4°C. The first ribosome pellet was discarded and the supernatant was further centrifuged at 150,000g for 4hr at 4°C. Then the pellet was collected, washed and resuspended with ATP buffer. For PTPs isolation 80°C hot water was added to the enriched proteasome suspension, incubated at 80°C for 20 min and processed as described in “Intracellular peptide isolation” section.

### Cell proliferation and survival assay

A total of 5000 HeLa cells were seeded in each well of a 96-well plate and grown either under hypoxia (1% O_2_) and Normoxia. After 24 h of seeding the cells were treated with puromycin at different concentration (from 250 ng/mL to 1000 ng/mL) for different time periods (from 0 h to 60 h). At each time point 100 µg of MTT was added per well and 6 h later 10% SDS (in 0.01N HCl) solution was added to dissolve the crystals. 16 h post-addition of SDS the colorimetric reading was taken at 570 nm & 690 nm. Similarly, Doxycycline treated and untreated PSMD2-KD inducible HEK293 cells were seeded at concentration (5000 cells/well) assay was performed as HeLa cells.

### LC-MS/MS and Mass-spectrometry analysis

Desalted peptides were analyzed by liquid chromatography coupled mass spectrometry. Samples of *in vitro* cleavage reactions were analyzed using Orbitrap Fusion Tribrid (*Thermo Scientific*) coupled to Ultimate 3000 Nano Systems (*Thermo Scientific*) following loading via an Acclaim PepMap 100 trap column (100 μm × 2 cm, nanoViper, C18, 5 μm, 100 å; *Thermo Scientific, Waltham, MA*) onto an Acclaim PepMap RSLC analytical column (75 μm × 50 cm, nanoViper, C18, 2 μm, 100 å; *Thermo Scientific*). The peptides were eluted with a linear 30 min gradient of 6–30%, followed by 3 min gradient of 30–34%, 5 min gradient of 34–76% and 10 min wash at 76% acetonitrile with 0.1% formic acid in water (at flow rates of 0.2 μl/min). These MS analyses done In positive mode using HCD when each cycle was set to a fixed cycle time of 2s and consisted of an Orbitrap full ms1 scan (range: 375-1575 m/z) at 60.000 resolution with AGC target: 1e6; maximum IT: 118 ms, followed 18 Orbitrap ms2 scans (range 140-2000) at 60,000 resolution: with AGC target: 4e5; maximum IT: 118 ms; isolation window: 1.4 m/z; HCD Collision Energy: 32%. Dynamic exclusion was set to 10 s and the “exclude isotopes” option was activated.

The rest of the peptidomics samples were analyzed using Q-Exacitive-Plus or Q-Exactive HF mass spectrometer (*Thermo Fisher*) coupled to nano HPLC. The peptides were resolved by reverse-phase chromatography on 0.075 × 180 mm fused silica capillaries (J&W) packed with Reprosil reversed-phase material (*Dr. Maisch; GmbH, Germany*). The peptides were eluted with a linear 60 min gradient of 5–28%, followed by a 15 min gradient of 28–95%, and a10 min wash at 95% acetonitrile with 0.1% formic acid in water (at flow rates of 0.15 μl/min). Mass spectrometry analysis by Q Exactive HF-X mass spectrometer (*Thermo Fisher Scientific*) was in positive mode using a range of m/z 300–1800, resolution 60,000 for MS1 and 15,000 for MS2, using repetitively full MS scan followed by high energy collisional dissociation (HCD) of the 18 most dominant ions selected from the first MS scan.

### Mass-spectrometry data analysis

MS data analysis was done with either MaxQuant version 1.6.7.0 (Cox and Mann, 2008) or the Trans Proteomic Pipeline (TPP) v5.2.0 Flammagenitus (Deutsch et al., 2015). MaxQuant searches were done using unspecific Digestion mode with minimal peptide length of 7 amino acids. Search criteria include oxidation of methionine and protein N-terminal acetylation as variable modifications. All other parameters were set as the default. The raw files were searched against the Homo sapiens UniProt fasta database (November 2017; 20,239 sequences) supplemented with the sequences of the fusion protein of CyclinB1-NT (1-88 aa residue) with NS3-protease. Candidates were filtered to obtain FDR of 1% at the peptide and the protein levels. No filter was applied to the number of peptides per protein. For quantification, the match between runs modules of MaxQuant were used, and the LFQ (label free quantification) normalization method was enabled.

TPP searches were done following RAW files conversion to mzML using MSConvert (ver 3.01157) using centroid and compressing peak list option. Searches were performed using Comet (2017.01 rev. 1) and high-resolution settings. They include ‘no enzyme’ cleavage specificity and oxidation of methionine and protein N-terminal acetylation as variable modifications. All searches were conducted against protein sequences downloaded from UniProt (https://doi.org/10.1093/nar/gky1049). The searches of *in vitro* cleavage reactions of cyclin B1 were done with a database composed of all Uniport human proteome (November 2017; 6721 sequences), all of Human proteasome-related proteins (based on “Proteasome” Keyword in Uniprot, November 2017; 169 sequence), the sequences of modified cyclin B1, FLAG-tagged ubiquitin and Myc-tagged ubiquitin, and the sequence of the cRAP contaminant database (cRAP database released Feb 2012; 115 entries). All these sequences were supplemented with decoy sequences. Searches of other experiments were done against the Homo sapiens UniProt fasta database (November 2017; 20,239 sequences) supplemented with the sequences of the fusion protein of CyclinB1-NT (1-88 aa residue) with NS3-protease and the sequence of the cRAP contaminant database (cRAP database released Feb 2012; 115 entries). Peptide Prophet was used to curate peptide-spectrum matches of FDR ≤1% and assign representative proteins/protein isoforms.

Label-free quantification of ubiquitin peptides was performed using Skyline version 4.2 (Pino et al., 2017). Peptide library was constructed based on database search results obtained for heart failure samples searched by MaxQuant as described above. The RAW files of these samples were used to calculate the sum of peak area for all peptides originated from ubiquitin.

### Bioinformatics data analysis and Statistics

The sample clustering and heatmap analysis of naked- and ubiquitinated-CyclinB1-NT *in vitro* cleavage experiment was performed as follows. The PSM counts per peptide were converted into a sample-by-peptide matrix and samples with less than 50 PSMs were discarded, along with the peptides observed only in single samples. The distances between the samples were obtained by performing MDS analysis as implemented in the plotMDS function from edgeR v. 3.26. (Robinson et al., 2010). The distances from resulting ordinations were utilized to cluster the samples using the Ward’s method. The heatmap was obtained by normalizing the PSM counts as counts-per-million and rescaling them to the maximum PSM counts per peptide. The maxima were kept track of and plotted alongside the heatmap. Analogous analyses were performed for P1 positions instead of peptides.

Cleavage site amino acid composition preferences were analyzed by building information-content sequence logos and differential sequence logos using dagLogo v. 1.22.2 and DiffLogo v. 2.8.0, respectively (Nettling et al., 2015).

For the heart failure peptidomics experiment the PSMs were obtained as described above. Ordination of samples in MDS axes was performed with edgeR and was based on protein-level PSM counts.

Differences between size distributions of cleavage peptides in multifactorial experimental designs were analyzed by fitting Poisson-family GLMs with lme4 v. 1.1- (Bates et al., 2015), inspecting significant model terms with ANOVA and running *post hoc* tests with emmeans v.1.3.5. Otherwise, pairwise differences were tested with the Mann-Whitney U test.

Most of the data manipulation steps, diagrams and statistical tests were performed in R v.3.6.1.

The prediction of protein disorder was done using IUPRED2A (Meszaros et al., 2018) (https://iupred2a.elte.hu/). The sequences of relevant proteins were downloaded from Uniport and the resulting FATSA file was submitted prediction with IUPRED2 using the default settings. The Disorder score for each protein was set as the average value of IUPRED score of its amino acids (calculated by in-house script). Boxplot graphs were generated using BoxPlotR (Spitzer et al., 2014) (http://shiny.chemgrid.org/boxplotr/) or Graphpad Prism V5.

### CyclinB1-NT cleavage site (P1) calculation

Peptides generated from both Naked-CyclinB1-NT and TetraUb-CyclinB1-NT by 20S or 26S *in vitro* were searched by TPP- and assigned PSMs (MS/MS-count) to each peptide. Only those peptides that showed up in two out of three replicates were consider for further analysis. P1 positions at cleavage sites were assigned with a value calculated from integrated MS/MS count of all the corresponding peptides. Next, the relative cleavage preference for 20S over 26S at each P1 position was calculated and represented as Log_2_ of the corresponding ratios. For P1-position with a zero value was consider as 0.1 for the relative cleavage preference calculation. In the case of *in vivo* P1 position relative cleavage preference calculation, a similar method was followed with a modification that considered peptides appeared even in one replicate from independnet intracellular peptidomic analysis repeats.

### Cryo-EM sample preparation and data collection

Human 20S proteasome was mixed with monoUb-CyclinB1-NT at a molar ratio of 1:5, incubated on ice and frozen within 5 minutes. A volume of 2 µL sample was placed onto a glow-discharged Quantifoil holey carbon grid (R2/1, 200 mesh, Quantifoil Micro Tools), which was freshly coated with graphene oxide (*Sigma Aldrich*). The grid was blotted utilizing Vitrobot Mark IV (*FEI company*), and flash frozen in liquid ethane. 20S alone sample was handled in the same way without added substrate and incubation.

Images were taken by using a Titan Krios transmission electron microscope (*Thermo Fisher*) operated at 300 kV and equipped with a Cs corrector (**Figure M14A and M15A**). Images were collected by using a K2 Summit direct electron detector (Gatan) in super-resolution mode at a nominal magnification of 18,000x, yielding a pixel size of 1.318 Å after 2 times binning (**Figure M14 and Table S1**). Each movie was dose-fractioned into 38 frames with a dose rate of 8 e^−^/pixel/s on the detector. The exposure time was 7.6 s with 0.2 s for each frame, generating a total dose of 38 e^−^/Å^2^. Defocus value varies from −0.9 to −1.8 µm. All of the images were collected by utilizing the SerialEM automated data collection software package (Mastronarde, 2005).

### Single particle cryo-EM data processing

3,125 movies for the 20S+monoUb-CyclinB1-NT sample and 748 movies for the 20S alone sample were collected for structural determination. Single particle analysis was executed in RELION 3.0. All images were aligned and summed using MotionCor2 software (Zheng et al., 2017). After CTF parameter determination using CTFFIND4 (Rohou and Grigorieff, 2015) and Gctf (Zhang, 2016), particle auto-picking, manual particle checking, and several rounds of reference-free 2D classification and 3D classification, 282,834 particles for 20S+monoUb-CyclinB1-NT and 154,436 particles for 20S alone remained for further processing. After applying CTF refinement and Bayesian polishing to the remaining particles, they were auto-refined without imposing any symmetry and reached 3.38 Å resolution for 20S+monoUb-CyclinB1-NT and 3.22 Å resolution for 20S alone after post-processing.

A further 3D classification into 5 classes was performed for each of the two datasets. In the 20S+monoUb-CyclinB1-NT dataset, class 1 with an obviously asymmetric gate configuration (in magenta, 29.1% of the population) is distinct from the other classes (**Figure M15B**). We then performed two more rounds of 2D classification on class 1 data to discard misaligned or broken particles. Further refinement on these cleaned-up particles led to a map at 4.47 Å resolution (denoted as S1; **Figure M15C and M15D**). For the other three classes with reasonably good structural features (2, 4, and 5), since they all appear symmetrically closed in the two rings and show highly similar features, their particles were combined and refined into a map at 3.88 Å resolution (denoted as S2; **Figure M15D**) with normal 20S configuration. For the 20S alone dataset, further classification did not yield significant differences between classes for the four classes showing reasonably good structural features (1 to 4; **Figure M14**), and their two gates all appear symmetrically closed. For further structural analysis, we used the reconstructions before postprocessing, since these maps present better structural feature connectivity.

**Figure M14.**
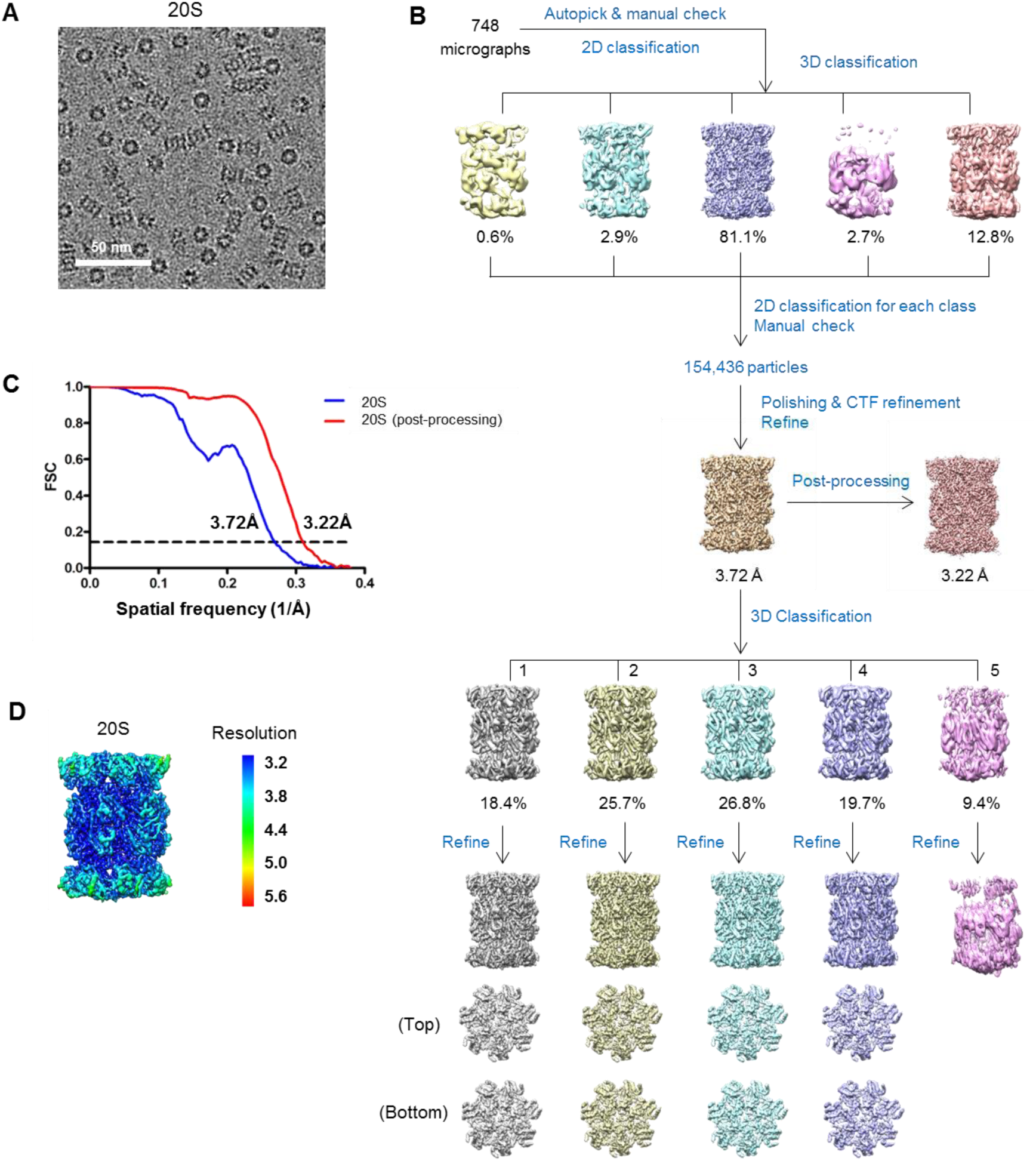
Workflow and validation for the cryo-EM data processing of 20S alone (**A**) Representative cryo-EM micrograph of 20S proteasome. (**B**) Micro 3D classification and refinement procedures. After 2D and 3D classifications to eliminate bad particles, there are 154,436 remaining cleaned-up particles, based on which we obtained an overall map for 20S alone sample. Further 3D classification did not yield significant differences between classes for the four classes with reasonably good structural features (1 to 4), and their two gates all appear symmetrically closed. (**C**) Resolution estimation of these obtained maps according to the gold standard FSC criterion of 0.143. (**D**) Local resolution estimation for the 20S alone map. The resolution color bar (in Å) is also shown.

**Figure M15.**
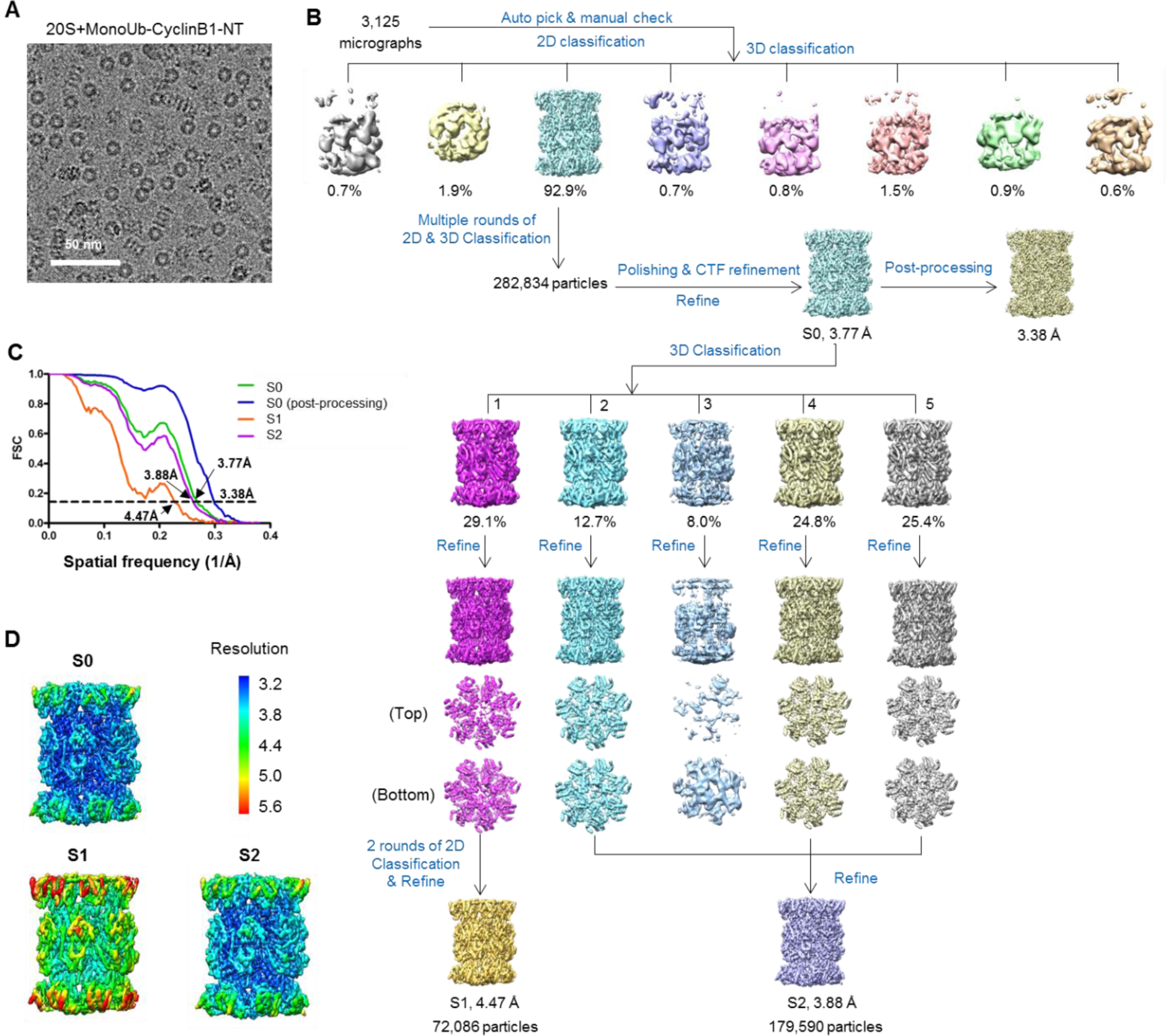
Workflow and validation for the cryo-EM data processing of 20S+monoUb-CyclinB1-NT. (**A**) Representative cryo-EM micrograph of 20S proteasome incubated with monoUb-CyclinB1-NT. (**B**) 3D classification and refinement procedures. After 2D and 3D classifications to eliminate bad particles, there are 282,834 remaining cleaned-up particles, based on which we obtained an overall map for 20S+monoUb-CyclinB1-NT (named as S0). A further 3D classification into 5 classes generated a class (class 1) with an obviously asymmetric gate configuration (in magenta, 29.1% of the population), which is distinct from the other classes. This class was further processed to generate a map denoted as S1. For the other three classes with reasonably good structural features (2, 4, and 5), since they all appear symmetric between the two rings and show highly similar features, their particles were combined and refined into a map denoted as S2. (**C**) Resolution estimation of these obtained maps according to the gold standard FSC criterion of 0.143. (**D**) Local resolution estimations for the S0, S1, and S2 maps. Their resolution color bars (in Å) are also shown.

### Pseudo-atomic model building and structural analysis

The available atomic model 6rgq of human 20S proteasome (Toste Rego and da Fonseca, 2019) was used as initial model. We performed further flexible fitting of the model against the corresponding map by using the real-space refinement function in Rosetta (DiMaio et al., 2015). The generated model was further refined utilizing PHENIX (Adams et al., 2010), and manually modified using COOT (Emsley and Cowtan, 2004).

UCSF Chimera and ChimeraX were used for figure generation (Goddard et al., 2018; Ludtke et al., 1999; Pettersen et al., 2004). For 20S+monoUb CyclinB1-NT, the difference map was generated by using a 180° rotated (around the pseudo 2-fold axis) 20S+monoUb CyclinB1-NT map minus its original unrotated map. Program *proc3d* in EMAN1.9 was used for differential map generation (Ludtke et al., 1999). For the difference map of 20S alone, similar procedure was adopted.

## Supplemental Information

**Figure S1.**
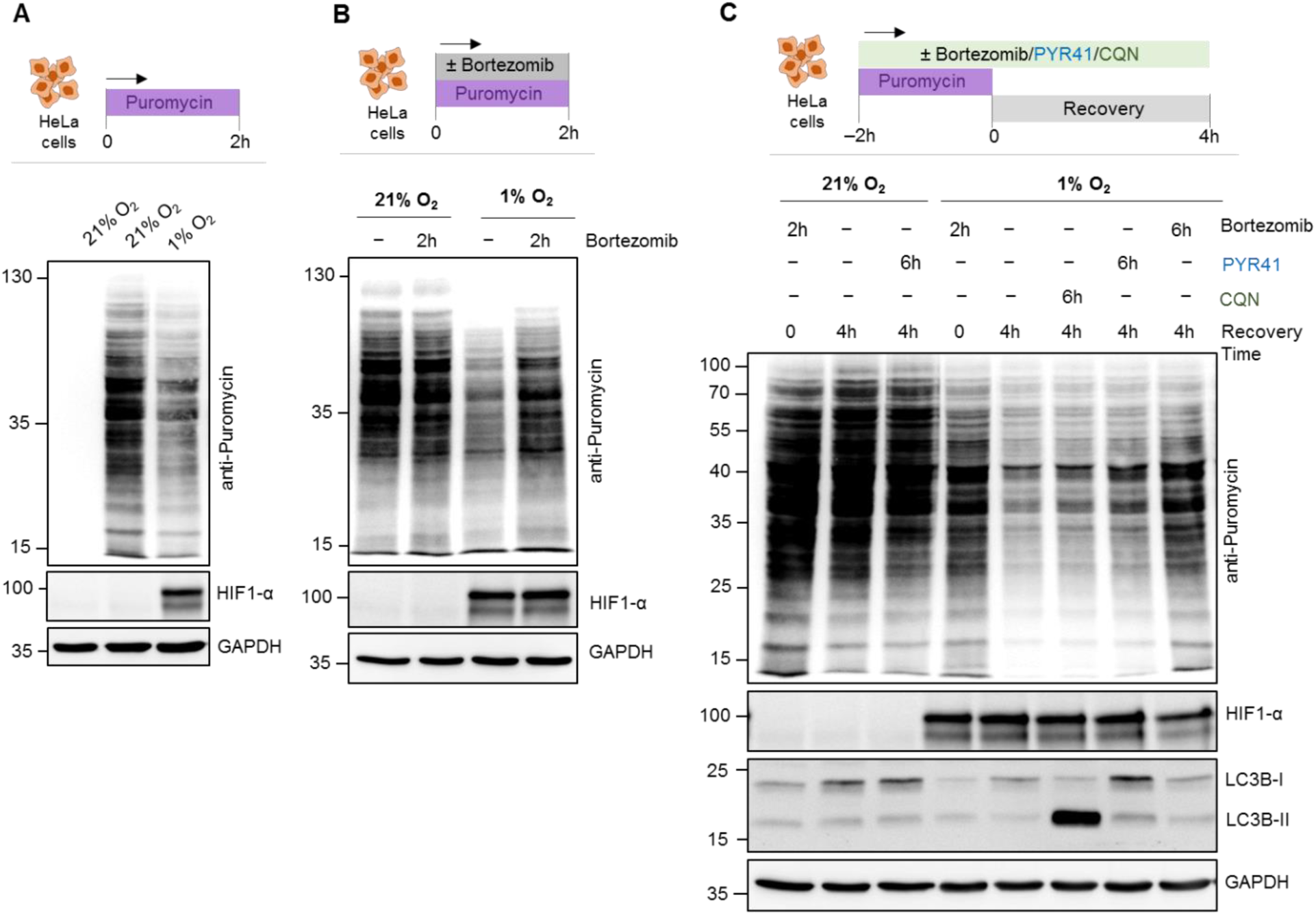
Puromycinated-nascent proteins were removed rapidly by ubiquitin-independent proteasome pathway under hypoxia, Related to Figure 1. (**A**) HeLa cells were grown under either normoxia or hypoxia (1% O_2_) for 24 h followed by 2 h puromycin (5 µg/mL) treatment or left untreated. Cell lysates were resolved by SDS-PAGE to detect puromycin-conjugates by IB. (**B**) HeLa cells grown under normoxia or hypoxia (1% O_2_) for 24 h followed were treated with puromycin (5 µg/mL) for 2 h followed by ±bortezomib (1 µM) then the cell lysates were resolved by SDS-PAGE to detect puromycin-conjugates by IB. (**C**) HeLa cells were grown under either normoxia or hypoxia (1% O_2_) for 24 h followed by 2 h puromycin (5 µg/mL), pulse with/without bortezomib (1 µM)/PYR (10 µM)/Chloroquine (50 µM) and then recovery for indicated time periods with/without the inhibitors as indicated. Cell lysates were resolved in SDS-PAGE to detect puromycin-conjugates by IB.

**Figure S2.**
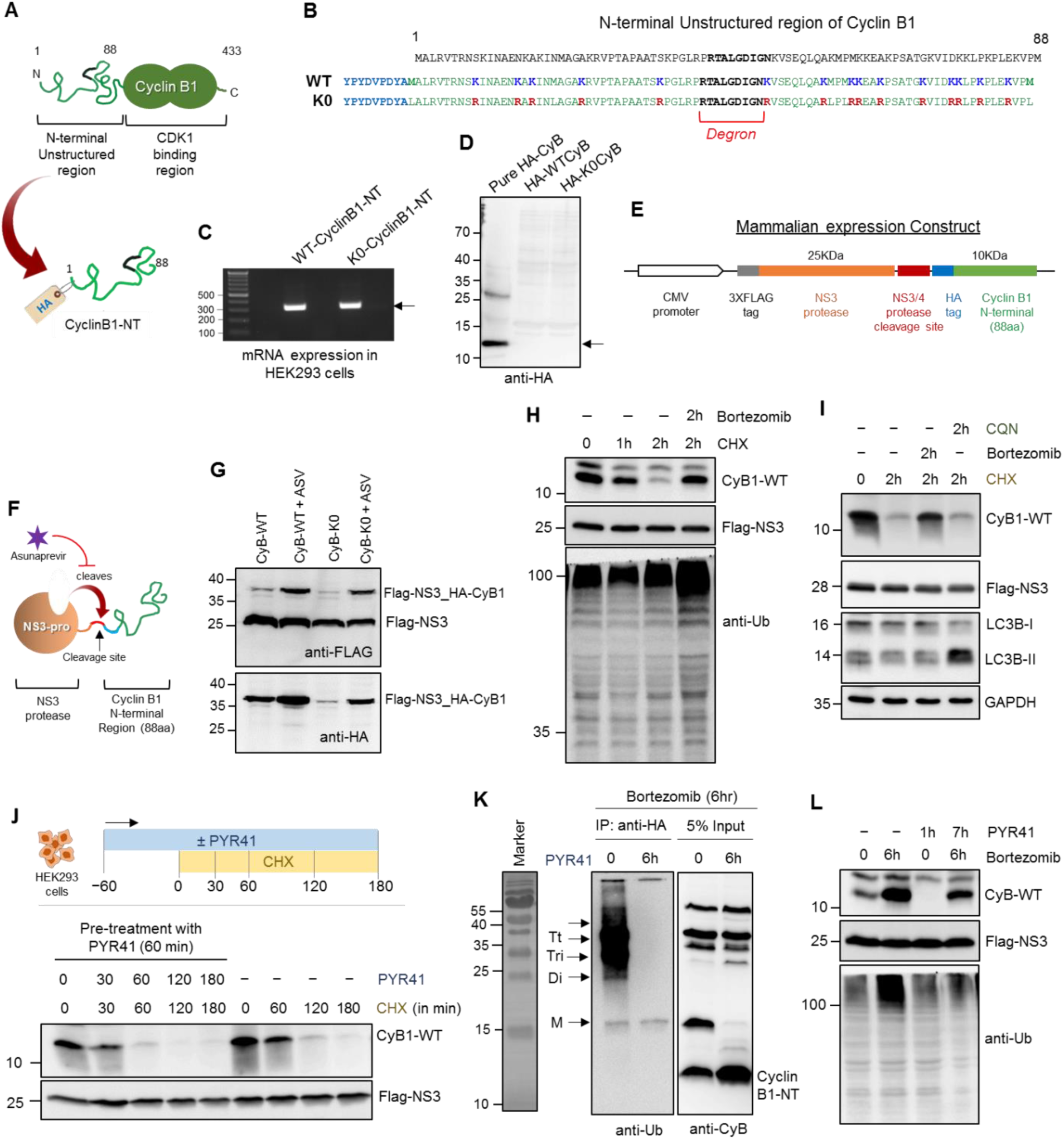
CyclinB1-NT is degraded by proteasome in a ubiquitin-independent manner, Related to Figure 2. (**A**) The cartoon figure represents the endogenous Cyclin B1 and the unstructured N-terminal region (C1-88 aa) taken for trans-expression with a HA-tag (HA-CyclinB1-NT). (**B**) The sequence of 88 aa residue long N-terminal region of Cyclin B1 consisting the Degron and 15 Lysine residues. Both wild-type (WT) and K0-mutant (all Lysine to Arginine) CyclinB1-NT was tagged with HA at N-terminus. (**C**) HEK293 cells were transfected with plasmid expressing either WT-or K0-HA-CyclinB1-NT. The RNA was isolated, and semiQ-PCR was performed to amplify the trans-expressed HA-CyclinB1-NT. The agarose gel picture represents the mRNA expression of both WT and K0-HA-CyclinB1-NT. (**D**) Cell lysate were prepared from the above transfected cells and resolved in SDS-PAGE along with a pure recombinant HA-CyclinB1-NT as positive control for IB. The IB image indicates no protein expression of the WT and K0 HA-CyclinB1-NT. (**E**) Cloning strategy for HA-CyclinB1-NT expression downstream to NS3-protease in mammalian expression vector. (**F**) The cartoon represents the mode of action of NS3-Pro which cleaves exactly before the HA tag of CyclinB1-NT and the activity of NS3-pro can be inhibited by an inhibitor – Asunaprevir (ASV) at 5 μM concentration. (**G**) The above construct (described in E) was expressed in HEK293T cells with or without 5 μM of Asunaprevir and 24 h post transfection cell lysates were subjected to IB using anti-HA and anti-Flag antibodies. (**H**) HA-CyclinB1-NT was trans-expressed in HEK293T cells with/without treatment of cycloheximide (50 μg/mL) and/or bortezomib (1 μM) for indicated time periods. IB was performed using indicated antibodies. (**I**) HA-CyclinB1-NT expressing HEK293T cells were treated with cycloheximide (50 μg/mL) and/or bortezomib (1 μM)/chloroquine (50 μM) for 2 h followed by IB. (**J**) HA-CyclinB1-NT expressing HEK293T cells were treated with cycloheximide (50 μg/mL) and/or PYR-41 (10 μM) for indicated time periods followed by IB. (**K**) HA-CyclinB1-NT expressing HEK293T cells were treated with bortezomib (1 μM) and/or PYR-41 (10 μM) for 6 h followed by immunoprecipitation of HA-CyclinB1-NT using anti-HA antibody. IB was performed using anti-ubiquitin antibody for immunoprecipitated fraction and anti-CyclinB1-NT for 5% Input. (**L**) HA-CyclinB1-NT expressing HEK293T cells were treated with/without bortezomib (1 μM) and/or PYR-41 (10 μM) for indicated time periods followed by IB.

**Figure S3.**
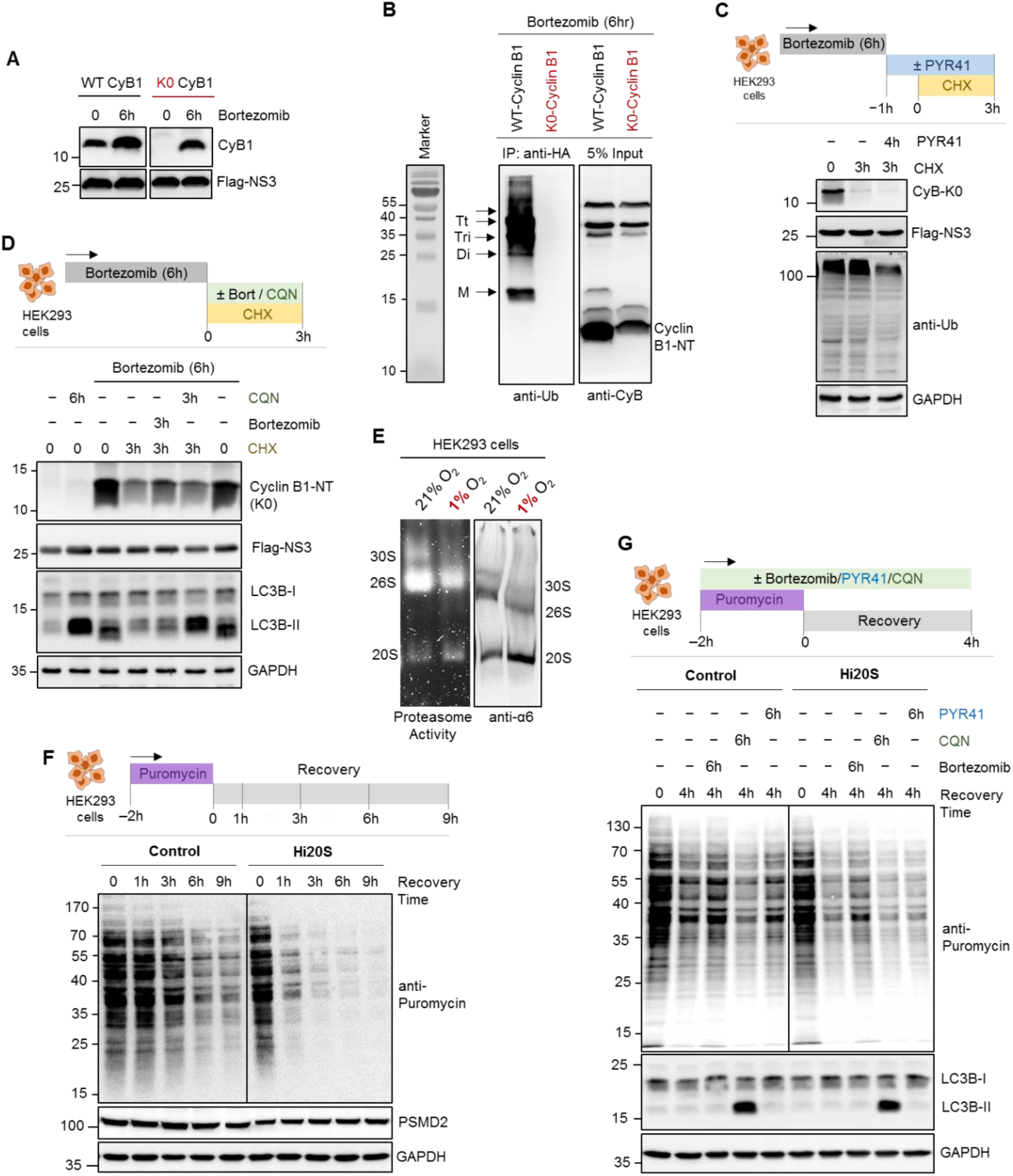
K0 CyclinB1-NT is degraded efficiently by proteasomes in a ubiquitin-independent manner similar as puromycin-conjugates, Related to Figure 2. (**A**) Wild type (WT) or K0-CyclinB1-NT were trans-expressed in HEK293T cells with/without treatment of bortezomib (1 μM) for 6 h. IB was performed using anti-CyclinB1-NT and anti-Flag antibodies. (**B**) K0-CyclinB1-NT expressing HEK293T cells were pulse treated with bortezomib (1 μM) for 6 h. Then bortezomib was removed and cells were grown with/without treatment of cycloheximide (50 μg/mL) and/or PYR-41 (10 μM) for indicated time periods followed by IB. (**C**) Wild type (WT) or K0-Cyclin B1-NT were trans-expressed in HEK293T cells followed by 6h pulse treatment of bortezomib (1 μM). HA-CyclinB1-NT was immunoprecipitated using anti-HA antibody. IB was performed using anti-ubiquitin antibody for immunoprecipitated fraction and anti-CyclinB1-NT for 5% Input. (**D**) K0-Cyclin B1-NT expressing HEK293T cells were pulse treated with bortezomib (1 μM). Then bortezomib was removed and cells were grown with/without treatment of cycloheximide (50 μg/mL) and/or bortezomib (1 μM)/Chloroquine (50 μM) for 2 h followed by IB. (**E**) HEK293T cells were grown either under normoxia or hypoxia (1% O_2_) for 24 h. Cells lysates were resolved in 4% native gel for proteasome in-gel activity assay or for native immunoblot (IB) using anti-α6 antibody. (**F**) PSMD2-KD inducible stable HEK293 cells were grown with/without doxycycline (1 μg/mL) and then given a pulse of puromycin (5 μg/mL) for 2 h. Followed by a recovery for different time periods as indicated and IB to detect puromycin-conjugates. (**G**) PSMD2-KD inducible stable HEK293 cells were grown with/without doxycycline (1 μg/mL) and then given a pulse of puromycin (5 μg/mL) for 2 h. Followed by a recovery for different time periods with/without bortezomib (1 μM)/chloroquine (50 μM)/ PYR-41 (10 μM) as indicated and IB was performed.

**Figure S4.**
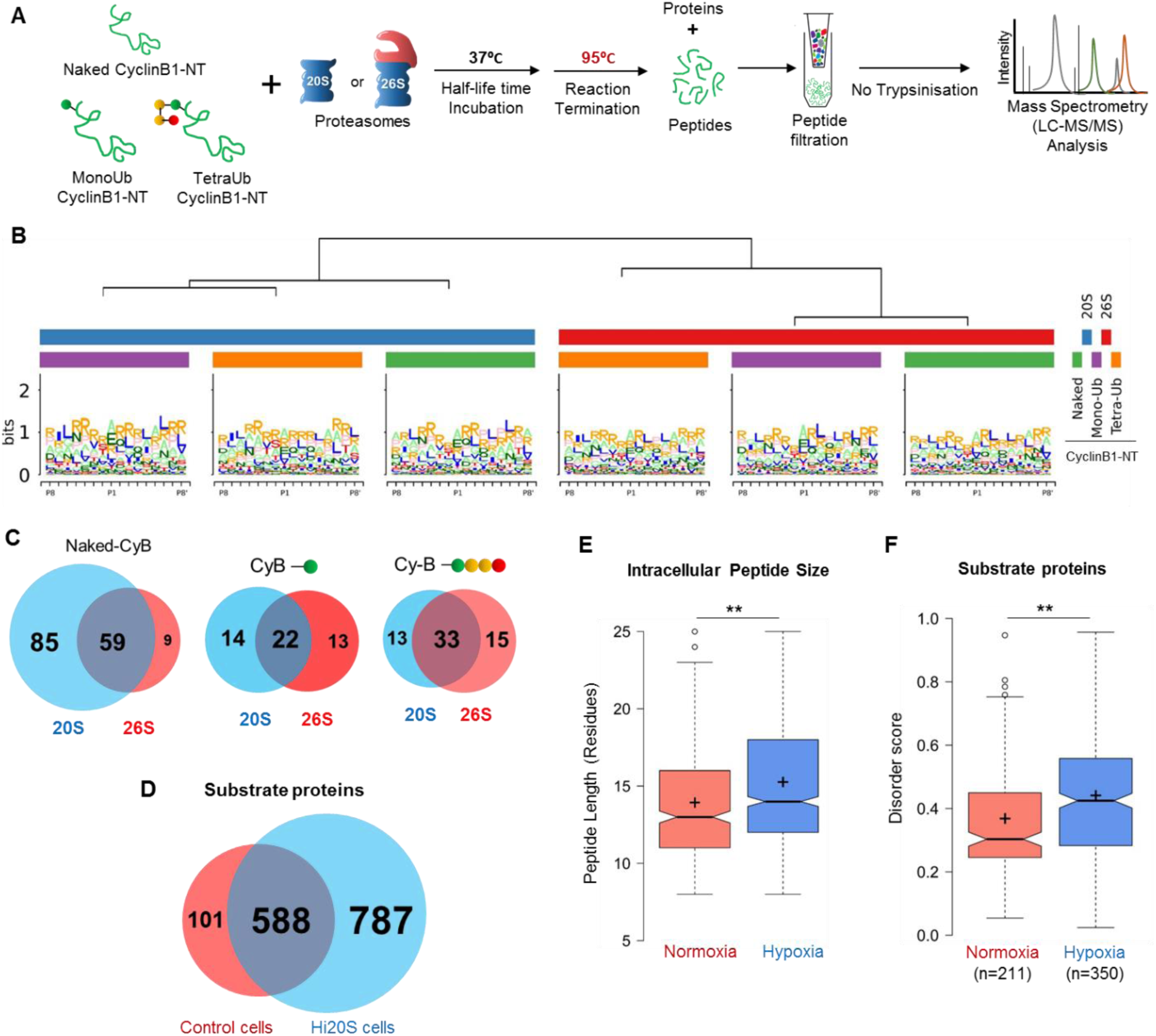
Intracellular peptide features represent signature of 20S proteasome in Hi20S and hypoxic cells, Related to Figure 4. (**A**) Naked-CyclinB1-NT, MonoUb-CyclinB1-NT or TetraUb-CyclinB1-NT were incubated separately at 37°C with either purified 20S or 26S proteasome at 1:50 (Substrate:Proteasome) molar ratio till the time points shorter than substrate half-life. Reactions were terminated by heating at 95°C and peptide products at each reaction were isolated and analyzed by LC-MS/MS as per Methods. (**B**) Sequence logo shows the amino acid preference for P1 cleavage sites on each substrate (above in A) for both proteasomes. (**C**) Venn diagrams represent the unique and common peptides generated by each proteasome from the given substrates. (**D**) Venn diagram represents potential proteasome substrates unique/common to each cellular condition (control, Hi20S). Proteins were assigned from intracellular peptide captured and scored based on intrinsic disordered elements. (**E**) Size distribution of intracellular peptides (Line, median; + mean size) obtained from HeLa cells grown under normoxia and hypoxia (from triplicate dataset). (**F**) Disorder score distribution of potential proteasome substrates unique to each cellular condition (normoxia and hypoxia). Proteins were assigned from intracellular peptide captured, and scored based on intrinsic disordered elements (notch, median; + mean).

**Figure S5.**
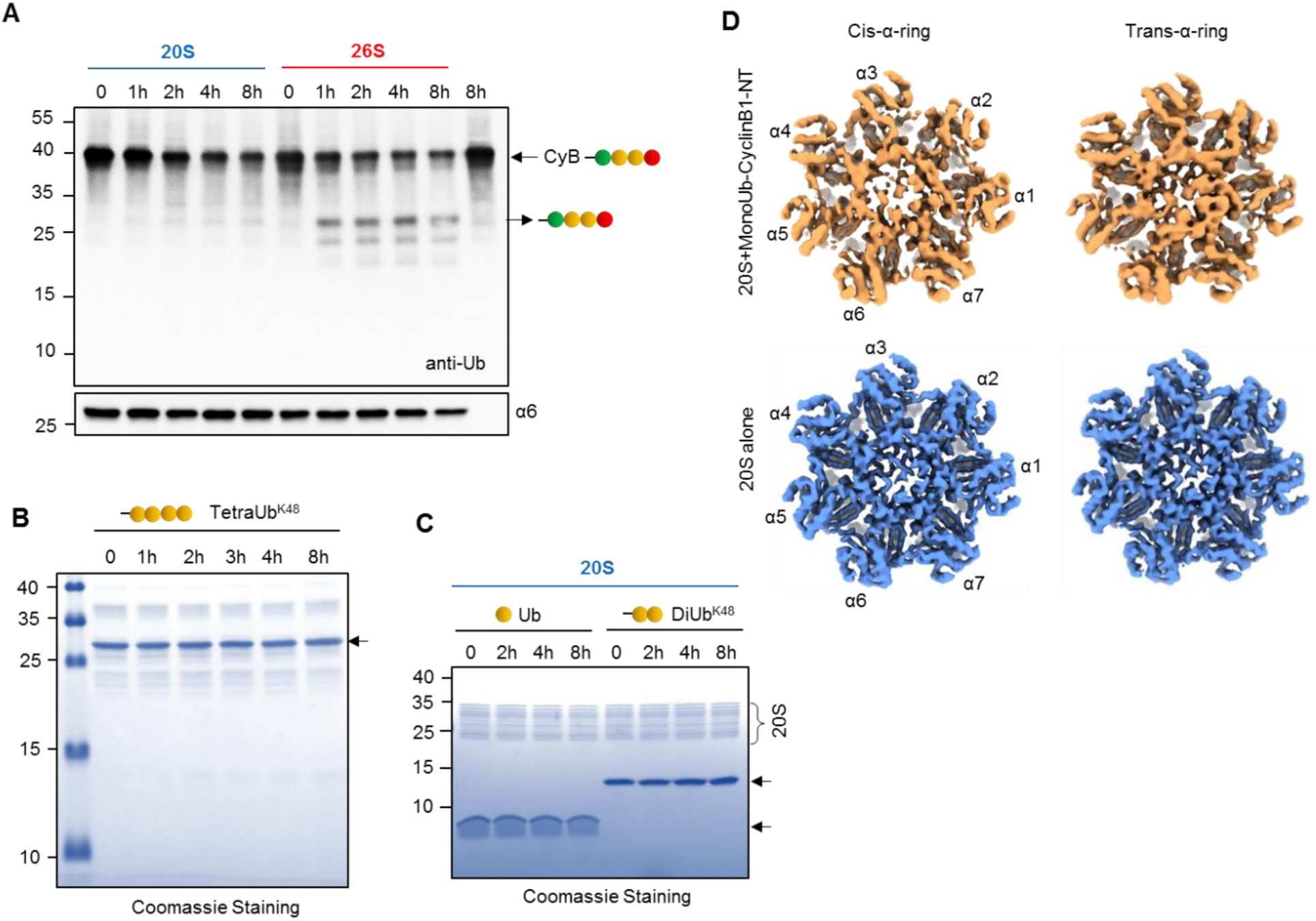
Evaluating the association of ubiquitin-conjugates with 20S proteasome, Related to Figure 5. (**A**) TetraUb-CyclinB1-NT was incubated at 37°C with purified 20S or 26S proteasome at 1:50 (Substrate:Proteasome) molar ratio for indicated time periods. The reaction mixture was resolved by Tris-Tricine PAGE and IB with anti-ubiquitin antibody to detect released ubiquitin units. (**B**) Free ubiquitin, DiUb^K48^ chain or (**C**) free TetraUb^K48^ chains were incubated at 37°C with purified 20S or 26S proteasome at 1:50 (Substrate:Proteasome) molar ratio for indicated time periods. The reaction mixture was resolved by Tris-Tricine PAGE and stained with Coomassie. (**D**) The views of the cis- and trans-α-rings of the cryo-EM map of S1 conformation (20S+monoUb CyclinB1-NT; in sandy brown; upper). The complex appears asymmetric between the cis α-ring and trans α-ring especially in the gate region, with notable loss of electron density mainly in the N-terminal tail regions of α2/α3/α4 subunits. The views of the cis- and trans-α-rings of the cryo-EM map of 20S alone (in blue; lower). The two rings appear in similar conformation with both gates in the closed configuration.

**Figure S6.**
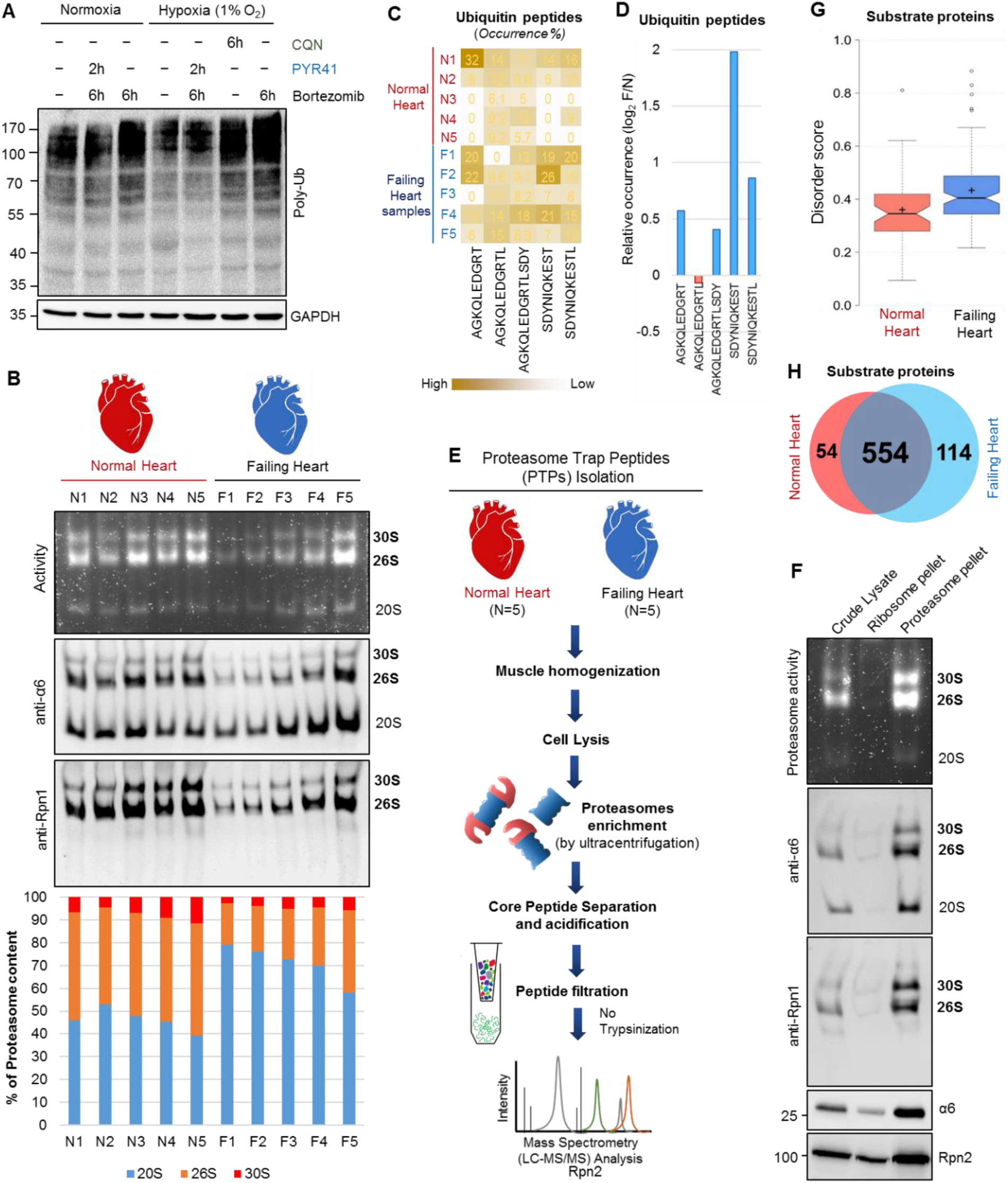
20S proteasome signature activity in human failing heart, Related to Figure 6. (**A**) Polyubiquitin content of HeLa cells grown under normoxia or under hypoxia either treated with chloroquine, PYR-41 or bortezomib for indicated time periods. IB using anti-ubiquitin antibody. (**B**) Proteasome activity (upper) and protein content (native IB: middle) of human heart muscle tissue under Failing (N = 5) or Normal (N = 5) conditions. Proteasome enrichment from frozen tissue followed the protocol in Methods and ratio of proteasome species for each sample was measured from anti-α6 probed native IB. The bar graph (lower) represents the percentage of proteasome content of Normal heart samples (N = 5) and Failing heart samples (N = 5). (**C**) Heat map represents the ubiquitin-derived peptides in heart muscle tissues. Human frozen heart muscle tissue under Failing (N = 5) and Normal (N = 5) conditions were subjected to intracellular peptide isolation following the protocol in Methods, quantitatively analyzed for ubiquitin peptides by LC-MS/MS. The values represent the relative occurrence of each ubiquitin peptide across 10 samples calculated from their LFQ-intensity. (**D**) The bar graph represents the relative occurrence of each ubiquitin peptides (calculated from integrated LFQ intensity) in Faling heart samples compared to normal heart samples. (**E**) Scheme describes approach for Proteasomal Trapped Peptides (PTPs) isolation and MS/MS analysis from human heart muscle tissues. (**F**) Qualitative analysis of proteasome enrichment method. Each fraction during the process (see Methods); e.g., crude lysates, ribosomal pellet and proteasome pellet were resolved in native gel to detect proteasome activity (upper) or proteasome protein content by native IB (middle). Lower denature PAGE IB shows two proteasome subunit levels.

**Figure S7.**
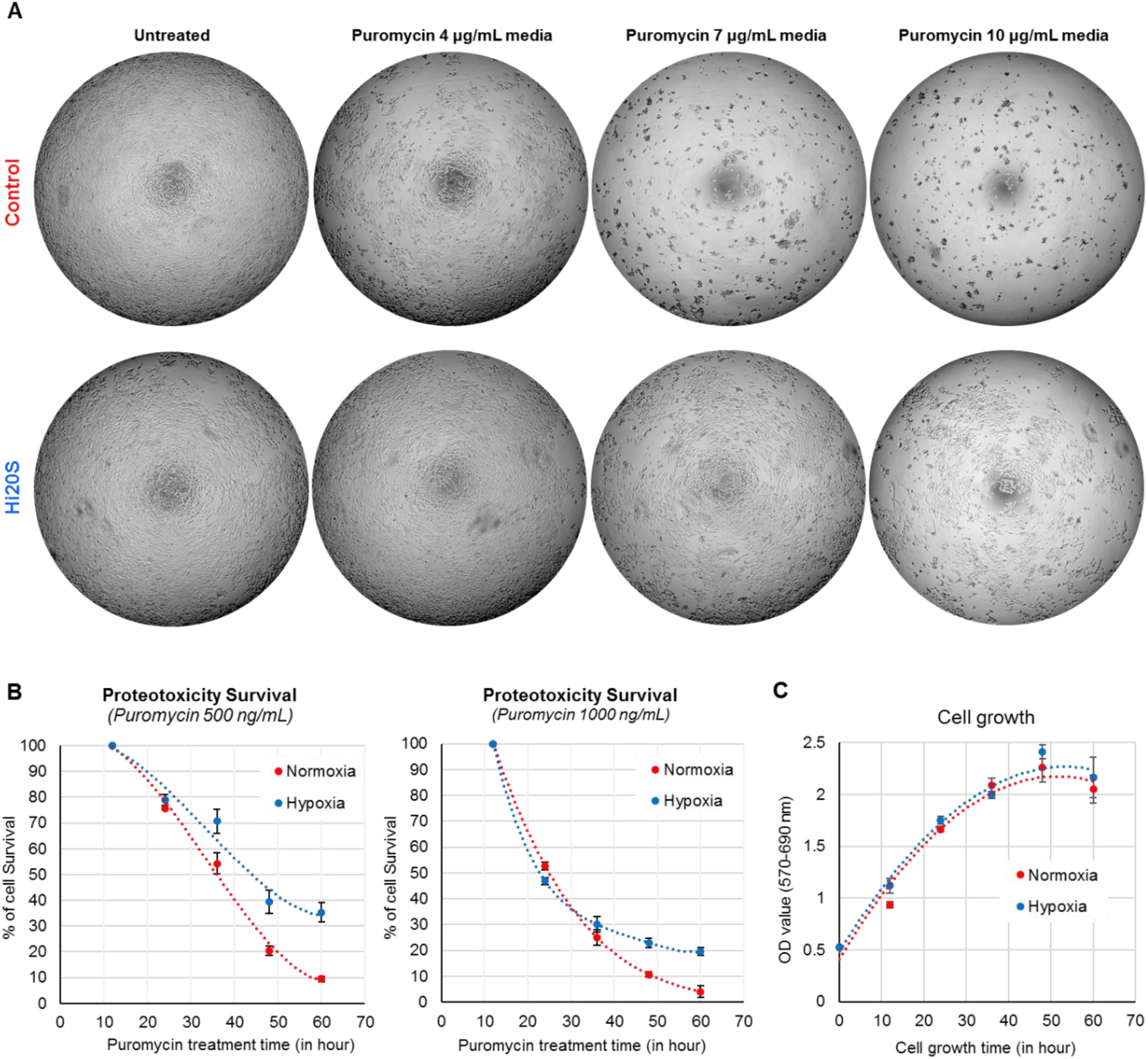
Increased levels of 20S proteasome in cells provide better survival under proteotoxic stress, Related to Figure 7. (**A**) Phase contrast images of culture plates showing survival of control and Hi20S cells after 36 h of puromycin treatment at increasing concentration. (**B**) Growth curve of HeLa cells growing under normoxia or hypoxia. Cell growth was calculated from MTT assay. Data represents the average of three experimental values (±SD). (**C**) The graphs represent cell survival upon puromycin exposure during hypoxia. HeLa cells were grown under normoxia or hypoxia (1% O_2_) with different puromycin concentration (500 ng/mL and 1000 ng/mL) for up to 60 h. Cell survival was quantified by MTT assay. Data represents the average of three experimental values (±SD).

**Table S1.** Cryo-EM data collection and refinement statistics.

|  | 20S alone | 20S+monoUb-CyclinB1-NT |  |  |
| --- | --- | --- | --- | --- |
|  |  | S0 | S1 | S2 |
| **Data collection** |  |  |  |  |
| EM equipment | Titan Krios |  | Titan Krios |  |
| Voltage (kV) | 300 |  | 300 |  |
| Detector | K2 Summit |  | K2 Summit |  |
| Pixel size (Å) | 1.318 |  | 1.318 |  |
| Electron dose (e<sup>-</sup>/Å<sup>2</sup>) | 38 |  | 38 |  |
| Exposure time (s) | 7.6 |  | 7.6 |  |
| Frames | 38 |  | 38 |  |
| Defocus range (μm) | -0.9 to -1.8 |  | -0.9 to -1.8 |  |
| **Reconstruction** |  |  |  |  |
| Software | Relion 3.0 |  | Relion 3.0 |  |
| Raw micrographs | 748 |  | 3,125 |  |
| Final particles | 154,436 | 282,834 | 72,086 | 179,590 |
| Final resolution (Å) | 3.22 | 3.38 | 4.47 | 3.88 |
| **Atomic modeling** |  |  |  |  |
| Software | Phenix & COOT |  | Rosetta, Phenix & COOT | Phenix & COOT |
| Ramachandran favored | 96.10% |  | 93.13% | 96.87% |
| Ramachandran allowed | 3.90% |  | 6.79% | 3.09% |
| Ramachandran outliers | 0.00% |  | 0.08% | 0.03% |

## REFERENCES

Adams, P.D., Afonine, P.V., Bunkoczi, G., Chen, V.B., Davis, I.W., Echols, N., Headd, J.J., Hung, L.W., Kapral, G.J., Grosse-Kunstleve, R.W., et al. (2010). PHENIX: a comprehensive Python-based system for macromolecular structure solution. Acta Crystallogr D Biol Crystallogr 66, 213–221.

Arrigo, A.P., Tanaka, K., Goldberg, A.L., and Welch, W.J. (1988). Identity of the 19S ‘prosome’ particle with the large multifunctional protease complex of mammalian cells (the proteasome). Nature 331, 192–194.

Bajorek, M., Finley, D., and Glickman, M.H. (2003). Proteasome disassembly and downregulation is correlated with viability during stationary phase. Current biology: CB 13, 1140–1144.

Baldin, V., Militello, M., Thomas, Y., Doucet, C., Fic, W., Boireau, S., Jariel-Encontre, I., Piechaczyk, M., Bertrand, E., Tazi, J., et al. (2008). A novel role for PA28gamma-proteasome in nuclear speckle organization and SR protein trafficking. Mol Biol Cell 19, 1706–1716.

Baraibar, M.A., and Friguet, B. (2012). Changes of the proteasomal system during the aging process. Prog Mol Biol Transl Sci 109, 249–275.

Bates, D., Machler, M., Bolker, B.M., and Walker, S.C. (2015). Fitting Linear Mixed-Effects Models Using lme4. J Stat Softw 67, 1–48.

Besche, H.C., Sha, Z., Kukushkin, N.V., Peth, A., Hock, E.M., Kim, W., Gygi, S., Gutierrez, J.A., Liao, H., Dick, L., et al. (2014). Autoubiquitination of the 26S Proteasome on Rpn13 Regulates Breakdown of Ubiquitin Conjugates. Embo Journal 33, 1159–1176.

Breusing, N., and Grune, T. (2008). Regulation of proteasome-mediated protein degradation during oxidative stress and aging. Biol Chem 389, 203–209.

Bulteau, A.L., Szweda, L.I., and Friguet, B. (2002). Age-dependent declines in proteasome activity in the heart. Arch Biochem Biophys 397, 298–304.

Bulteau, A.L., Szweda, L.I., and Friguet, B. (2006). Mitochondrial protein oxidation and degradation in response to oxidative stress and aging. Exp Gerontol 41, 653–657.

Cadenas, E., and Davies, K.J. (2000). Mitochondrial free radical generation, oxidative stress, and aging. Free Radic Biol Med 29, 222–230.

Chondrogianni, N., Georgila, K., Kourtis, N., Tavernarakis, N., and Gonos, E.S. (2015). 20S proteasome activation promotes life span extension and resistance to proteotoxicity in Caenorhabditis elegans. FASEB J 29, 611–622.

Cohen-Kaplan, V., Livneh, I., Avni, N., Fabre, B., Ziv, T., Kwon, Y.T., and Ciechanover, A. (2016). p62- and ubiquitin-dependent stress-induced autophagy of the mammalian 26S proteasome. Proc Natl Acad Sci U S A 113, E7490–E7499.

Cox, J., and Mann, M. (2008). MaxQuant enables high peptide identification rates, individualized p.p.b.-range mass accuracies and proteome-wide protein quantification. Nat Biotechnol 26, 1367–1372.

Dasgupta, S., Castro, L.M., Dulman, R., Yang, C., Schmidt, M., Ferro, E.S., and Fricker, L.D. (2014). Proteasome inhibitors alter levels of intracellular peptides in HEK293T and SH-SY5Y cells. PLoS One 9, e103604.

Davies, K.J. (2001). Degradation of oxidized proteins by the 20S proteasome. Biochimie 83, 301–310.

Day, S.M., Divald, A., Wang, P., Davis, F., Bartolone, S., Jones, R., and Powell, S.R. (2013). Impaired assembly and post-translational regulation of 26S proteasome in human end-stage heart failure. Circ Heart Fail 6, 544–549.

de Araujo, C.B., Heimann, A.S., Remer, R.A., Russo, L.C., Colquhoun, A., Forti, F.L., and Ferro, E.S. (2019). Intracellular Peptides in Cell Biology and Pharmacology. Biomolecules 9.

Demasi, M., and da Cunha, F.M. (2018). The physiological role of the free 20S proteasome in protein degradation: A critical review. Biochim Biophys Acta Gen Subj 1862, 2948–2954.

Deutsch, E.W., Mendoza, L., Shteynberg, D., Slagel, J., Sun, Z., and Moritz, R.L. (2015). Trans-Proteomic Pipeline, a standardized data processing pipeline for large-scale reproducible proteomics informatics. Proteomics Clin Appl 9, 745–754.

DiMaio, F., Song, Y., Li, X., Brunner, M.J., Xu, C., Conticello, V., Egelman, E., Marlovits, T., Cheng, Y., and Baker, D. (2015). Atomic-accuracy models from 4.5-A cryo-electron microscopy data with density-guided iterative local refinement. Nat Methods 12, 361–365.

Ding, Z., Xu, C., Sahu, I., Wang, Y., Fu, Z., Huang, M., Wong, C.C.L., Glickman, M.H., and Cong, Y. (2019). Structural Snapshots of 26S Proteasome Reveal Tetraubiquitin-Induced Conformations. Mol Cell 73, 1150–1161 e1156.

Emmerich, N.P.N., Nussbaum, A.K., Stevanovic, S., Priemer, M., Toes, R.E., Rammensee, H.G., and Schild, H. (2000). The human 26 S and 20 S proteasomes generate overlapping but different sets of peptide fragments from a model protein substrates. J Biol Chem 275, 21140–21148.

Emsley, P., and Cowtan, K. (2004). Coot: model-building tools for molecular graphics. Acta Crystallogr D Biol Crystallogr 60, 2126–2132.

Enenkel, C. (2018). The paradox of proteasome granules. Curr Genet 64, 137–140.

Fabre, B., Lambour, T., Garrigues, L., Ducoux-Petit, M., Amalric, F., Monsarrat, B., Burlet-Schiltz, O., and Bousquet-Dubouch, M.P. (2014). Label-free quantitative proteomics reveals the dynamics of proteasome complexes composition and stoichiometry in a wide range of human cell lines. J Proteome Res 13, 3027–3037.

Fajerman, I., Schwartz, A.L., and Ciechanover, A. (2004). Degradation of the Id2 developmental regulator: targeting via N-terminal ubiquitination. Biochemical and Biophysical Research Communications 314, 505–512.

Finley, D. (2009). Recognition and processing of ubiquitin-protein conjugates by the proteasome. Annu Rev Biochem 78, 477–513.

Fuhrmann, D.C., and Brune, B. (2017). Mitochondrial composition and function under the control of hypoxia. Redox Biol 12, 208–215.

Giordano, F.J. (2005). Oxygen, oxidative stress, hypoxia, and heart failure. J Clin Invest 115, 500–508.

Glickman, M., and Coux, O. (2001). Purification and characterization of proteasomes from Saccharomyces cerevisiae. Current protocols in protein science Chapter 21, Unit 21 25.

Glickman, M.H., and Ciechanover, A. (2002). The ubiquitin-proteasome proteolytic pathway: destruction for the sake of construction. Physiological reviews 82, 373–428.

Glickman, M.H., Rubin, D.M., Coux, O., Wefes, I., Pfeifer, G., Cjeka, Z., Baumeister, W., Fried, V.A., and Finley, D. (1998). A subcomplex of the proteasome regulatory particle required for ubiquitin-conjugate degradation and related to the COP9-signalosome and eIF3. Cell 94, 615–623.

Goddard, T.D., Huang, C.C., Meng, E.C., Pettersen, E.F., Couch, G.S., Morris, J.H., and Ferrin, T.E. (2018). UCSF ChimeraX: Meeting modern challenges in visualization and analysis. Protein Sci 27, 14–25.

Groll, M., Bajorek, M., Kohler, A., Moroder, L., Rubin, D.M., Huber, R., Glickman, M.H., and Finley, D. (2000). A gated channel into the proteasome core particle. Nature structural biology 7, 1062–1067.

Groll, M., Ditzel, L., Lowe, J., Stock, D., Bochtler, M., Bartunik, H.D., and Huber, R. (1997). Structure of 20S proteasome from yeast at 2.4 A resolution. Nature 386, 463–471.

Guterman, A., and Glickman, M.H. (2004). Complementary roles for Rpn11 and Ubp6 in deubiquitination and proteolysis by the proteasome. J Biol Chem 279, 1729–1738.

Guzy, R.D., and Schumacker, P.T. (2006). Oxygen sensing by mitochondria at complex III: the paradox of increased reactive oxygen species during hypoxia. Experimental physiology 91, 807–819.

Hanna, J., Leggett, D.S., and Finley, D. (2003). Ubiquitin depletion as a key mediator of toxicity by translational inhibitors. Mol Cell Biol 23, 9251–9261.

Hendil, K.B., Kriegenburg, F., Tanaka, K., Murata, S., Lauridsen, A.M., Johnsen, A.H., and Hartmann-Petersen, R. (2009). The 20S proteasome as an assembly platform for the 19S regulatory complex. Journal of molecular biology 394, 320–328.

Hershko, A., Ganoth, D., Sudakin, V., Dahan, A., Cohen, L.H., Luca, F.C., Ruderman, J.V., and Eytan, E. (1994). Components of a system that ligates cyclin to ubiquitin and their regulation by the protein kinase cdc2. J Biol Chem 269, 4940–4946.

Hohn, T.J., and Grune, T. (2014). The proteasome and the degradation of oxidized proteins: part III-Redox regulation of the proteasomal system. Redox Biol 2, 388–394.

Huang, C., Lewis, C., Borg, N.A., Canals, M., Diep, H., Drummond, G.R., Goode, R.J., Schittenhelm, R.B., Vinh, A., Zhu, M., et al. (2018). Proteomic Identification of Interferon-Induced Proteins with Tetratricopeptide Repeats as Markers of M1 Macrophage Polarization. J Proteome Res 17, 1485–1499.

Jung, T., and Grune, T. (2008). The proteasome and its role in the degradation of oxidized proteins. IUBMB Life 60, 743–752.

King, R.W., Peters, J.M., Tugendreich, S., Rolfe, M., Hieter, P., and Kirschner, M.W. (1995). A 20S complex containing CDC27 and CDC16 catalyzes the mitosis-specific conjugation of ubiquitin to cyclin B. Cell 81, 279–288.

Kisselev, A.F., Akopian, T.N., Woo, K.M., and Goldberg, A.L. (1999). The sizes of peptides generated from protein by mammalian 26 and 20 S proteasomes. Implications for understanding the degradative mechanism and antigen presentation. J Biol Chem 274, 3363–3371.

Kleifeld, O., Doucet, A., Prudova, A., auf dem Keller, U., Gioia, M., Kizhakkedathu, J.N., and Overall, C.M. (2011). Identifying and quantifying proteolytic events and the natural N terminome by terminal amine isotopic labeling of substrates. Nat Protoc 6, 1578–1611.

Kumar Deshmukh, F., Yaffe, D., Olshina, M.A., Ben-Nissan, G., and Sharon, M. (2019). The Contribution of the 20S Proteasome to Proteostasis. Biomolecules 9.

Leggett, D.S., Glickman, M.H., and Finley, D. (2005). Purification of proteasomes, proteasome subcomplexes, and proteasome-associated proteins from budding yeast. Methods Mol Biol 301, 57–70.

Livnat-Levanon, N., Kevei, E., Kleifeld, O., Krutauz, D., Segref, A., Rinaldi, T., Erpapazoglou, Z., Cohen, M., Reis, N., Hoppe, T., et al. (2014). Reversible 26S proteasome disassembly upon mitochondrial stress. Cell reports 7, 1371–1380.

Ludtke, S.J., Baldwin, P.R., and Chiu, W. (1999). EMAN: semiautomated software for high-resolution single-particle reconstructions. J Struct Biol 128, 82–97.

Mansour, W., Nakasone, M.A., von Delbruck, M., Yu, Z., Krutauz, D., Reis, N., Kleifeld, O., Sommer, T., Fushman, D., and Glickman, M.H. (2015). Disassembly of Lys11 and mixed linkage polyubiquitin conjugates provides insights into function of proteasomal deubiquitinases Rpn11 and Ubp6. J Biol Chem 290, 4688–4704.

Marshall, R.S., Li, F., Gemperline, D.C., Book, A.J., and Vierstra, R.D. (2015). Autophagic Degradation of the 26S Proteasome Is Mediated by the Dual ATG8/Ubiquitin Receptor RPN10 in Arabidopsis. Mol Cell 58, 1053–1066.

Marshall, R.S., and Vierstra, R.D. (2018). Proteasome storage granules protect proteasomes from autophagic degradation upon carbon starvation. Elife 7.

Mastronarde, D.N. (2005). Automated electron microscope tomography using robust prediction of specimen movements. J Struct Biol 152, 36–51.

Mayor, T., Sharon, M., and Glickman, M.H. (2016). Tuning the proteasome to brighten the end of the journey. American journal of physiology Cell physiology 311, C793–C804.

Meszaros, B., Erdos, G., and Dosztanyi, Z. (2018). IUPred2A: context-dependent prediction of protein disorder as a function of redox state and protein binding. Nucleic Acids Res 46, W329–W337.

Myers, N., Olender, T., Savidor, A., Levin, Y., Reuven, N., and Shaul, Y. (2018). The Disordered Landscape of the 20S Proteasome Substrates Reveals Tight Association with Phase Separated Granules. Proteomics 18, e1800076.

Nettling, M., Treutler, H., Grau, J., Keilwagen, J., Posch, S., and Grosse, I. (2015). DiffLogo: a comparative visualization of sequence motifs. nBMC Bioinformatics 16, 387.

Njomen, E., Osmulski, P.A., Jones, C.L., Gaczynska, M., and Tepe, J.J. (2018). Small Molecule Modulation of Proteasome Assembly. Biochemistry 57, 4214–4224.

Okonko, D.O., and Shah, A.M. (2015). Heart failure: mitochondrial dysfunction and oxidative stress in CHF. Nature reviews Cardiology 12, 6–8.

Orlowski, M., and Wilk, S. (2003). Ubiquitin-independent proteolytic functions of the proteasome. Arch Biochem Biophys 415, 1–5.

Osmulski, P.A., and Gaczynska, M. (2002). Nanoenzymology of the 20S proteasome: proteasomal actions are controlled by the allosteric transition. Biochemistry 41, 7047–7053.

Osmulski, P.A., Hochstrasser, M., and Gaczynska, M. (2009). A tetrahedral transition state at the active sites of the 20S proteasome is coupled to opening of the alpha-ring channel. Structure 17, 1137–1147.

Peters, L.Z., Hazan, R., Breker, M., Schuldiner, M., and Ben-Aroya, S. (2013). Formation and dissociation of proteasome storage granules are regulated by cytosolic pH. J Cell Biol 201, 663–671.

Pettersen, E.F., Goddard, T.D., Huang, C.C., Couch, G.S., Greenblatt, D.M., Meng, E.C., and Ferrin, T.E. (2004). UCSF Chimera--a visualization system for exploratory research and analysis. J Comput Chem 25, 1605–1612.

Pickering, A.M., and Davies, K.J. (2012). Degradation of damaged proteins: the main function of the 20S proteasome. Prog Mol Biol Transl Sci 109, 227–248.

Pino, L.K., Searle, B.C., Bollinger, J.G., Nunn, B., MacLean, B., and MacCoss, M.J. (2017). The Skyline ecosystem: Informatics for quantitative mass spectrometry proteomics. Mass Spectrom Rev.

Predmore, J.M., Wang, P., Davis, F., Bartolone, S., Westfall, M.V., Dyke, D.B., Pagani, F., Powell, S.R., and Day, S.M. (2010). Ubiquitin proteasome dysfunction in human hypertrophic and dilated cardiomyopathies. Circulation 121, 997–1004.

Rabl, J., Smith, D.M., Yu, Y., Chang, S.C., Goldberg, A.L., and Cheng, Y. (2008). Mechanism of gate opening in the 20S proteasome by the proteasomal ATPases. Mol Cell 30, 360–368.

Raynes, R., Pomatto, L.C., and Davies, K.J. (2016). Degradation of oxidized proteins by the proteasome: Distinguishing between the 20S, 26S, and immunoproteasome proteolytic pathways. Mol Aspects Med 50, 41–55.

Reinheckel, T., Sitte, N., Ullrich, O., Kuckelkorn, U., Davies, K.J., and Grune, T. (1998). Comparative resistance of the 20S and 26S proteasome to oxidative stress. Biochem J 335 (Pt 3), 637–642.

Robinson, M.D., McCarthy, D.J., and Smyth, G.K. (2010). edgeR: a Bioconductor package for differential expression analysis of digital gene expression data. Bioinformatics 26, 139–140.

Rohou, A., and Grigorieff, N. (2015). CTFFIND4: Fast and accurate defocus estimation from electron micrographs. J Struct Biol 192, 216–221.

Saeki, Y. (2017). Ubiquitin recognition by the proteasome. J Biochem 161, 113–124.

Saez, I., and Vilchez, D. (2014). The Mechanistic Links Between Proteasome Activity, Aging and Age-related Diseases. Current genomics 15, 38–51.

Sahara, K., Kogleck, L., Yashiroda, H., and Murata, S. (2014). The mechanism for molecular assembly of the proteasome. Adv Biol Regul 54, 51–58.

Shringarpure, R., Grune, T., and Davies, K.J. (2001). Protein oxidation and 20S proteasome-dependent proteolysis in mammalian cells. Cell Mol Life Sci 58, 1442–1450.

Shringarpure, R., Grune, T., Mehlhase, J., and Davies, K.J. (2003). Ubiquitin conjugation is not required for the degradation of oxidized proteins by proteasome. J Biol Chem 278, 311–318.

Singh, S.K., Sahu, I., Mali, S.M., Hemantha, H.P., Kleifeld, O., Glickman, M.H., and Brik, A. (2016). Synthetic Uncleavable Ubiquitinated Proteins Dissect Proteasome Deubiquitination and Degradation, and Highlight Distinctive Fate of Tetraubiquitin. J Am Chem Soc 138, 16004–16015.

Solaini, G., Baracca, A., Lenaz, G., and Sgarbi, G. (2010). Hypoxia and mitochondrial oxidative metabolism. Biochim Biophys Acta 1797, 1171–1177.

Spitzer, M., Wildenhain, J., Rappsilber, J., and Tyers, M. (2014). BoxPlotR: a web tool for generation of box plots. Nat Methods 11, 121–122.

Sun, H., and Brik, A. (2019). The Journey for the Total Chemical Synthesis of a 53 kDa Protein. Acc Chem Res.

Suskiewicz, M.J., Sussman, J.L., Silman, I., and Shaul, Y. (2011). Context-dependent resistance to proteolysis of intrinsically disordered proteins. Protein Sci 20, 1285–1297.

Suzuki, R., Moriishi, K., Fukuda, K., Shirakura, M., Ishii, K., Shoji, I., Wakita, T., Miyamura, T., Matsuura, Y., and Suzuki, T. (2009). Proteasomal turnover of hepatitis C virus core protein is regulated by two distinct mechanisms: a ubiquitin-dependent mechanism and a ubiquitin-independent but PA28gamma-dependent mechanism. J Virol 83, 2389–2392.

Tait, S.W., de Vries, E., Maas, C., Keller, A.M., D’Santos, C.S., and Borst, J. (2007). Apoptosis induction by Bid requires unconventional ubiquitination and degradation of its N-terminal fragment. J Cell Biol 179, 1453–1466.

Tomko, R.J., Jr., and Hochstrasser, M. (2013). Molecular architecture and assembly of the eukaryotic proteasome. Annu Rev Biochem 82, 415–445.

Tonoki, A., Kuranaga, E., Tomioka, T., Hamazaki, J., Murata, S., Tanaka, K., and Miura, M. (2009). Genetic evidence linking age-dependent attenuation of the 26S proteasome with the aging process. Mol Cell Biol 29, 1095–1106.

Toste Rego, A., and da Fonseca, P.C.A. (2019). Characterization of Fully Recombinant Human 20S and 20S-PA200 Proteasome Complexes. Mol Cell 76, 138–147 e135.

Tsvetkov, P., Mendillo, M.L., Zhao, J., Carette, J.E., Merrill, P.H., Cikes, D., Varadarajan, M., van Diemen, F.R., Penninger, J.M., Goldberg, A.L., et al. (2015). Compromising the 19S proteasome complex protects cells from reduced flux through the proteasome. Elife 4.

Tsvetkov, P., Sokol, E., Jin, D., Brune, Z., Thiru, P., Ghandi, M., Garraway, L.A., Gupta, P.B., Santagata, S., Whitesell, L., et al. (2017). Suppression of 19S proteasome subunits marks emergence of an altered cell state in diverse cancers. Proc Natl Acad Sci U S A 114, 382–387.

Unno, M., Mizushima, T., Morimoto, Y., Tomisugi, Y., Tanaka, K., Yasuoka, N., and Tsukihara, T. (2002). The structure of the mammalian 20S proteasome at 2.75 A resolution. Structure 10, 609–618.

Wang, X., Yen, J., Kaiser, P., and Huang, L. (2010). Regulation of the 26S proteasome complex during oxidative stress. Sci Signal 3, ra88.

Wen, X., and Klionsky, D.J. (2016). The proteasome subunit RPN10 functions as a specific receptor for degradation of the 26S proteasome by macroautophagy in Arabidopsis. Autophagy 12, 905–906.

Whitby, F.J., Masters, E.I., Kramer, L., Knowlton, J.R., Yao, Y., Wang, C., and Hill, C.P. (2000). Structural basis for the activation of 20S proteasomes by 11S regulators. Nature 408, 115–120.

Yamano, H., Tsurumi, C., Gannon, J., and Hunt, T. (1998). The role of the destruction box and its neighbouring lysine residues in cyclin B for anaphase ubiquitin-dependent proteolysis in fission yeast: defining the D-box receptor. EMBO J 17, 5670–5678.

Yau, R.G., Doerner, K., Castellanos, E.R., Haakonsen, D.L., Werner, A., Wang, N., Yang, X.W., Martinez-Martin, N., Matsumoto, M.L., Dixit, V.M., et al. (2017). Assembly and Function of Heterotypic Ubiquitin Chains in Cell-Cycle and Protein Quality Control. Cell 171, 918–933 e920.

Yu, H., and Matouschek, A. (2017). Recognition of Client Proteins by the Proteasome. Annual review of biophysics.

Zhang, K. (2016). Gctf: Real-time CTF determination and correction. J Struct Biol 193, 1–12.

Zheng, S.Q., Palovcak, E., Armache, J.P., Verba, K.A., Cheng, Y., and Agard, D.A. (2017). MotionCor2: anisotropic correction of beam-induced motion for improved cryo-electron microscopy. Nat Methods 14, 331–332.

Zhong, Q., Gao, W., Du, F., and Wang, X. (2005). Mule/ARF-BP1, a BH3-only E3 ubiquitin ligase, catalyzes the polyubiquitination of Mcl-1 and regulates apoptosis. Cell 121, 1085–1095.

